# Identifying loci with different allele frequencies among cases of eight psychiatric disorders using CC-GWAS

**DOI:** 10.1101/2020.03.04.977389

**Authors:** Wouter J. Peyrot, Alkes L. Price

## Abstract

Psychiatric disorders are highly genetically correlated, and many studies have focused on their shared genetic components. However, little research has been conducted on the genetic differences between psychiatric disorders, because case-case comparisons of allele frequencies among cases currently require individual-level data from cases of both disorders. We developed a new method (CC-GWAS) to test for differences in allele frequency among cases of two different disorders using summary statistics from the respective case-control GWAS; CC-GWAS relies on analytical assessments of the genetic distance between cases and controls of each disorder. Simulations and analytical computations confirm that CC-GWAS is well-powered and attains effective control of type I error. In particular, CC-GWAS identifies and discards false positive associations that can arise due to differential tagging of a shared causal SNP (with the same allele frequency in cases of both disorders), e.g. due to subtle differences in ancestry between the input case-control studies. We applied CC-GWAS to publicly available summary statistics for schizophrenia, bipolar disorder and major depressive disorder, and identified 116 independent genome-wide significant loci distinguishing these three disorders, including 21 CC-GWAS-specific loci that were not genome-wide significant in the input case-control summary statistics. Two of the CC-GWAS-specific loci implicate the genes *KLF6* and *KLF16* from the Kruppel-like family of transcription factors; these genes have been linked to neurite outgrowth and axon regeneration. We performed a broader set of case-case comparisons by additionally analyzing ADHD, anorexia nervosa, autism, obsessive-compulsive disorder and Tourette’s Syndrome, yielding a total of 196 independent loci distinguishing eight psychiatric disorders, including 72 CC-GWAS-specific loci. We confirmed that loci identified by CC-GWAS replicated convincingly in applications to data sets for which independent replication data were available. In conclusion, CC-GWAS robustly identifies loci with different allele frequencies among cases of different disorders using results from the respective case-control GWAS, providing new insights into the genetic differences between eight psychiatric disorders.

## Introduction

Psychiatric disorders are highly genetically correlated, and many studies have focused on their shared genetic components, including genetic correlation estimates of up to 0.7^1–3^ and recent identification of 109 pleotropic loci across a broad set of eight psychiatric disorders^3^. However, much less research has been conducted on the genetic differences between psychiatric disorders, and biological differences between psychiatric disorders are poorly understood. Currently, differential diagnosis between disorders is often challenging and treatments are often non-disorder-specific, highlighting the importance of studying genetic differences between psychiatric disorders.

A recent study^4^ progressed the research on genetic differences between disorders by comparing individual-level data of 24k SCZ cases vs. 15k BIP cases, yielding two significantly associated loci. However, ∼25% of the cases were discarded compared to the respective case-control data (owing to non-matching ancestry and genotyping platform). Methods that analyse case-control summary statistics may be advantageous, because they make use of all genotyped samples and because summary statistics are often broadly publicly available^5^. Indeed, several methods have been developed to analyse GWAS summary statistics of two complex traits^3,6–13^, but none of these methods can be used to conduct a case-case comparison (see Discussion). Currently, case-case comparisons of allele frequencies among cases of two disorders require individual-level data from cases of both disorders.

In this study, we propose a new method (CC-GWAS) to compare cases of two disorders based on the respective case-control GWAS summary statistics. CC-GWAS relies on a new genetic distance measure (*F*_*ST,causal*_) quantifying the genetic distances between cases and controls of different disorders. We first apply CC-GWAS to publicly available GWAS summary statistics of the mood and psychotic disorders^3^, schizophrenia (SCZ)^14,15^, bipolar disorder (BIP)^16^ and major depressive disorder (MDD)^17^. Subsequently, we analyse all comparisons of eight psychiatric disorders by additionally analysing attention deficit/hyperactivity disorder (ADHD)^18^, anorexia nervosa (AN)^19^, autism spectrum disorder (ASD)^20^, obsessive–compulsive disorder (OCD)^21^, Tourette’s Syndrome and Other Tic Disorders (TS)^22^. Finally, we replicate CC-GWAS results of SCZ vs. MDD based on subsets of the data for which independent replication data were available.

## Results

### Overview of methods

CC-GWAS detects differences in allele frequencies among cases of two different disorders A and B by analysing case-control GWAS summary statistics for each disorder. CC-GWAS relies on the analytical variances and covariances of genetic effects of causal variants distinguishing caseA vs. controlA (A1A0), caseB vs. controlB (B1B0), and caseA vs. caseB (A1B1); expectations of these variances and covariances can be derived based on estimates of the SNP-based heritabilities (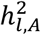 and 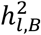), lifetime population prevalences (*K*_*A*_ and *K*_*B*_), genetic correlation (*r*_*g*_), and number of independent causal variants (*m*).

CC-GWAS weights the effect sizes from the respective case-control GWAS using weights that minimize the expected squared difference between estimated and true A1B1 effect sizes; we refer to these as CC-GWAS ordinary least squares (CC-GWAS_OLS_) weights (see Methods). The CC-GWAS_OLS_ weights are designed to optimize power to detect A1B1, and depend on sample size, sample overlap, and the expected variances and covariances of effect sizes. The CC-GWAS_OLS_ weights may be susceptible to type I error for SNPs with nonzero A1A0 and B1B0 effect sizes but zero A1B1 effect size, which we refer to as “stress test” SNPs. To mitigate this, CC-GWAS also computes sample size independent weights based on infinite sample size; we refer to these as CC-GWAS Exact (CC-GWAS_Exact_) weights (see Methods). (At very large sample sizes, the CC-GWAS_OLS_ weights converge to the CC-GWAS_Exact_ weights.) CC-GWAS reports a SNP as statistically significant if it achieves *P*<5 ×10^−8^ using CC-GWAS_OLS_ weights *and P*<10^−4^ using CC-GWAS_Exact_ weights, balancing power and type I error (see Simulations). Specifically, using the CC-GWAS_OLS_ weights optimizes power and protect against type I error at null-null SNPs (with no impact on either disorder), while using the CC-GWAS_Exact_ weights protects against type I error at stress test SNPs. For both the CC-GWAS_OLS_ test and the CC-GWAS_Exact_ test, p-values are obtained by dividing the estimated betas by the respective standard errors, and assuming that the resulting z-score follows a standard normal distribution. For statistically significant SNPs, CC-GWAS outputs CC-GWAS_OLS_ effect sizes reflecting direction and magnitude of effect. We note that the CC-GWAS_OLS_ weights assume that A1A0 and B1B0 effect sizes for causal variants follow a bivariate normal distribution. Violation of this assumption may result in increased type I error when using the CC-GWAS_OLS_ weights only; as noted above, the CC-GWAS_Exact_ weights protect against type I error in such scenarios (CC-GWAS can thus be applied to pairs of disorders with any bivariate genetic architecture). Importantly, CC-GWAS identifies and discards false positive associations that can arise due to differential tagging of a causal stress test SNP (with the same allele frequency in cases of both disorders), e.g. due to subtle differences in ancestry between the input case-control studies. Specifically, CC-GWAS screens the region around every genome-wide significant candidate CC-GWAS variant for evidence of a differentially linked stress test SNP, and conservatively filters the candidate CC-GWAS variant when suggestive evidence of a differentially linked stress test SNP is detected (see Methods). When there is substantial uncertainty in population prevalence, a range of possible disorder prevalences can be specified for the CC-GWAS_Exact_ component of CC-GWAS to maximally prevent inflated type I error at stress test SNPs (while specifying the most likely disorder prevalence for the CC-GWAS_OLS_ component, as the specified disorder prevalence does not impact type I error at null-null SNPs; see *Main simulations*). We further note that sample overlap of controls increases the power of CC-GWAS, by providing a more direct comparison of caseA vs. caseB. Further details of the CC-GWAS method are provided in the Methods section; we have released open-source software implementing the method (see URLs).

The CC-GWAS_OLS_ weights depend on a population-level quantity that we refer to as *F*_*ST,causal*_, the average normalized squared difference in allele frequencies of causal variants. *F*_*ST,causal*_ is derived based on the SNP-based heritabilities (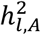 and 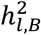), lifetime population prevalences (*K*_*A*_ and *K*_*B*_), genetic correlation (*r*_*g*_), and number of independent causal variants (*m*) (see Methods for a detailed discussion of the assumed number of causal SNPs in applications of CC-GWAS). *F*_*ST,causal*_ allows for a direct comparison of cases and controls using 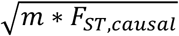 as a genetic distance measure, where the square root facilitates 2-dimensional visualization (see Figure 1 and Methods). We note that the distances can be intuitively interpreted as (i) the square root of the average squared difference in allele frequency at causal SNPs, (ii) proportional to the average power in GWAS (assuming equal sample sizes and numbers of causal SNPs), (iii) heritability on the observed scale based on 50/50 ascertainment (although the heritability has no clear interpretation when comparing overlapping small subsets of the population), and (iv) an indication of the accuracy of polygenic risk prediction.

**Figure 1.**
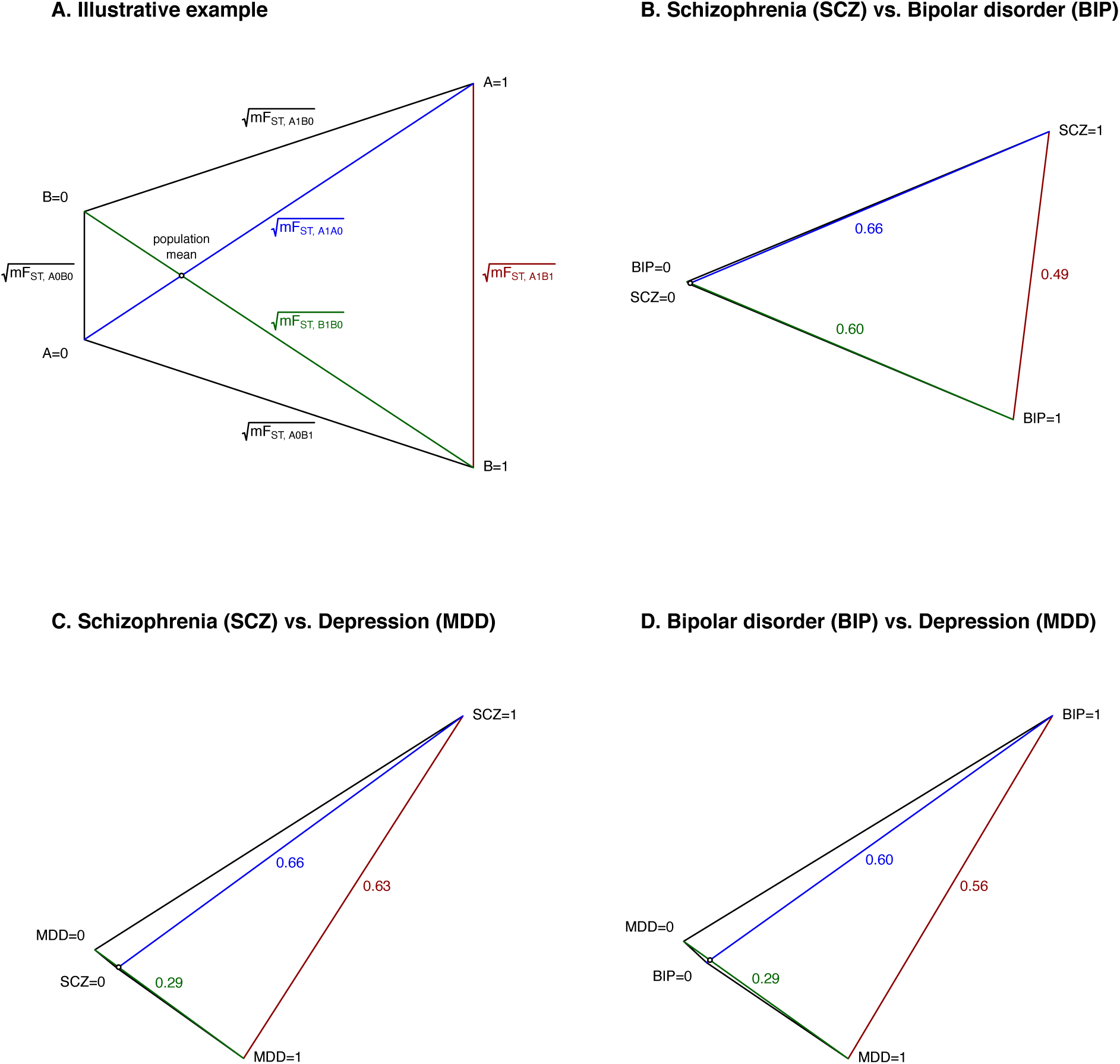
Genetic distance between cases and/or controls of SCZ, BIP and MDD. We report genetic distances for (A) an illustrative example, (B) SCZ vs. BIP, (C) SCZ vs. MDD and (D) SCZ vs. BIP. Genetic distances are displayed as 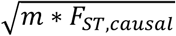, where *m* is the number of independent causal variants and the square root facilitates 2-dimensional visualization. The quantity *m* * *F*_*ST,causal*,_ is derived based on the respective population prevalences, SNP-based heritabilities and genetic correlations (reported in Table 1). Approximate standard errors of *m* * *F*_*ST,causal,A1B1*_ are 0.04 for SCZ vs. BIP, 0.02 for SCZ vs. MDD and 0.03 for BIP vs. MDD (see Methods). The cosine of the angle between the lines A1-A0 and B1-B0 is equal to the genetic correlation between disorder A and disorder B (see Methods). For SCZ and BIP, despite the large genetic correlation (*r*_*g*_ = 0.7), the genetic distance between SCZ cases and BIP cases is only slightly smaller 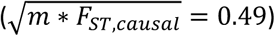 than the case-control distances for SCZ (0.66) and BIP (0.60), because of the doubly strong ascertainment (due to low disorder prevalences) in SCZ cases and BIP cases and because a genetic correlation of 0.7 is still considerably smaller than a genetic correlation of e.g. 0.9 (Figure S20). For SCZ and MDD (*r*_*g*_ = 0.31), the genetic distance between MDD cases and SCZ cases (0.63) is larger than for MDD case-control (0.29) (Panel C) owing to the larger prevalence and lower heritability of MDD, consistent with our empirical findings (see below). For MDD and BIP (*r*_*g*_ = 0.33), genetic distances are similar to MDD and SCZ (Panel D). Numerical results are reported in Table S11.

### Main simulations

We assessed the power and type I error of CC-GWAS using both simulations with individual-level data and analytical computations (see Methods). We compared four methods: CC-GWAS; the CC-GWAS_OLS_ component of CC-GWAS; the CC-GWAS_Exact_ component of CC-GWAS; and a naïve method that uses weight +1 for A1A0 and −1 for B1B0 (Delta method). All four methods are unpublished; we are not currently aware of any published method for performing case-case GWAS using case-control summary statistics (see Discussion). We assessed (i) power to detect causal SNPs with case-control effect sizes for both disorders drawn from a bivariate normal distribution (allele frequencies A0≠A1, B0≠B1, A1≠B1); (ii) type I error for “null-null” SNPs, defined as SNPs with no effect on either disorder (A0=A1, B0=B1, A1=B1); and (iii) type I error for “stress test” SNPs, defined as SNPs with A0≠A1, B0≠B1, A1=B1 (see above). Default parameter settings were loosely based on the genetic architectures of SCZ and MDD with liability-scale *h*^2^=0.2, prevalence *K*=0.01, and sample size 100,000 cases + 100,000 controls for disorder A; liability-scale *h*^2^=0.1, prevalence *K*=0.15, and sample size 100,000 cases + 100,000 controls for disorder B; genetic correlation *r*_*g*_=0.5 between disorders; and *m*=5,000 causal SNPs affecting both disorders with causal effect sizes following a bivariate normal distribution. For these parameter settings, the weights are 0.86 for A1A0 and –0.55 for B1B0 for the CC-GWAS_OLS_ component, and 0.99 and –0.85 respectively for the CC-GWAS_Exact_ component. The CC-GWAS_OLS_ component assigns relatively more weight to A1A0 (0.86/0.55=1.56) than the CC-GWAS_Exact_ component (0.99/0.85=1.16), because of the larger heritability and lower prevalence of A1A0 (implying higher signal to noise ratio at the same case-control sample size^23,24^). The CC-GWAS_OLS_ weights are shrunk in comparison to the CC-GWAS_Exact_ weights, accounting for the imperfect signal to noise ratio at finite sample size (however, CC-GWAS *p*-values are insensitive to rescaling the weights at a fixed ratio). Each stress test SNP was specified to explain 0.10% of liability-scale variance in A and 0.29% of liability-scale variance in B (resulting in allele frequency A1=B1); we focused on large-effect stress test SNPs to provide a maximally stringent assessment of the robustness of CC-GWAS to stress test SNPs. Other parameter settings were also explored.

**Table 1.** Summary of CC-GWAS results for schizophrenia, bipolar disorder and major depressive disorder. For each pair of schizophrenia (SCZ)^15^, bipolar disorder (BIP)^16^ and major depressive disorder (MDD)^17^, we report the case-control sample sizes, #SNPs, the most likely prevalence (*K*)^84^, liability-scale heritability estimated using stratified LD score regression^37–39^ (*h*^2^), genetic correlation estimated using cross-trait LD score regression^2^ (*r*_*g*_), CC-GWAS_OLS_ weights (based on the most likely prevalences), number of independent genome-wide significant loci for each case-control comparison, number of independent genome-significant CC-GWAS loci, and number of independent genome-significant CC-GWAS loci that are CC-GWAS-specific. CC-GWAS_Exact_ weights are equal to (1 − *K*_*A*_) for disorder A and − (1 − *K*_*B*_) for disorder B. We specified a range of prevalences to the CC-GWAS_Exact_ component for SCZ (0.4%-1.0%)^15,84^, BIP (0.5%-2.0%)^16^, and MDD (16%-30%)^17,84^ (yielding 2 × 2 = 4 CC-GWAS_Exact_ p-values per comparison, all required to be < 10^−4^).

|  |  |  |  |  |  |  |  |  |  | Number of significant independent loci |  |  |  |
| --- | --- | --- | --- | --- | --- | --- | --- | --- | --- | --- | --- | --- | --- |
|  |  |  |  |  |  |  |  |  |  | CC-GWAS |  |  |  |
|  |  |  |  |  |  |  |  |  |  | CC-GWAS |  |  |  |
|  |  |  |  |  |  |  |  |  |  | CC-GWAS |  |  |  |
| A1A0 (N case/ N control) | B1B0 (N case/ N control) | # SNPs | A1A0 | | B1B0 | | $r_g$ | OLS weights | | A1A0 | B1B0 | all | CC-GWAS specific |
| | | | $K$ (%) | $h^2$ | $K$ (%) | $h^2$ | | | | | | | |
| SCZ (40,675/64,643) | BIP (20,352/31,358) | 4,548,414 | 0.40 | 0.20 | 1.00 | 0.20 | 0.70 | 0.55/-0.43 |  | 139 | 15 | **12** | **7** |
| SCZ (40,675/64,643) | MDD (170,756/329,443) | 4,483,387 | 0.40 | 0.20 | 16.00 | 0.10 | 0.31 | 0.77/-0.51 |  | 139 | 50 | **99** | **10** |
| BIP (20,352/31,358) | MDD (170,756/329,443) | 6,265,453 | 1.00 | 0.20 | 16.00 | 0.10 | 0.33 | 0.58/-0.43 |  | 14 | 53 | **10** | **4** |

Results of analytical computations are reported in Figure 2 and Table S1; simulations with individual-level data produced identical results (Table S2), thus we focus primarily on results of analytical computations. We reached three main conclusions. First, CC-GWAS attains similar power as the CC-GWAS_OLS_ component by itself, higher power than the CC-GWAS_Exact_ component by itself, and much higher power than the Delta method (Figure 2A); we note that this is a best-case scenario for CC-GWAS, as the simulated bivariate genetic architecture follows the CC-GWAS assumptions. As expected, power increases with increasing sample size and decreases with increasing genetic correlation. The power of CC-GWAS to detect case-case differences lies in between the power of the input A1A0 and B1B0 summary statistics to detect case-control differences (Figure S1). Second, all methods perfectly control type I error at null-null SNPs, with a per-SNP type I error rate < 5 ×10^−8^ (Figure 2B). Third, although the CC-GWAS_OLS_ component has a severe type I error problem at stress test SNPs (particularly when the genetic correlation is large), CC-GWAS (incorporating the CC-GWAS_Exact_ component) attains effective control of type I error at stress test SNPs (per-SNP type I error rate < 10^−4^; Figure 2C), an extreme category of SNPs that is not likely to occur often in empirical data. Indeed, a per-study type I error rate < 0.05 at stress test SNPs (which is the standard in GWAS using 5×10^−8^ as the significance threshold^25^), would allow for the extreme scenario of 500 (500 * 10^−4^ = 0.05) large-effect stress test SNPs (we note that 100 stress test SNPs in our simulations already explain 29% of liability variance in disorder B, and that the per-SNP type I error rate is much smaller for small-effect stress test SNPs, see below). Notably, with increasing sample size the CC-GWAS_OLS_ weights converge towards the CC-GWAS_Exact_ weights (Figure S2), resulting in decreasing type I error rate for stress test SNPs. In conclusion, CC-GWAS balances the high power of the CC-GWAS_OLS_ component with effective control of type I error of the CC-GWAS_Exact_ component.

**Figure 2.**
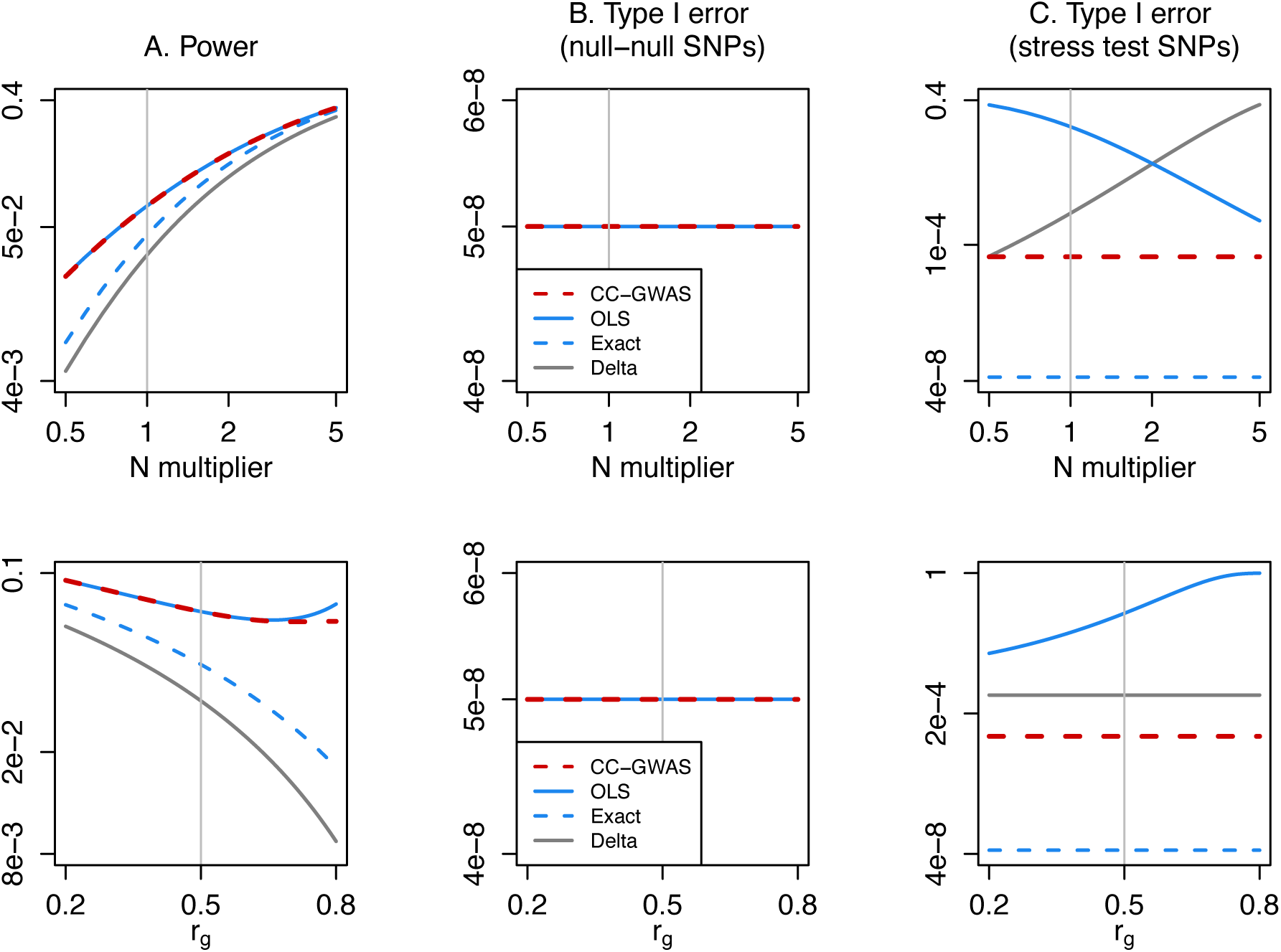
Power and type I error of CC-GWAS. We report (A) the power to detect SNPs with effect sizes following a bivariate normal distribution, (B) the type I error rate for loci with no effect on A1A0 or B1B0 (“null-null” SNPs) and (C) the type I error rate for SNPs with the same allele frequency in A1 vs. B1 that explain 0.10% of variance in A1 vs. A0 and 0.29% of variance in B1 vs. B0 (“stress test” SNPs), for each of four methods: CC-GWAS, the CC-GWAS_OLS_ component, the CC-GWAS_Exact_ component, and a naïve Delta method (see text). Default parameter settings are: *h*^2^=0.2, prevalence *K*=0.01, and sample size 100,000 cases + 100,000 controls for disorder A; liability-scale *h*^2^=0.1, prevalence *K*=0.15, and sample size 100,000 cases + 100,000 controls for disorder B; *m*=5,000 causal SNPs for each disorder; and genetic correlation *r*_*g*_=0.5 between disorders. Numerical results of these analytical computations are reported in Table S1, and confirmed with simulation results in Table S2.

We performed 12 secondary analyses, yielding the following conclusions. First, results were similar when varying *K*_*B*_, 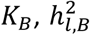 and *m* (Figure S3); in particular, results were similar when increasing from *m*=5,000 causal SNPs to *m*=10,000 causal SNPs (Figure S4; we used *m*=5,000 causal SNPs in our main assessment because this setting corresponds to higher absolute power). Second, sample overlap in controls increases power, as this provides a more direct case-case comparisons (Figure S5). Third, the type I error rate of CC-GWAS is slightly less 1 in 10,000 for stress test SNPs explaining a large proportion of case-control variance (0.29% for disorder B in Figure 2C), but is much smaller for stress test SNPs explaining less variance (Figure S6). Fourth, when employing a more stringent *p*-value threshold for the CC-GWAS_Exact_ component of CC-GWAS than the default threshold of 10^−4^, both the power for causal SNPs and the type I error rate for stress test SNPs decrease (Figure S7). We believe that the default threshold of 10^−4^ provides sufficient protection against type I error of stress test SNPs, which cannot be numerous (e.g. 100 independent stress test SNPs as defined in Figure 2C would explain 29% of liability-scale variance in disorder B). Fifth, SNPs with small MAF of 0.01 and large odds ratios yielded no increase in type I error in auxiliary simulations (Table S2D). Sixth, we assessed CC-GWAS using the type S error rate, defined as the proportion of significantly identified loci (true positives) identified with the wrong sign^26,27^; this is an appealing metric for CC-GWAS, whose complexity precludes metrics based on distributions of test statistics. We determined that the type S error rate was negligible: less than 1 in 1,000,000 significantly identified variants for all parameter settings of Figure 2A (Table S3). Seventh, a direct case-case GWAS (which requires individual-level data) may be more powerful than CC-GWAS when a great majority of cases can be included (Figure S8), although results may contain false positive associations due to differential tagging of a causal stress test SNP (see *Assessing the robustness of CC-GWAS*). Eighth, when case-case GWAS results are available, a method incorporating these results can be applied to further increase power (CC-GWAS+; Figure S9); CC-GWAS+ is related to the MTAG^6^ method, as the CC-GWAS_OLS+_ weights are roughly proportional to the MTAG weights, however, an important advantage of CC-GWAS+ is that it adequately controls type I error at stress test SNPs, while MTAG does not (Supplementary Note; Table S4). Ninth, if disorder prevalence(s) are misspecified, the power of CC-GWAS will either decrease or increase, the type I error rate at stress test SNPs will respectively decrease or increase, and the type I error at null-null SNPs will not change (Figure S10); as noted above, CC-GWAS allows for specifying a range of possible disorder prevalences in CC-GWAS_Exact_ to prevent type I error at stress test SNPs. (The CC-GWAS_OLS_ component protects against type I error at null-null SNPs, which is not impacted by misspecification of the disorder prevalence. Hence, the most likely disorder prevalence is specified for the CC-GWAS_OLS_ component). Tenth, misspecifying the heritabilities and genetic correlation had little impact on power, no impact on type I error at null-null SNPs, and no impact on type I error at stress test SNPs (Figure S11). Eleventh, misspecifying of the number of causal SNPs *m* had a modest impact on power, no impact on type I error at null-null SNPs, and no impact on type I error at stress test SNPs (Figure S12). Twelfth, application of CC-GWAS to two simulated disorders with different sets of causal variants (violating the CC-GWAS assumption that effect sizes for causal variants follow a bivariate normal distribution; see Overview of methods) attained similar power and type I error as application of CC-GWAS to two disorders with the same set of causal variants (Table S5).

### Assessing the robustness of CC-GWAS

We performed two sets of analyses to further assess the robustness of CC-GWAS: we performed simulations to assess the robustness of CC-GWAS to false positive associations due to differential tagging of a causal stress test SNP (i.e. a shared causal SNP with the same allele frequency in cases of both disorders), and we applied CC-GWAS to two sets of empirical case-control GWAS summary statistics for the same disorder (for which no case-case associations are expected).

We first assessed the robustness of CC-GWAS to false positive associations due to differential tagging of a causal stress test SNP (Figure 3A); as noted above, CC-GWAS screens the region around every genome-wide significant candidate CC-GWAS SNP for evidence of a differentially linked stress test SNP, and conservatively filters the candidate CC-GWAS SNP when suggestive evidence of a differentially linked stress test SNP is detected (see Methods and Table S6). We simulated GWAS summary statistics using real LD patterns in two distinct populations: British UK Biobank samples and “other European” UK Biobank samples (defined as non-British and non-Irish) (see Methods); we chose these two populations based on their large available sample size of individual-level data (ensuring accurate LD estimates) and similar (or greater) pairwise genetic distance as the pairs of populations in our empirical analyses (see Methods). We simulated causal stress test SNPs and compared four methods/scenarios: CC-GWAS (with filtering) in the scenario where the causal stress test SNP is genotyped/imputed; CC-GWAS (with filtering) in the scenario where the causal stress test SNP is *not* genotyped/imputed; CC-GWAS with no filtering (for which it is irrelevant whether the causal stress test SNP is genotyped/imputed); and direct case-case GWAS (with no filtering). We simulated 10,000 causal stress test SNPs with an average of 417 nearby SNPs in a 100kb window (the range that LD typically spans^28,29^). We report the per-locus type I error rate: the number of loci with at least one genome-wide significant tagging SNP divided by the number of loci tested. The parameter settings were identical to those of the stress test SNPs in Figure 2C, with corresponding sample sizes, CC-GWAS_OLS_ weights and CC-GWAS_Exact_ weights.

**Figure 3.**
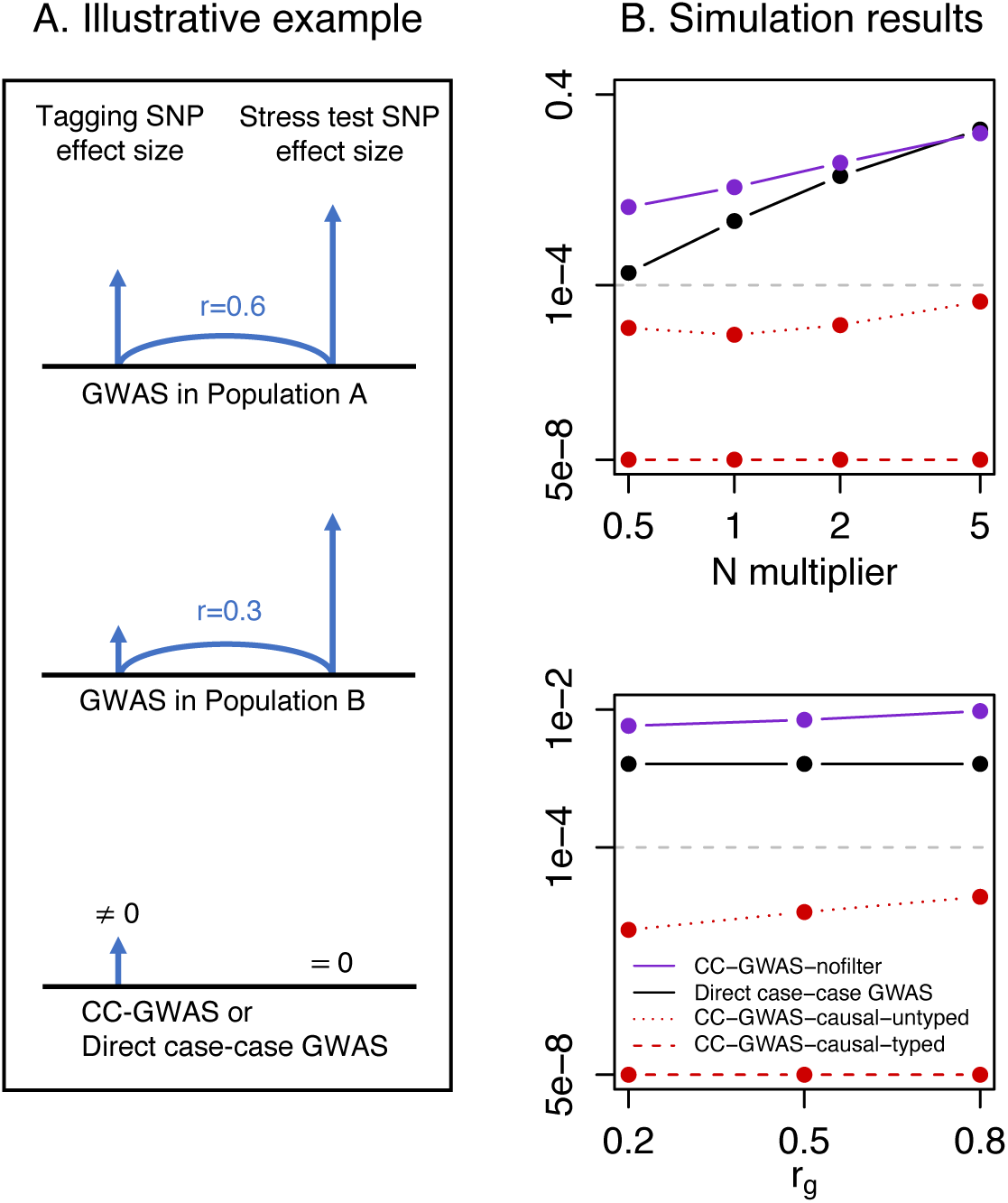
Type I error of CC-GWAS due to differential tagging of a causal stress test SNP. (A) Illustrative example of how differential tagging of a causal stress test SNP can lead to type I error. (B) Simulation results of type I error due to differential tagging. We report the per-locus type I error rate, defined as the number of loci with at least one genome-wide significant tagging SNP divided by the number of loci tested, for each of four methods/scenarios: CC-GWAS, causal stress test SNP genotyped/imputed (denoted CC-GWAS-causal-typed); CC-GWAS, causal stress test SNP not genotyped/imputed (denoted CC-GWAS-causal-untyped); CC-GWAS, no filter; and Direct case-case GWAS (see text). Default parameter settings are: *h*^2^=0.2, prevalence *K*=0.01, and sample size 100,000 cases + 100,000 controls for disorder A; liability-scale *h*^2^=0.1, prevalence *K*=0.15, and sample size 100,000 cases + 100,000 controls for disorder B; *m*=5,000 causal SNPs for each disorder; and genetic correlation *r*_*g*_=0.5 between disorders. Per-locus type I error rates <5 ×10^−8^ were truncated to 5 ×10^−8^ for visualization purposes. All simulation standard errors were 0 for CC-GWAS-causal-typed (zero false positives across all simulations performed); ≤2.7 ×10^−5^ for CC-GWAS-causal-untyped; ≤1.7 ×10^−3^ for CC-GWAS, no filter; and ≤2.1 ×10^−3^ for Direct case-case GWAS. Numerical results are reported in Table S7.

Results are reported in Figure 3B and Table S7. We reached four main conclusions. First, CC-GWAS (with filtering) attained effective control of type I error, with per-locus type I error rate <5 ×10^−8^ in the scenario where the causal stress test SNP is genotyped/imputed. Second, CC-GWAS (with filtering) also attained effective control of type I error, with per-locus type I error rate <10^−4^, in the scenario where the causal stress test SNP is *not* genotyped/imputed. Analogous to our main simulations above, a per-study type I error rate < 0.05 (which is the standard in GWAS using 5 ×10^−8^ as the significance threshold^25^) due to tagging of a causal stress test SNP would allow for the extreme scenario of 500 (500 ×10^−4^=0.05) large-effect stress test SNPs subject to differential tagging that are not genotyped/imputed (we note that 100 stress test SNPs in our simulations already explain 29% of liability variance in disorder B). Third, CC-GWAS with no filtering suffered a per-locus type I error rate of up to 0.07, underscoring the necessity of applying the filter in CC-GWAS. Fourth, the direct case-case GWAS suffered a per-locus type I error rate of up to 0.09. Thus, the robustness of CC-GWAS (with filtering) to differential tagging represents a major improvement over direct case-case GWAS (with no filtering).

We performed four secondary analyses of differential tagging, yielding the following conclusions. First, the per-tagging SNP type I error rate (the number of genome-wide significant tagging SNPs divided by the number of tagging SNPs tested) was much smaller than the per-locus type I error rate (Table S7); however, the per-locus type I error rate is the most appropriate metric, as it takes only one false positive tagging SNP in a stress test locus to draw false positive conclusions about association. Second, when simulating stress test SNPs explaining less variance, CC-GWAS (with filtering) attained effective control of type I error, with per-locus type I error rate <10^−4^ both in the scenario where the causal stress test SNP is genotyped/imputed and in the scenario where the causal stress test SNP is *not* genotyped/imputed (Table S7). Third, type I error rate was well controlled when simulating stress test SNPs explaining varying amounts of variance for specific parameter settings based on all 28 pairs of 8 psychiatric disorders analysed in this study (Table S7). Fourth, when applying various perturbations to the CC-GWAS filtering criteria, the overall conclusions did not change (Table S8).

We next applied CC-GWAS to two sets of empirical case-control GWAS summary statistics for the same disorder, for which no case-case associations are expected. We focused on breast cancer (BC), as this is a disorder with two sets of independent, publicly available GWAS summary statistics in very large sample size (i.e. well-powered for discovery analyses). We compared BC case-control GWAS results of 61,282 cases + 45,494 controls (OncoArray sample in ref.^30^) vs. 46,785 cases + 42,892 controls (iCOGs sample in ref.^30^) (see URLs); these case-control GWAS were both well-powered, identifying 66 and 49 independent loci respectively. To run CC-GWAS, we specified the number of independent causal SNPs at 7,500^31^ (see Methods for a detailed discussion of the assumed number of causal SNPs in applications of CC-GWAS). The CC-GWAS_OLS_ weights and CC-GWAS_Exact_ weights are reported in Table S9, along with the disorder-specific parameters used to derive these weights. The CC-GWAS analyses yielded no genome-wide significant case-case association. Notably, CC-GWAS identified two independent genome-wide significant candidate loci (containing a total of 5 SNPs) prior to filtering for differential tagging of stress test SNPs. All SNPs in these two loci very clearly met the filtering criteria (Table S10), and remained filtered when applying various perturbations to the filtering criteria (Table S8). We emphasize that these BC vs. BC analyses were only intended to test the robustness of CC-GWAS, as CC-GWAS is intended for comparing two *different* disorders with genetic correlation < 0.8 (see Methods). Nevertheless, our analyses of BC vs. BC further validate the robustness of CC-GWAS.

### CC-GWAS identifies 116 loci with different allele frequencies among cases of SCZ, BIP and MDD

We applied CC-GWAS to publicly available summary statistics for SCZ,^15^ BIP^16^ and MDD^17^ (Table 1; see URLs). To run CC-GWAS, we assumed 10,000 independent causal SNPs for each psychiatric disorder^32^ (see Methods for a detailed discussion of the assumed number of causal SNPs in applications of CC-GWAS). The underlying CC-GWAS_OLS_ weights and CC-GWAS_Exact_ weights used by CC-GWAS are reported in Table 1, along with the disorder-specific parameters used to derive these weights. The CC-GWAS_OLS_ weights are based on the expected genetic distances between cases and/or controls (*F*_*ST,causal*_) (Figure 1B-D and Table S11). For each disorder, we specified a range of disorder prevalences to CC-GWAS_Exact_ (Table 1; see *Overview of Methods*). The threshold of genome-wide significance for the CC-GWAS_OLS_ component was set to *P*<5 ×10^−8^ for each pair of disorders (see Discussion).

We defined *independent* genome-wide significant CC-GWAS loci by clumping correlated SNPs (*r*^2^≥0.1) in 3MB windows and collapsing remaining SNPs within 250kb^15^ (Table S12). We defined *CC-GWAS-specific* loci as loci for which none of the genome-wide significant SNPs had *r*^2^>0.8 with any of the genome-wide significant SNPs in the input case-control GWAS results (Table S12). We note that this definition of CC-GWAS-specific loci may include loci previously reported as genome-wide significant in analyses of other case-control data sets (see below). We further note that CC-GWAS loci that are not CC-GWAS-specific also contribute to our understanding of differences between different disorders.

For each pair of SCZ, BIP and MDD, the total number of independent CC-GWAS loci and number of independent CC-GWAS-specific loci are reported in Table 1. The CC-GWAS analysis identified 121 loci, summed across pairs of disorders: 12 for SCZ vs. BIP, 99 for SCZ vs. MDD, and 10 for BIP vs. MDD. CC-GWAS loci were considered shared between two pairs of disorders when at least one pair of genome-wide significant SNPs for the respective pairs of disorders had *r*^2^>0.8 (Table S12). Thus, 4 of the CC-GWAS loci were shared between SCZ vs. BIP and SCZ vs. MDD and 1 was shared between SCZ vs. MDD and BIP vs. MDD, resulting in 116 independent CC-GWAS loci. 5 of the 12 SCZ vs. BIP loci were also significant in the SCZ case-control comparison while none were significant in the BIP case-control comparison (consistent with the larger SCZ case-control sample size); 89 of the 99 SCZ vs. MDD loci were also significant in SCZ case-control comparison while only 1 of those was also significant in the MDD case-control comparison (consistent with the relative genetic distances in Figure 1C); and 5 of the 10 BIP vs. MDD loci were also significant in the BIP case-control comparison while only 1 was significant in the MDD case-control comparison. The remaining 21 (7+10+4) loci were CC-GWAS-specific (and independent from each other); 8 of these loci have not been reported previously (conservatively defined as: no SNP in 1000 Genomes^33^ with *r*^2^>0.8 with a genome-wide significant CC-GWAS SNP in the locus reported for any trait in the NHGRI GWAS Catalog^34^; Table S12).

Notably, the CC-GWAS_Exact_ component did not filter out any variants identified using the CC-GWAS_OLS_ component, i.e. all SNPs with *P*<5 ×10^−8^ using CC-GWAS_OLS_ weights also had *P*<10^−4^ using CC-GWAS_Exact_ weights (for all disorder prevalences in the specified ranges), because the CC-GWAS_OLS_ weights were relatively balanced. In addition, no variants were excluded based on the filtering step to exclude potential false positive associations due to differential tagging of a causal stress test SNP.

For each CC-GWAS locus, the respective input case-control effect sizes for each disorder are reported in Figure 4 and Table S13. Details of the 21 CC-GWAS-specific loci are reported in Table 2, and details of the remaining 100 CC-GWAS loci are reported in Table S13 (the locus names reported in these Tables incorporate results of our SMR analysis^35^; see below). For all 21 CC-GWAS-specific loci, the input case-control effect sizes were non-significant with opposing signs. For CC-GWAS-specific SCZ vs. BIP loci, the input case-control effect sizes had comparable magnitudes for SCZ and BIP, reflecting their similar SNP-heritabilities and prevalences (case-control effect sizes are on the standardized observed scale based on 50/50 case-control ascertainment). For CC-GWAS-specific SCZ vs. MDD and BIP vs. MDD loci, the input case-control effect sizes were smaller for MDD due to its lower SNP-heritability and higher prevalence, implying much lower observed-scale SNP-heritability and *F*_*ST,causal*_ (Figure 1). For the remaining 100 loci, 4 of 5 had opposing case-control association signs in the respective input GWAS results for SCZ vs. BIP, 43 of 89 had opposing signs for SCZ vs. MDD, and 4 of 6 had opposing signs for BIP vs. MDD.

**Table 2.**
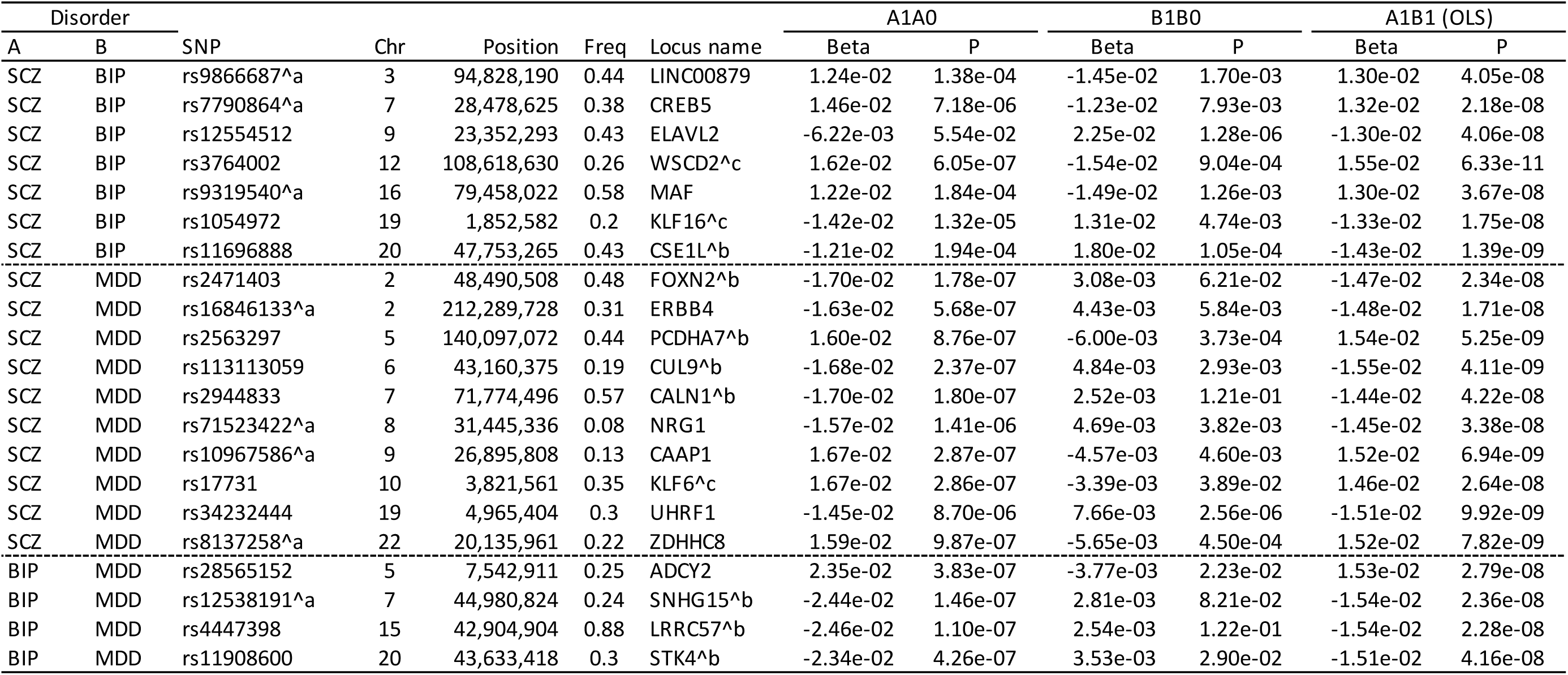
List of 21 CC-GWAS-specific loci for SCZ, BIP and MDD. For each CC-GWAS-specific locus, we report the lead CC-GWAS SNP and its chromosome, physical position, and reference allele frequency, the locus name, the respective case-control effect sizes and *p*-values, and the CC-GWAS_OLS_ case-case effect size and *p*-value. Effect sizes are reported on the standardized observed scale based on 50/50 case-control ascertainment. ^a^denotes loci that have not been reported previously^34^. ^b^denotes locus names based on (most) significant SMR results. ^c^denotes locus names based on exonic lead SNPs. Remaining locus names are based on nearest gene, and do not refer to any inferred biological function. Case-case effect sizes and *p*-values for the CC-GWAS_Exact_ component are reported in Table S13. SCZ, schizophrenia; BIP, bipolar disorder; MDD, major depressive disorder.

**Figure 4.**
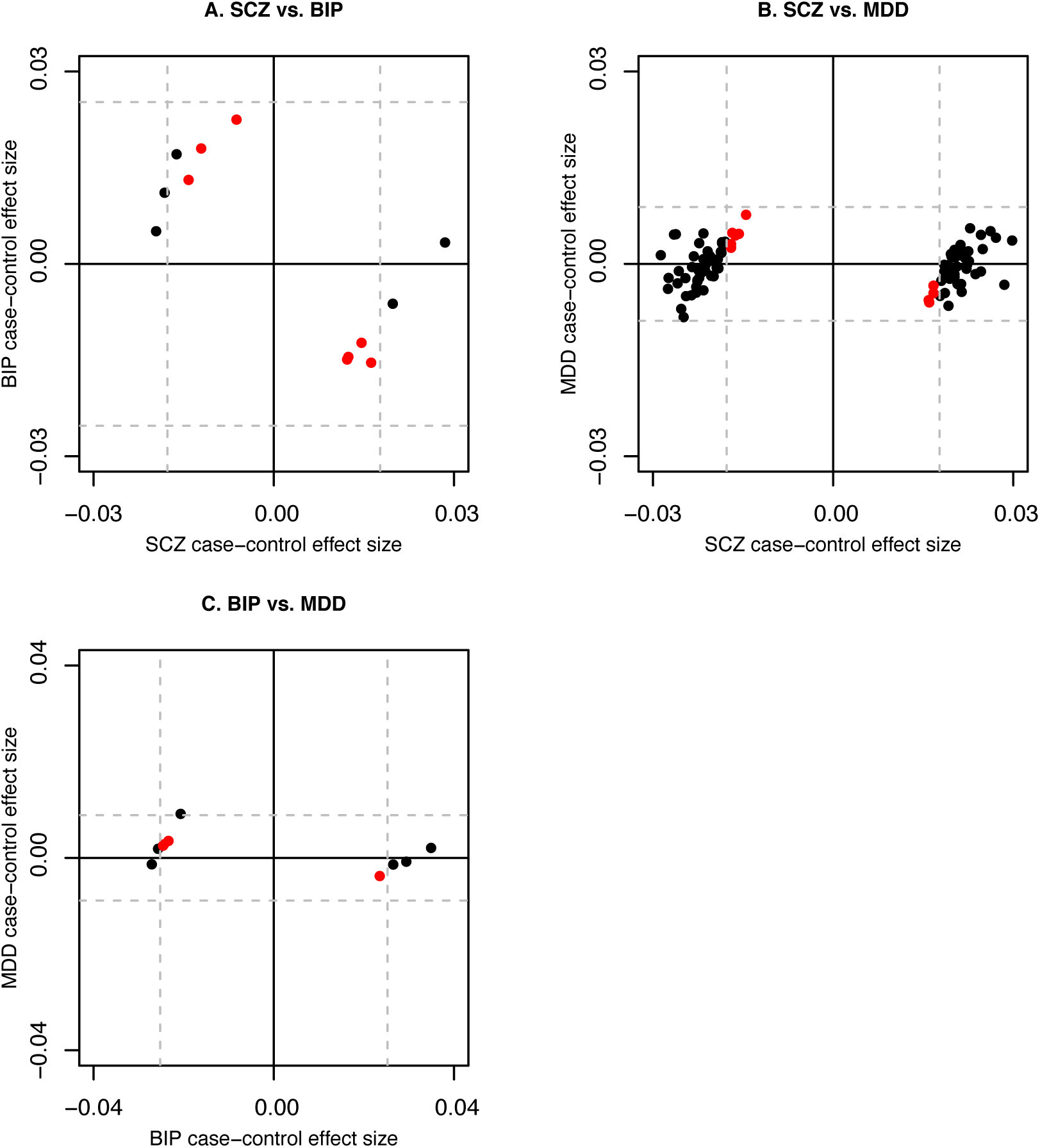
Case-control effect sizes at CC-GWAS loci for SCZ, BIP and MDD. We report the respective case-control effect sizes for lead SNPs at CC-GWAS loci for (A) SCZ vs. BIP, (B) SCZ vs. MDD and (C) BIP vs. MDD. Effect sizes are reported on the standardized observed scale based on 50/50 case-control ascertainment. Red points denote CC-GWAS-specific loci, and black points denote remaining loci. Dashed lines denote effect-size thresholds for genome-wide significance. All red points (denoting lead SNPs for CC-GWAS-specific loci) lie inside the dashed lines for both disorders; in panel A, one black point (denoting the lead SNP for a CC-GWAS locus that is not CC-GWAS-specific) lies inside the dashed lines for both SCZ and BIP, because the lead SNP is not genome-wide significant for SCZ but is in LD with a SNP that is genome-wide significant for SCZ. Numerical results are reported in Table S13. SCZ, schizophrenia; BIP, bipolar disorder; MDD, major depressive disorder.

We performed eight secondary analyses, yielding the following conclusions. First, when we employed a more stringent *p*-value threshold for the CC-GWAS_Exact_ component of CC-GWAS than the default threshold of 10^−4^, significant loci were removed only when a threshold of 10^−6^ or below was employed (Table S14). Second, results were similar when applying a different clumping strategy to define independent loci (Table S15). Third, when conservatively correcting input summary statistics for their stratified LD score regression (S-LDSC) intercept^36–39^ (similar to Turley et al.^6^), the number of significant CC-GWAS independent loci decreased (e.g. from 99 to 75 for SCZ vs. MDD; Table S16). However, we believe this correction is overly conservative, as S-LDSC intercept attenuation ratios^40^ were relatively small, implying little evidence of confounding (Table S17); we note that any confounding, if present, would have a similar impact on CC-GWAS results as on the confounded input summary statistics (when applying no correction for S-LDSC intercept) (see Methods and Table S18). Fourth, results were little changed when estimating heritabilities provided as input to CC-GWAS using LD score regression (LDSC)^36^ instead of S-LDSC^37–39^, despite systematically lower heritability estimates (Table S19); the robustness of these results is consistent with our simulations (Figure S11). Fifth, when changing the number of causal SNPs *m* to 5,000 (resp. 20,000), slightly fewer (resp. slightly more) loci were detected (Table S20). Sixth, results were little changed when varying the disorder prevalences used by CC-GWAS_OLS_ (in addition to the disorder prevalences used by CC-GWAS_Exact_) (Table S21). Seventh, the same number of CC-GWAS-specific loci (21) were identified when applying a more stringent definition of CC-GWAS-specific loci (all genome-wide significant SNPs in the input case-control GWAS results are either >3Mb away, or 250kb-3Mb away with *r*^2^<0.1; Table S12). Eighth, we extended the SCZ vs. BIP analysis by applying CC-GWAS+ (see above) to incorporate case-case summary statistics^4^, but this did not increase the number of independent CC-GWAS-specific loci (Table S22).

### CC-GWAS-specific loci implicate known and novel disorder genes

We used two approaches to link the 21 CC-GWAS-specific loci to genes (Table 2). First, we linked exonic lead SNPs to the corresponding genes. Second, we used the SMR test for colocalization^35^ (see URLs) to identify CC-GWAS loci with significant associations between gene expression effect sizes in *cis* across 14 tissues (13 GTEx v7 brain tissues^41^ and a meta-analysis of eQTL effects in brain tissues^42^; see URLs) and CC-GWAS_OLS_ case-case effect sizes (*P*<0.05 divided by the number of probes tested for each pair of disorders; see Methods), and used the HEIDI test for heterogeneity^35^ to exclude loci with evidence of linkage effects (*P*<0.05) (see Methods and Table S23). Below, we highlight 4 CC-GWAS-specific loci from Table 2, representing both known and novel findings.

The CC-GWAS-specific SCZ vs. MDD CC-GWAS locus defined by lead SNP rs2563297 (chr5:140,097,072) produced significant SMR colocalization results for 11 gene-tissue pairs representing 7 unique genes (Table S23). The 7 unique genes included 5 protocadherin alpha (*PCDHA*) genes, which play a critical role in the establishment and function of specific cell-cell connections in the brain^43^, and the *NDUFA2* gene, which has been associated with Leigh syndrome (an early-onset progressive neurodegenerative disorder)^44^. Significant CC-GWAS SNPs in this locus have previously been associated to schizophrenia^45–48^ (in data sets distinct from our input schizophrenia GWAS^15^, in which this locus was not significant due to sampling variance and/or ancestry differences), depressive symptoms^6^, neuroticism^49^, educational attainment^48,50^, intelligence^51^, blood pressure^52,53^, and a meta-analyses of schizophrenia, education and cognition^48^, implying that this is a highly pleiotropic locus.

The CC-GWAS-specific SCZ vs. MDD locus defined by lead SNP rs2944833 (chr7:71,774,496) produced a significant SMR colocalization result for one gene-tissue pair, involving the *CALN1* gene in meta-analyzed brain eQTL^42^ (Table S23). *CALN1* plays a role in the physiology of neurons and is potentially important in memory and learning^54^. Indeed, SNPs in this locus have previously been associated to educational attainment^50,55^, intelligence^51,56^, cognitive function^57^, and a meta-analysis of schizophrenia, education and cognition^48^. Again, this implies that CC-GWAS can increase power to detect associated loci in the input case-control GWAS data sets analyzed here.

Finally, two distinct CC-GWAS-specific loci implicated genes in the Kruppel-like family of transcription factors. The CC-GWAS-specific SCZ vs. BIP locus defined by lead SNP rs1054972 (chr19:1,852,582) lies within an exon of *KLF16*, and the CC-GWAS-specific SCZ vs. MDD locus defined by lead SNP rs17731 (chr10:3,821,561) lies within an exon of *KLF6*. The respective case-control effect sizes suggest that rs1054972 and rs17731 both have an impact on SCZ, but have not yet reached significance in the respective case-control analyses (P=1.3e–5 and P=2.9e–7 respectively; Table 2 and Table S24). *KLF16* and *KLF6* play a role in DNA-binding transcription factor activity and in neurite outgrowth and axon regeneration^58^, and we hypothesize they may play a role in the previously described schizophrenia pathomechanism of synaptic pruning^59^. Furthermore, the *KLF5* gene from the same gene family has previously been reported to be downregulated in post-mortem brains of schizophrenia patients^60^. At the time of our analyses, *KLF16* and *KLF6* had not previously been associated to schizophrenia; *KLF6* has very recently been associated to schizophrenia in a meta-analysis of East Asian and European populations^47^, but *KLF16* has still not been associated to schizophrenia. This implies that CC-GWAS can identify novel disorder genes.

### CC-GWAS identifies 196 loci distinguishing cases of eight psychiatric disorders

We applied CC-GWAS to all 28 pairs of eight psychiatric disorders by analysing ADHD^18^, AN^19^, ASD^20^, OCD^21^, and TS^22^ in addition to SCZ^15^, BIP^16^ and MDD^17^ (Table 3; see URLs). To run CC-CWAS, we assumed 10,000 independent causal SNPs for each psychiatric disorder^32^ (see Methods for a detailed discussion of the assumed number of causal SNPs in applications of CC-GWAS). The underlying CC-GWAS_OLS_ weights and CC-GWAS_Exact_ weights used by CC-GWAS are reported in Table S14. The CC-GWAS_OLS_ weights are based on the expected genetic distances between cases and/or controls (*F*_*ST,causal*_) (Figure S13 and Table S11). For each disorder, we specified a range of disorder prevalences to CC-GWAS_Exact_ (Table 3; see *Overview of Methods*). The threshold of significance for the CC-GWAS_OLS_ component was set to *P*<5×10^−8^ for each pair of disorders (see Discussion).

**Table 3.**
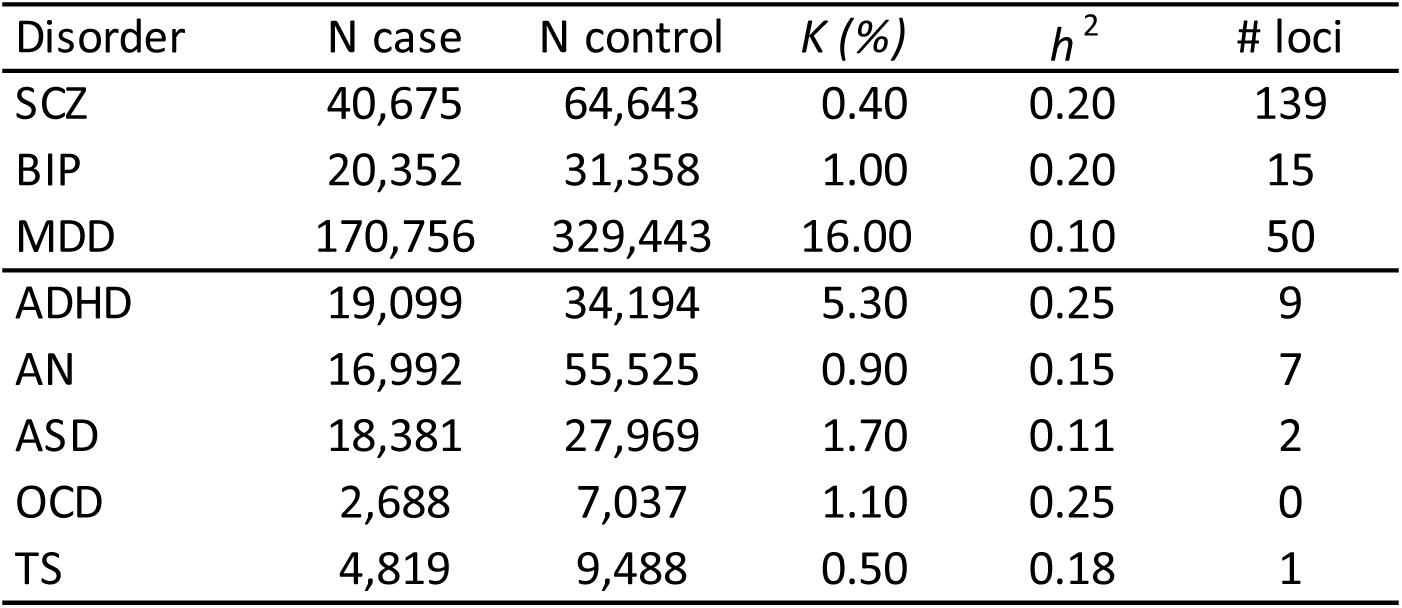
List of eight psychiatric disorders. For each of schizophrenia (SCZ)^15^, bipolar disorder (BIP)^16^, major depressive disorder (MDD)^17^, attention deficit/hyperactivity disorder (ADHD)^18^, anorexia nervosa (AN)^19^, autism spectrum disorder (ASD)^20^, obsessive–compulsive disorder (OCD)^21^, Tourette’s Syndrome and Other Tic Disorders (TS)^22^, we report the case-control sample size, the most likely prevalence (*K*)^84^, liability-scale heritability estimated using stratified LD score regression^37–39^ (*h*^2^), and number of independent genome-wide significant case-control loci. The genetic correlation estimates (another input parameter of CC-GWAS) are presented in Table 4, and the CC-GWAS_OLS_ and CC-GWAS_Exact_ weights are presented in Table S14. We specified a range of prevalences to the CC-GWAS_Exact_ component for SCZ (0.4%-1.0%)^15,84^, BIP (0.5%-2.0%)^16^, MDD (16.0%-30.0%)^17,84^, ADHD (2.5%-10%), AN (0.5%-2.0%), ASD (1.0%-4.0%), OCD (0.5%-2.0%) and TS (0.25%-1.0%) (yielding 2 × 2 = 4 CC-GWAS_Exact_ p-values per comparison, all required to be < 10^−4^).

For each pair of psychiatric disorders, the total number of independent CC-GWAS loci and number of independent CC-GWAS-specific loci are reported in Table 4. The CC-GWAS analysis identified 313 loci, summed across pairs of disorders (0 to 99 loci per pair of disorders). Many of the loci were shared between pairs of disorders, resulting in 196 independent loci; in particular, 49 SCZ case-control loci were shared across 9 pairs of disorders (SCZ and one other disorder), explaining 85 of the 117 overlapping loci. 124 of the 196 loci were also significant in one (or both) of the two input case-control comparisons. The remaining 72 loci were CC-GWAS-specific; 32 (44%) of these loci have not previously been reported in the NHGRI GWAS catalog^34^. The proportion of independent loci that were CC-GWAS-specific (72/196) was larger than in our above analysis of SCZ, BIP and MDD (21/116), but the proportions were more similar when summing across pairs of disorders (83/313 and 21/121, respectively); the difference between 72/196 and 83/313 reflects the fact that CC-GWAS-specific loci are less likely to be shared between pairs of disorders. Notably, the CC-GWAS_Exact_ component (based on the most likely disorder prevalences) filtered out loci identified using the CC-GWAS_OLS_ component for three pairs of disorders: from 9 to 1 for SCZ vs. OCD, from 30 to 19 for SCZ vs. TS, and from 3 to 2 for ADHD vs. OCD, a consequence of highly imbalanced CC-GWAS_OLS_ weights for these specific pairs of disorders due to differences in sample sizes of the input case-control GWAS (Table S14). In addition, one SCZ vs. ASD locus was excluded by specifying a range of disorder prevalences instead of only the most likely prevalences for the CC-GWAS_Exact_ component (reducing the number of SCZ vs. ASD loci from 41 to 40). However, the CC-GWAS_Exact_ component did not filter out any variants identified using the CC-GWAS_OLS_ component for the remaining 24 pairs of disorders. Seven loci were excluded (reducing the number of CC-GWAS loci from 320 to 313) based on the filter to exclude potential false positive associations due to differential tagging of a causal stress test SNP: 1 of 1 locus for SCZ vs. OCD and 6 of 19 loci for SCZ vs. TS (Table S10). All 7 loci were filtered by the criterion specific to small sample sizes (see Methods, Supplementary Note and Table S6), as OCD and TS had small sample sizes (Table 3); 3 of the 7 loci were CC-GWAS-specific. Perturbation of filtering criteria had little impact on results (Table S8 and Supplementary Note). Fewer CC-GWAS-specific loci (63 instead of 72) were identified when applying a more stringent definition of CC-GWAS-specific loci (all genome-wide significant SNPs in the input case-control GWAS results are either >3Mb away, or 250kb-3Mb away with *r*^2^<0.1; Table S12).

**Table 4.** Summary of CC-GWAS results for eight psychiatric disorders. For each pair of disorders, we report the genetic correlation estimated using cross-trait LD score regression^2^ (*r*_*g*_) (lower left) and the number of independent genome-significant CC-GWAS loci (number of CC-GWAS-specific loci in parentheses) (upper right). The CC-GWAS_OLS_ weights and number of SNPs tested are reported in Table S14. SCZ, schizophrenia; BIP, bipolar disorder; MDD, major depressive disorder; ADHD, attention deficit/hyperactivity disorder; AN, anorexia nervosa; ASD, autism spectrum disorder; OCD, obsessive–compulsive disorder; TS, Tourette’s Syndrome and Other Tic Disorders.

| rg\# loci | SCZ | BIP | MDD | ADHD | ANO | ASD | OCD | TS |
| --- | --- | --- | --- | --- | --- | --- | --- | --- |
| SCZ | - | 12 (7) | 99 (10) | 43 (14) | 41 (5) | 40 (10) | 0 (0) | 13 (4) |
| BIP | 0.70 | - | 10 (4) | 8 (6) | 5 (2) | 3 (0) | 1 (1) | 5 (3) |
| MDD | 0.31 | 0.33 | - | 9 (2) | 6 (1) | 3 (2) | 0 (0) | 0 (0) |
| ADHD | 0.16 | 0.18 | 0.44 | - | 4 (3) | 1 (0) | 2 (2) | 2 (2) |
| AN | 0.26 | 0.10 | 0.28 | 0.01 | - | 1 (1) | 0 (0) | 2 (1) |
| ASD | 0.25 | 0.17 | 0.34 | 0.37 | 0.11 | - | 1 (1) | 1 (1) |
| OCD | 0.32 | 0.27 | 0.25 | -0.20 | 0.42 | 0.10 | - | 1 (1) |
| TS | 0.11 | 0.08 | 0.23 | 0.19 | 0.08 | 0.16 | 0.50 | - |

For each CC-GWAS locus, the respective input case-control effect sizes for each disorder are reported in Figure S14 and Table S13. Details of the 72 CC-GWAS-specific loci and of the remaining 124 CC-GWAS are reported in Table S13. For all 72 CC-GWAS-specific loci, the input case-control effect sizes were non-significant with opposing signs. For the remaining 124 CC-GWAS loci, 73 had opposite case-control association signs in the respective input GWAS results. Results of SMR analyses^35^ of the CC-GWAS-specific loci are reported in Table S23.

### CC-GWAS loci replicate in independent data sets

We investigated whether case-case associations identified by CC-GWAS replicate in independent data sets. Of the eight psychiatric disorders, only SCZ and MDD had sufficient sample size to perform replication analyses of the SCZ vs. MDD results based on publicly available GWAS results of subsets of the data^14,61^. The other psychiatric disorders had much lower sample sizes (Table 3), precluding replication efforts based on subset data. For SCZ vs. MDD, we applied CC-GWAS to publicly available summary statistics for subsets of the SCZ data^14^ and MDD data^61^ (discovery data; Table S25; see URLs). We replicated these findings using independent summary statistics constructed by subtracting these summary statistics from the full SCZ data^15^ and full MDD data^17^ using MetaSubtract^62^ (replication data; Table S25). If the discovery data are not exact subsets of the full data, we anticipate that replication results would be conservative, as independent signals from the discovery data would be subtracted from the full data when producing the replication data. To further validate the CC-GWAS method, we also analysed three case-case comparisons of three autoimmune disorders with publicly available GWAS results for independent discovery and replication data sets with substantial sample sizes (Crohn’s disorder (CD)^63^, ulcerative colitis (UC)^63^ and rheumatoid arthritis (RA)^64^; Table S25; see URLs; for CD and UC, replication data were obtained used MetaSubtract^62^; for RA, replication data were publicly available). We assumed *m* = 1,000 independent causal SNPs for each autoimmune disorder^32^. The genetic distances between the autoimmune disorders are displayed in Figure S13; the CC-GWAS_OLS_ weights, CC-GWAS_Exact_ weights and number of CC-GWAS loci and CC-GWAS-specific loci are reported in Table S25; and details of the CC-GWAS loci are reported in Table S26. We replicated the results of the CC-GWAS analysis of the autoimmune disorders (discovery data^63,64^) using independent Immunochip replication data^63,64^, which was available for 62 loci (Table S25).

Results for these four pairs of disorders are reported in Figure 5, Table S25 and Table S26. For SCZ vs. MDD, the CC-GWAS discovery analysis identified 57 independent loci (less than the 99 independent loci in Table 1, due to smaller sample size), of which 53 (93%) had the same effect sign in the CC-GWAS replication analysis and 29 (51%) had same sign and CC-GWAS_OLS_ p-values <0.05. The power of the replication sample for SCZ vs. MDD was considerable smaller than of the discovery sample (effect size SE 2 times larger, corresponding to 4 times smaller effective sample size). The replication slope (based on a regression of replication vs. discovery effect sizes^65^) was equal to 0.57 (SE 0.06) (Figure 5A), which was comparable to the replication slopes for SCZ case-control (0.62, SE 0.05) and MDD case-control (0.60, SE 0.11) genome-wide significant loci using the same discovery and replication data sets (Figure S15, Table S25 and Table S27); we hypothesize that all slopes were smaller than 1 owing to within-disorder heterogeneity^1^. For the autoimmune disorders, 62 independent CC-GWAS loci were available for replication, of which 62 (100%) had the same effect sign in the CC-GWAS replication analysis and 58 (94%) had same sign and CC-GWAS_OLS_ p-values <0.05. For the autoimmune disorders, power for discovery and replication were similar (similar effect size SE and effective sample size). The replication slope for the three autoimmune disorders was equal to 0.83 (SE 0.03) (Figure 5B), comparable to the corresponding case-control replication slopes (Figure S15). We further investigated the replication of the subset of 22 CC-GWAS-specific loci (9 for SCZ vs. MDD and 13 for the 3 autoimmune disorders), pooling the 4 replication studies to overcome the limited number of CC-GWAS-specific loci. We obtained a replication slope of 0.70 (SE 0.07) for the 22 CC-GWAS-specific loci (Figure 5C), which was borderline significantly different (*P*=0.07) from the slope of 0.83 (0.02) for the 97 remaining loci (Figure 5D); we note that CC-GWAS-specific loci had smaller case-case effect sizes and are thus expected to be more susceptible to winner’s curse^66^ (and to attain a lower replication slope).

**Figure 5.**
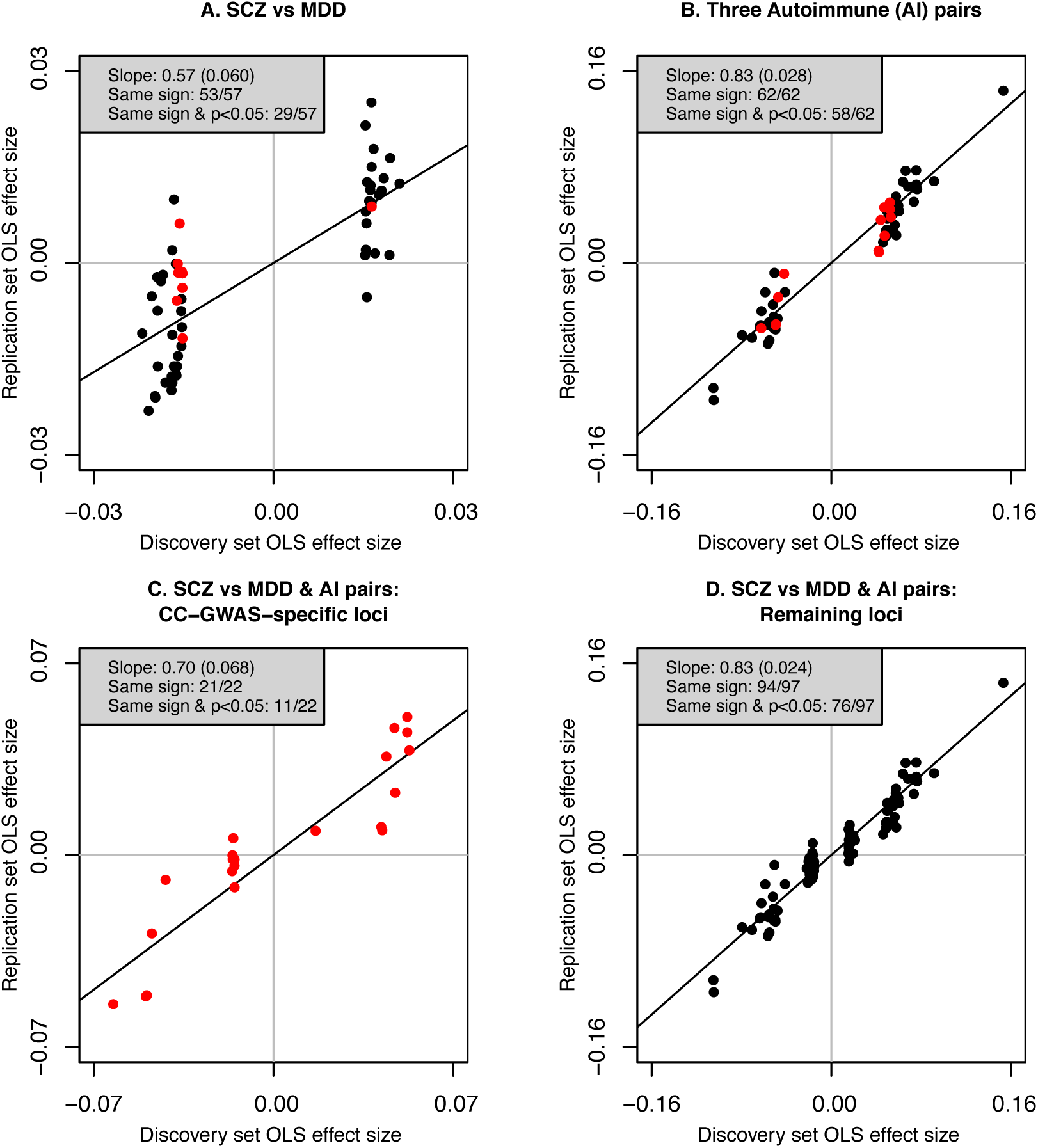
Independent replication of CC-GWAS results. We report replication case-case CC-GWAS_OLS_ effect sizes vs. discovery case-case CC-GWAS_OLS_ effect sizes for (A) schizophrenia (SCZ) vs. major depressive disorder (MDD), (B) three autoimmune disorders, (C) SCZ vs. MDD and three autoimmune disorders, restricting to CC-GWAS-specific loci, and (D) SCZ vs. MDD and three autoimmune disorders, restricting to remaining loci. We also report regression slopes (SE in parentheses), effect sign concordance, and effect sign concordance together with replication *P*_*OLS*_<0.05. Red points denote CC-GWAS-specific loci, and black points denote remaining loci. Numerical results are reported in Table S26, and corresponding case-control replication results are reported in Figure S15 and Table S27.

We sought to perform an additional set of replication analyses using independent replication data that was obtained without requiring the use of MetaSubtract. We focused on breast cancer (BC; see *Assessing the robustness of CC-GWAS*) and RA (see above), as these were disorders with two sets of independent, publicly available GWAS summary statistics in large sample size (i.e. well-powered for replication analyses) without requiring the use of MetaSubtract; although comparing BC cases to RA cases is of low biological interest, this analysis is useful for assessing the robustness of the CC-GWAS method. The CC-GWAS discovery analysis identified 19 CC-GWAS loci. We obtained a replication slope of 0.91 (SE 0.03) across the 19 loci, and a similar replication slope of 0.86 (s.e. 0.11) across the 3 CC-GWAS-specific loci (Figure S16). We conclude that case-case associations identified by CC-GWAS replicate convincingly in independent data sets.

## Discussion

We identified 196 independent loci with different allele frequencies among cases of eight psychiatric disorders by applying our CC-GWAS method to the respective case-control GWAS summary statistics. 116 of these loci had different allele frequencies among cases of the mood and psychotic disorders^3^ (SCZ, BIP and MDD). 72 of the 196 loci were CC-GWAS-specific, highlighting the potential of CC-GWAS to produce new biological insights. In particular, the lead SNPs of two distinct loci were located in exons of *KLF6* and *KLF16*, which have been linked to neurite outgrowth and axon regeneration^58^; we hypothesize that these genes may be involved in the role of synaptic pruning in SCZ^59^. We confirmed the robustness of CC-GWAS via simulations, analytical computations, empirical analysis of BC vs. BC, and independent replication of empirical CC-GWAS results. CC-GWAS excluded 7 of 321 loci across 28 disorder pairs based on its filtering step to discard potential false positive associations that can arise due to differential tagging of a causal stress test SNP, and excluded 1 additional locus by conservatively specifying ranges of disorder prevalences instead of the most likely disorder prevalences.

Although there exist other methods to combine GWAS results from two disorders or complex traits^3,6–13^, CC-GWAS is the first method to compare allele frequencies among cases of two disorders based on the respective case-control GWAS summary statistics – and also the first method to compute the genetic distance between cases and/or controls of two disorders (using *F*_*ST,causal*_). Below, we discuss how CC-GWAS differs from 6 other types of methods. First, the methods of refs.^6–8^ combine GWAS results of correlated disorders or traits to increase power. These methods can be applied to increase the power of case-control analyses, but not to perform case-case analyses; specifically, these methods would require case-case summary statistics to perform a case-case comparison. (Separately, the differences between CC-GWAS+ (which incorporates case-case summary statistics) and MTAG^6^ are discussed above; see *Main simulations*, Supplementary Note and Table S4.) Our CC-GWAS-specific loci are inherently different from the loci identified by these methods, because CC-GWAS computes a weighted difference while those methods compute a weighted sum of the respective case-control GWAS results. Second, the GWIS method^9^ provides a general framework to derive GWAS results from a function of phenotypes, but does not naturally extend to case-case comparisons because it is currently unclear whether case-case effect sizes can be directly expressed as a mathematical function of case-control effect sizes for each disorder (ref.^9^ and personal communication, M. Nivard). Third, the ASSET method^10^ conducts subset-based meta-analyses to increase power and explore subsets of studies for effects that are in the same or possibly opposite directions; CC-GWAS differs from ASSET^10^ by applying weights based on the genetic distance between cases and controls to specifically test difference in allele frequency among cases of two disorders. The method of ref.^11^ investigates results from meta-analysis and provides a statistic representing the posterior predicted probability of association (*m-value*) for each study included in the meta-analysis. The method of ref.^11^ can identify SNPs with predicted disorder-specific effects (i.e. SNPs with a large *m*-value for one disorder and low *m*-value for another disorder): these SNPs are expected to have different allele frequencies among cases, but CC-GWAS models the case-case comparison more directly while explicitly controlling for potential false positive detection of stress test SNPs and providing a formal test of significance. In recent work from the Cross-Disorder Group of the Psychiatric Genomics Consortium^3^, the same eight psychiatric disorders were analyzed with the ASSET method^10^ and the method of ref.^11^ yielding (i) 146 independent genome-wide loci based on the ASSET method^10^, including 109 pleiotropic loci with *m*-values^11^ larger than 0.9 for more than one disorder, and (ii) 11 loci with opposing directions of effect across disorders based on analyses of ASSET-loci with suggestive significance (*P*<10^−6^) with subsequent false discovery rate (*FDR*) correction in *both* of the respective disorders. The 146 ASSET-loci analyzed in ref.^3^ overlapped with 57/196 (29%) of CC-GWAS loci and 5/72 (7%) of CC-GWAS-specific loci (Table S13), confirming that CC-GWAS is different from ASSET (we note that CC-GWAS analyses were based on different input GWAS results for AN, MDD and SCZ). The 11 loci with opposing effects from ref.^3^ are expected to have different allele frequencies among cases, but CC-GWAS is more inclusive as it does not require controlling *FDR* in both of the respective case-control GWAS results and because loci with similar direction of case-control effects can still have different allele frequency among cases of both disorders. Of the 11 loci with opposing effects^3^, 8 overlapped with CC-GWAS loci for the same set of disorders, confirming that CC-GWAS detects these loci as expected while being more inclusive. Fourth, another potential approach for identifying SNPs with disorder-specific effects is to identify all SNPs that have a genome-wide significant effect in at least one of the two disorders and do *not* have a genome-wide significant effect in a meta-analysis of the two disorders. However, we determined that this approach is less powerful than CC-GWAS (Supplementary Note, Figure S17 and Figure S18). Fifth, the *mtCOJO* method^12^ estimates genetic effects conditional on other traits. Some mtCOJO loci and CC-GWAS loci may overlap, but CC-GWAS addresses a conceptually different question than conditional analyses. In ref.^13^, *mtCOJO*^12^ is applied on case-control GWAS results of SCZ, BIP, MDD, ADHD and ASD to identify putative disorder-specific SNPs by correcting GWAS results of one disorder for the causal relationships with the four other disorders. Of the 162 CC-GWAS loci detected in comparisons of these five disorders, 100/162 (62%) of CC-GWAS loci and 8/51 (16%) of CC-GWAS-specific loci overlapped with loci from ref.^13^, confirming that CC-GWAS is different from *mtCOJO*^12^ (we note that CC-GWAS analyses were based on different GWAS results for MDD only). In summary, although some of these previous methods^3,6–13^ identify loci that are expected to have different allele frequencies among cases, none of these methods explicitly compares the allele frequency among cases of different disorders or explicitly controls for potential false positives at stress test SNPs.

Finally, the most natural method to compare CC-GWAS to is a case-case GWAS based on individual-level data, as performed in ref.^4^ for SCZ vs. BIP based on individual level data from ref.^14^ and ref.^16^ respectively. CC-GWAS identified 12 SCZ vs. BIP loci (or 10 when applied to data from ref.^14^ and ref.^16^, as in ref.^4^) compared to 2 SCZ vs. BIP loci identified in ref.^4^, which discarded ∼25% of the cases compared to the respective case-control data (owing to non-matching ancestry and genotyping platform). The 2 loci identified in ref.^4^ were sub-genome-wide significant or genome-wide significant in our CC-GWAS analyses, with same direction of effect (CC-GWAS_OLS_ p=1.9 ×10^−6^ and p=1.6 ×10^−9^, both CC-GWAS_Exact_ p<10^−4^; Table S28). The 12 loci identified by CC-GWAS were all sub-genome-wide significant in the analyses of ref.^4^, with same direction of effect (median p=1.6 ×10^−4^; range 1.4 ×10^−7^ to 5.2 ×10^−2^; Table S29). We estimated a cross-trait genetic correlation^2^ between the results from ref.^4^ and the CC-GWAS_Exact_ results (resp. the CC-GWAS_OLS_ results) of 1.02 (s.e. 0.02) (resp. 0.99; s.e. 0.02). We note these comparisons between results from ref.^4^ and CC-GWAS do not constitute an independent replication, as the samples analyzed in the two studies were largely overlapping, but they do confirm that CC-GWAS is generally concordant with a direct case-case comparison. We further note two advantages of CC-GWAS over a direct case-case GWAS. First, CC-GWAS is much less sensitive to subtle allele frequency differences due to differences in ancestry and/or genotyping platform (Table S30), because the case-case comparison accounts for the allele frequency in matched controls by comparing case-control effects. Second, CC-GWAS filters potential false positive associations due to differential tagging of a causal stress test SNP (with the same allele frequency in cases of both disorders). This is not possible in a direct case-case GWAS based on data from cases alone, as the filtering criteria require information about case-control effect sizes; this filter will become more important as sample sizes continue to increase (Figure 3B).

The CC-GWAS method has several limitations. First, the CC-GWAS_OLS_ case-case effect size estimates depend on the sample sizes of the input case-control GWAS summary statistics, because the CC-GWAS_OLS_ weights depend on these sample sizes. This bias-variance tradeoff can be avoided by using CC-GWAS_Exact_ effect size estimates (which are independent of sample size), e.g. in genetic correlation analyses^2^. Second, the choice of the threshold for the CC-GWAS_Exact_ *p*-values in CC-GWAS is somewhat arbitrary, but we believe 10^−4^ is a reasonable choice as it (i) effectively protects against false positives due to stress test SNPs (Figure 2C and Figure S6), which cannot be numerous (e.g. 100 independent stress test SNPs as defined in Figure 2C would explain 29% of liability-scale variance in disorder B), and (ii) has only limited impact on the power of CC-GWAS (Figure 2A); other choices of this threshold produced identical results in most of our empirical analyses (Table S14). Third, for significant CC-GWAS-specific loci (*P*<5 ×10^−8^) with input case-control *p-*values > 5 ×10^−8^, CC-GWAS does not provide a formal assessment of which case-control effect(s) the locus is associated to. However, the case-control *p*-values at the locus can provide suggestive evidence. In some instances (e.g. SCZ vs. MDD), CC-GWAS-specific loci are likely to largely derive from one of the disorders (SCZ) due to differential power (Figure 1 and Table 2), but this does not limit the value of identifying novel loci distinguishing these disorders. Fourth, we have analyzed 28 pairs of disorders using the significance threshold of 5×10^−8^, which may raise concerns about multiple testing. However, we note that the threshold of 5×10^−8^ has been shown to be much more conservative than false discovery rate^67^ (FDR) approaches^41,68^, and that our use of the 5×10^−8^ threshold is analogous to the use of this threshold in GWAS studies that analyze many traits (e.g. 58 traits in ref.^69^). We specifically verified that, in the data that we analyzed, the threshold of 5×10^−8^ is more conservative than either a per-pair of disorders FDR of 0.05 or a global FDR of 0.05 (Table S31). We note that many of the p-values of the CC-GWAS loci that we identified (including the CC-GWAS-specific loci reported in Table 2) are close to 5×10^−8^ (Table S13), but this is equally true of the case-control GWAS loci for the disorders that we analyzed (Table S32), as this is a general property of polygenic architectures. Fifth, when comparing cases of more than two disorders, CC-GWAS must be applied in pairwise fashion. Extending CC-GWAS to more than two disorders is a direction for future research. Sixth, the CC-GWAS_OLS_ component of CC-GWAS assumes that SNP effect sizes for the two disorders follow a bivariate normal distribution. In principle, CC-GWAS_OLS_ could be extended to account for different sets of causal variants for both disorders^70^; this is a direction for future research, but we note that application of CC-GWAS to two simulated disorders with different sets of causal variants (violating the shared causal variant assumption of CC-GWAS) attained similar power as application of CC-GWAS to two disorders with the same set of causal variants (Table S5). Seventh, we have not explored the application of CC-GWAS to improve case-case polygenic risk prediction. This is an important direction for future research; we note that the question of how to best incorporate cases and controls into case-case polygenic risk prediction is a complex topic^71^, and that analyses of polygenic risk prediction require individual-level validation data. Eighth, we recommend that CC-GWAS should only be applied to compare cases of different disorders with genetic correlation <0.8, as stress test SNPs may occur more often for pairs of disorders with very high genetic correlation; analyses of disorders with very high genetic correlation (including comparisons of the same disorder across sexes^72,73^) is an important direction for future research. Ninth, interpretation of CC-GWAS results depends on the methods used in the respective case-control studies for data collection and analyses. In particular: (i) if one disorder is analyzed using random population controls and the other disorder is analyzed using screened controls, CC-GWAS power and effect size estimates may be impacted and type I error at stress test SNPs may be slightly inflated (although we expect that these effects would be modest, as the genetic distance between cases and controls would generally be much larger than the genetic distance between random population controls and screened controls; in addition, the type I error at null-null SNPs would not be inflated); (ii) if cases of the two disorders have larger comorbidity than expected by chance, CC-GWAS power and effect size estimates will be attenuated; (iii) if a substantial proportion of cases of one disorder are misdiagnosed as having the other disorder, CC-GWAS power and effect size estimates will be attenuated; and (iv) if one or both disorders are corrected for a genetically correlated covariate, this will impact the case-control GWAS results^74^ and thus the interpretation of the CC-GWAS results (however, none of the input case-control GWAS of the eight psychiatric disorders that we analysed were corrected for genetically correlated covariates; Table S33). Tenth, the filtering criteria to avoid false positive associations due to differential tagging of a causal stress test SNP (Table S6) are ad hoc and somewhat arbitrary. However, we verified that applying perturbations to these filtering criteria had little impact on our results, both in extensive simulations (Table S8) and in analyses of empirical data (Table S8). In particular, the filtering criteria were effective (per-locus type I error <10^−4^) in simulations in which the causal stress test SNP was not genotyped/imputed (Figure 3B, Table S7); the type I error in this scenario increases slightly with sample size, but the FDR is more stable (because the true-positive rate increases as well; Figure S19). We believe that results obtained using CC-GWAS are robust to differential tagging of causal stress test SNPs, based on our simulation results and replication in independent data. We note that CC-GWAS is more robust to differential tagging than direct case-case GWAS (Figure 3B). Eleventh, CC-GWAS could in principle be susceptible to confounding effects on allele frequencies due to subtle ancestry differences between the two input case-control GWAS (irrespective of differential tagging). However, for null-null SNPs, we verified that type I error rate is not inflated because there are no case-control allele frequency differences (other than due to sampling variance) in the input case-control GWAS (Table S30). For stress-test SNPs (which cannot be numerous; see above), there is a somewhat increased risk of false positives if the allele frequency difference between populations is on the order of 0.20 for a common SNP (which is typical for differences *between* continental populations^75^), but no increased risk of false positives if the allele frequency differences between populations is on the order of 0.05 for a common SNP (which is typical of differences *within* a continental population^76^) (Table S30); although SNPs can be highly differentiated within a continental population in rare instances^76^, the existence of a SNP that is both a stress-test SNP and highly differentiated within a continental population is very unlikely. Twelfth, CC-GWAS was designed to compare two disorders (with different definitions of controls and potential overlap of cases), but it is also of interest to compare subtypes within a disorder^77^ (with same definitions of controls and no overlap of cases). However, we confirmed via simulation that CC-GWAS can be applied to subtypes (replacing the CC-GWAS_Exact_ approach with the Delta method; Table S34).

In conclusion, we have shown that CC-GWAS can reliably identify loci with different allele frequencies among cases (including both case-control loci and CC-GWAS-specific loci), providing novel biological insights into the genetic differences between cases of eight psychiatric disorders. Thus, CC-GWAS helps promote the ambitious but important goal of better clinical diagnoses and more disorder-specific treatment of psychiatric disorders.

## URLs

CC-GWAS software: https://github.com/wouterpeyrot/CC-GWAS;

CC-GWAS results for 8 psychiatric disorders: https://data.broadinstitute.org/alkesgroup/CC-GWAS/;

R software: https://www.r-project.org/

LDSC software: https://github.com/bulik/ldsc;

SMR software: https://cnsgenomics.com/software/smr/;

PLINK1.9 software: www.cog-genomics.org/plink/1.9/;

GWAS results for BC: http://bcac.ccge.medschl.cam.ac.uk/bcacdata/

GWAS results for ADHD, AN, ASD, BIP, BIP vs. SCZ, MDD (Wray 2018), OCD, SCZ (Ripke 2014), and TS: https://www.med.unc.edu/pgc/results-and-downloads/;

GWAS results for MDD (Howard 2019): https://datashare.is.ed.ac.uk/handle/10283/3203;

GWAS results for SCZ (Pardinas 2018): https://walters.psycm.cf.ac.uk/;

GWAS results for CD and UC: https://www.ibdgenetics.org/downloads.html;

GWAS results for RA: http://www.sg.med.osaka-u.ac.jp/tools.html;

eQTL data of 13 GTEx v7 brain tissues and meta-analysis of eQTL effects in brain tissues: https://cnsgenomics.com/software/smr/#DataResource.

## Methods

### Quantifying genetic distances between cases and/or controls of each disorder

We derive *F*_*ST,causal*_ for the comparisons *A*1*A*0, *B*1*B*0, *A*1*B*1, *A*1*B*0, *A*0*B*1, *A*0*B*0 as follows. Consider two disorders *A* and *B* with lifetime prevalences *K*_*A*_ and *K*_*B*_, liability-scale heritabilities 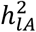 and 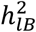, and genetic correlation *r*_*g*_. Assume the heritabilities and genetic correlation have been assessed on data of *m* independent SNPs, and assume these SNPs impact both traits with effects following a bivariate normal distribution. First, the heritabilities are transposed to the observed scales, 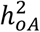 and 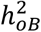, with proportions of cases of 0.5 in line with refs.^24,78^

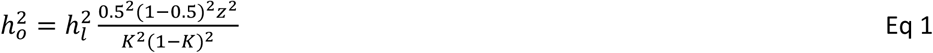

where *z* is the height of the standard normal distribution at threshold *T* defined as *K* = *P*(*x* > *T* | *x*∼*N*(0,1)). The coheritability is also expressed on this scale as: 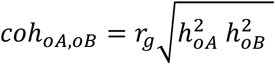. The average variance explained per SNP in *A* is 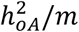 and in *B* 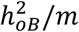, and the average genetic covariance per SNP is *coh*_*oA,oB*_/*m*. For SNP *i*, the allele frequencies of the reference allele in cases and controls are represented as *p*_*i,A*1_, *p*_*i,A*0_, *p*_*i,B*1_,and *p*_*i,B*0_.

Throughout the paper, the effect-sizes *A* are chosen to be on the standardized observed scale (i.e. with standardized genotype and standardized phenotype) based on 50/50 case-control ascertainment. There are two advantages of using the standardized observed scale instead of the liability scale. The first advantage is that computing case-control (A1A0 and B1B0) effect sizes on this scale from case-control summary statistics does not require knowledge of disease prevalence, whereas computing case-control effect sizes on the liability scale from case-control summary statistics requires knowledge of disease prevalence. The second advantage is that this scale is applicable to case-case effect sizes, whereas we are currently unaware of any way to apply the liability scale to case-case effect sizes. We further note that we use the 50/50 case-control ratio to define the standardized observed scale so that effect sizes on this scale do not depend on sample case-control ratios. When assuming Hardy-Weinberg equilibrium and assuming small effect sizes (typical for polygenic disorders), the *β* of linear regression of standardized case-control status *A*1*A*0 on standardized genotype *G*_*i*_ can be approximated in terms of allele frequencies as

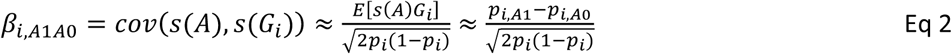

Simulations confirm that these approximations are justified for loci with 0.5 < *OR* < 2 (Table S35). For comparison, all estimated effect sizes for SCZ^15^, BIP^16^ and MDD^17^ have 0.7 < *OR* < 1.4.

The derivations of *F*_*ST,causal, A*1*A*0_ and *F*_*ST,causal, B*1*B*0_ follow from Eq 2. On the standardized scale the variance explained equals the square of the beta. Because the loci are assumed independent, the average variance explained gives

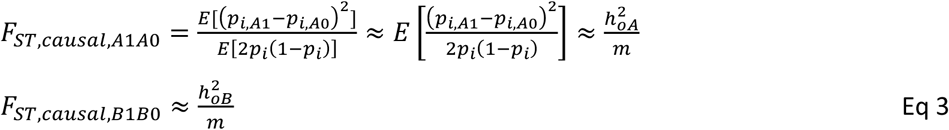

The first approximations has been proposed^75^ to obtain stable estimates of *F*_*ST*_ and assumes the standardized SNP effects are equally distributed across the allele frequency spectrum.

Deriving *F*_*ST,causal*_ for the comparisons *A*1*B*1, *A*1*B*0, *A*0*B*1 and *A*0*B*0 requires some additional steps. First note the covariance of *β* _*i, A*1*A*0_ and *β* _*i, B*1*B*0_ equals

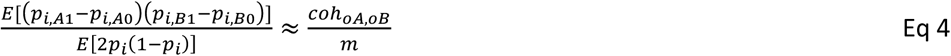

Second, note that allele frequencies and difference in allele frequencies can be rewritten as

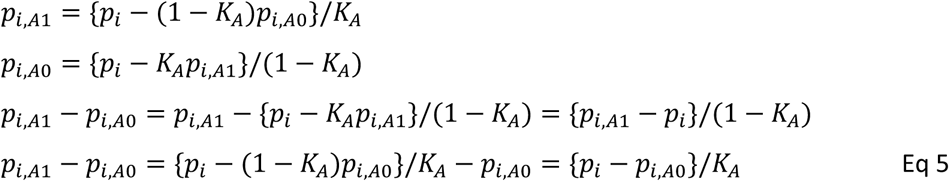

Substituting Eq 5 in Eq 3 and Eq 4, gives

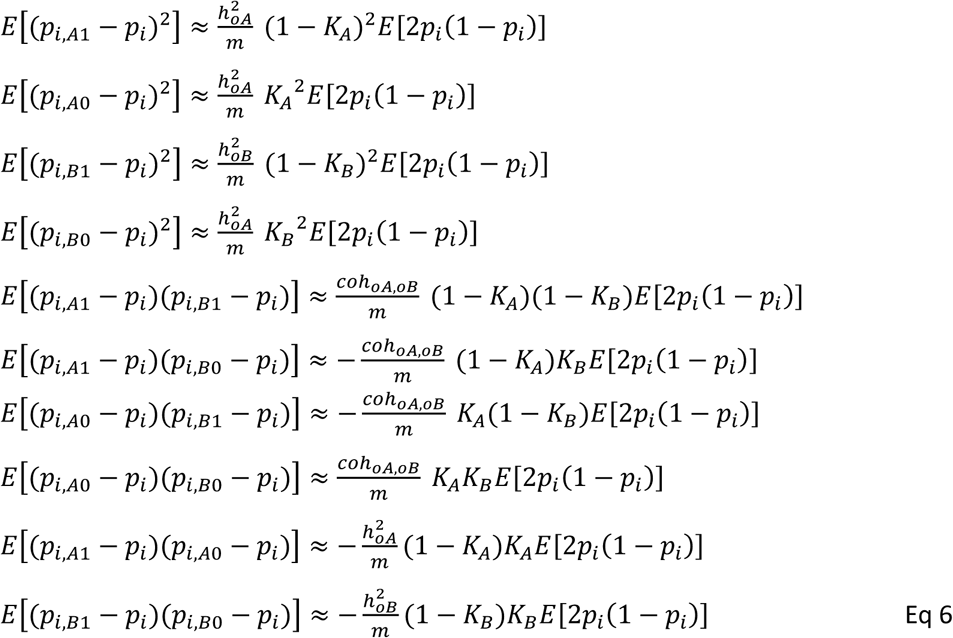

While noting that that the *F*_*ST,causal,XY*_ is estimated as

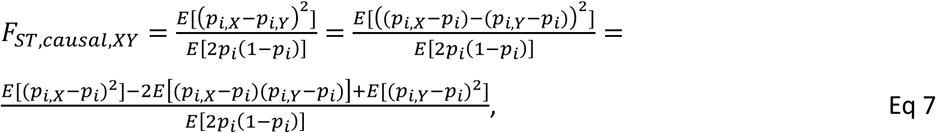

we find

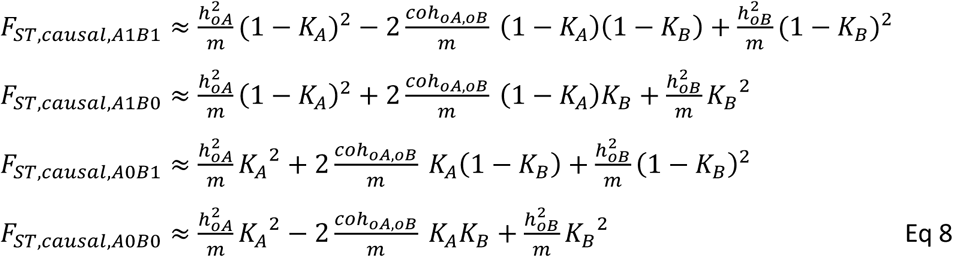

while noting that 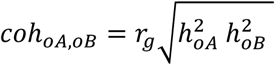. These approximations of *F*_*ST,causal*_ were confirmed with simulations (Table S36). (CC-GWAS depends on the assumption that all *m* SNPs have an impact on both traits and that effects sizes follow a bivariate normal distribution (see below). However, we note that the equations of *F*_*ST,causal*_ require less stringent assumptions. To illustrate, the equations also hold when simulating data (see below) of 1,000 independent SNPs of which 500 have no impact on either disorder, 154 have uniform and similar sized effects on both disorders, 282 have uniform effects on disorder A only, and 64 have uniform effects on disorder B only (Table S36B). Thus, *F*_*ST,causal,A*1*B*1_ between cases of both disorders does not depend on the assumption that the two disorders have the same set of causal variants, but instead depends on the genetic correlation (see also Eq 8) reflecting the similarity of both disorders. We note the same genetic correlation (and *F*_*ST,causal,A*1*B*1_) can be obtained when (i) all causal SNPs impact both traits with relatively lower concordance of effect sizes, or when (ii) a part of the causal SNPs impact both disorders with relatively higher concordance of effect sizes and a part of the causal SNPs impact only one of both disorders.^70^)

We now proceed with using *F*_*ST,causal*_ to display *A*1, *A*0, *B*1 and *B*0 in a 2-dimensional plot (Figure 1 and Figure S13). Because the loci are assumed independent, 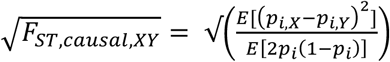 can be interpreted as a Euclidian distance measure in an *m*-dimensional space, where the 4 points *A*1, *A*0, *B*1 and *B*0 are defined by their *m* allele frequencies (e.g. 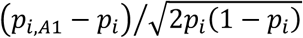 represents the coordinate of point *A*1 on the axis corresponding to SNP *i*). The allele frequency in the full population represents the origin (population mean). While realizing that the lines (*A*1-*A*0) and (*B*1-*B*0) must both go through the population mean, one can see that *A*1, *A*0, *B*1, *B*0 and the population mean can be represented in a 2-dimensional plot. From Eq 3 and Eq 6, it follows that the distance between the population mean and *A*1 (resp. *B*1) equals 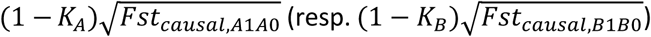. While noting that the distance between *A*1 and *B*1 equals 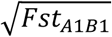, we know the lengths of the three sides of triangle defined by the population mean, *A*1 an *B*1. The law of cosines gives the angle of the lines (population mean - *A*1) and (population mean - *B*1) as

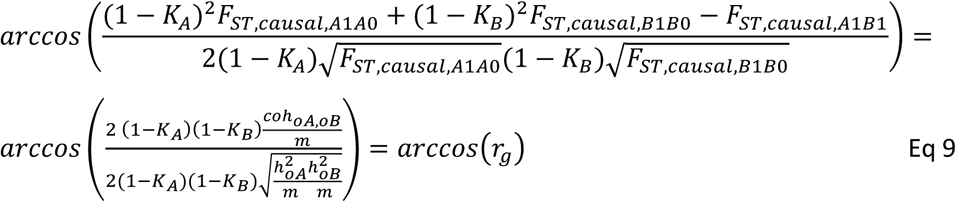

Thus, the genetic correlation *r*_*g*_ is equal to the cosines of the angle of the lines (population mean - *A*1) and (population mean - *B*1). It can analogously be shown that the angle between (population mean - *A*0) and (population mean -*B*0) is the same, and that the angle between (population mean - *A*1) and (population mean - *B*0) equals 180 minus the angle between (population mean - *A*1) and (population mean - *B*1), which confirms the use of *F*_*ST,causal*_ to display *A*1, *A*0, *B*1, *B*0 and the population mean in a 2-dimensional plot. To aid further interpretation, the perpendicular projection of line (*A*1 − *A*0) on line (*B*1 − *B*0) has a length equal to *r*_*g*_ times length (*A*0-*A*1) (i.e. 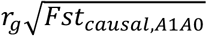), because *r*_*g*_ equals the cosines between these lines.

In application, we derive *F*_*ST,causal*_ analytically based on the heritabilities, population prevalences and genetic correlation. We note two important differences between *F*_*ST,causal*_ and the *F*_*ST*_ from population genetics^75^. First, we restrict our definition of *F*_*ST,causal*_ to independent SNPs, while *F*_*ST*_ from population genetics is based on all genome-wide SNPs. If one where to extend *F*_*ST,causal*_ to genome-wide SNPs, *F*_*ST,causal*_ at loci with large LD-scores would be larger than at SNPs with low LD-scores due to tagging. In contrast, the *F*_*ST*_ from population genetics is mainly attributable to drift and more or less evenly distributed over the genome (except for small effects of selection). Second, *F*_*ST,causal*_ between cases and controls is of the order of magnitude of 10^−6^ depending on the number of SNPs *m* considered. In contrast, the *F*_*ST*_ between European and East Asian has been estimated^75^ at 0.11. Because of the low magnitude of *F*_*ST,causal*_, we report *m* ∗ *F*_*ST,causal*_ in Figure 1 and Figure S13 (note that *m* ∗ *F*_*ST,causal*_ is independent of *m* when other parameters are fixed, because the equations for *F*_*ST,causal*_ have *m* in the denominator (see Eq 3 and Eq 8)).

The purpose of *F*_*ST,causal*_ is to aid intuition to the bivariate genetic architecture of two disorders and to develop the CC-GWAS method (see further), and we do not provide formal standard errors of *F*_*ST,causal*_. However, an approximation of the standard errors of *F*_*ST,causal*_ can be obtained from the standard errors of the estimates of the heritabilities and co-heritability (typically assessed with methods like LD score regression). For *Fst*_*A*1*A*0_ and *Fst*_*B*1*B*0_, this follows directly from Eq 3. For *Fst*_*A*1*B*1_, we assume that the error of the three terms in Eq 8 are independent. This assumption is likely violated, but it serves in obtaining approximations of the standard errors of *F*_*ST,causal*_ (reported in the legend of Figure 1).

### CC-GWAS method

The CC-GWAS method relies on *F*_*ST,causal*_, and assumes that all *m* SNPs impact both disorders with effect sizes following a bivariate normal distribution. CC-GWAS weights the effect sizes from the respective case-control GWAS using weights that minimize the expected squared difference between estimated and true A1B1 effect sizes; we refer to these as ordinary least squares (CC-GWAS_OLS_) weights. To obtain the CC-GWAS_OLS_ weights, we analytically derive the expected coefficients of regressing the causal effect sizes *A*1*B*1 on the GWAS results of *A*1*A*0 and *B*1*B*1

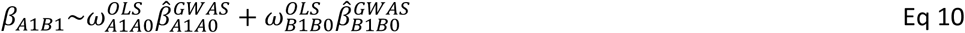

Ordinary least square (OLS) regression gives

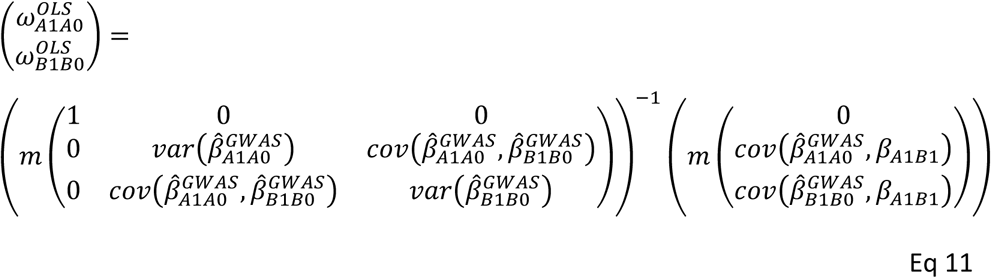

When assuming error terms are independent from effect sizes, we find

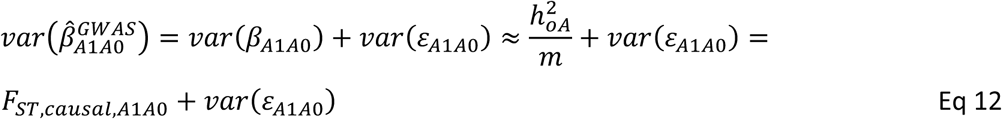

and the analogue for 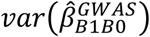. For the covariance of the GWAS results, we find based on Eq 4

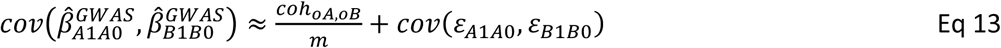

(At the end of this section, we discuss the expectation and estimation of the variance and covariance of error terms as well as scaling of odds ratios to standardized observed scale based on 50/50 case-control ascertainment.) The expectation of the covariance between the GWAS results and A_*A*1*B*1_ follow from Eq 6 as

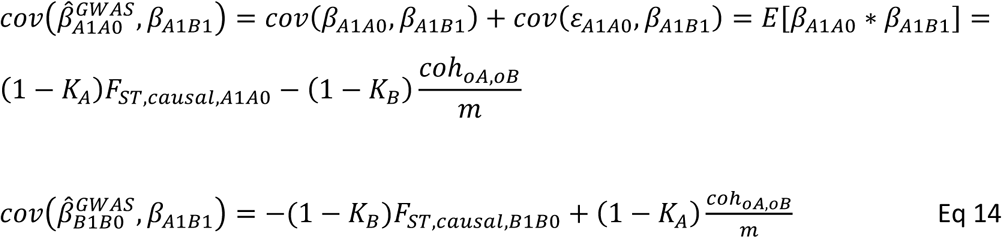

Thus, the CC-GWAS_OLS_ weights are defined in Eq 11 to minimize the expected squared distance between estimated and causal effect sizes *β*_*A*1*B*1_.

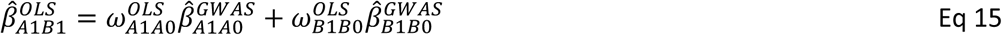

To summarize, the CC-GWAS_OLS_ weights depend on the number of independent causal SNPs, the heritabilities, population prevalences, the genetic correlation, and the variance and covariance of error terms of the betas (depending on sample sizes *N*_*A*1_, *N*_*A*0_, *N*_*B*1_, *N*_*B*0_ and the sample overlap between *A*0 and *B*0).

The CC-GWAS_OLS_ weights may be susceptible to type I error for SNPs with nonzero A1A0 and B1B0 effect sizes but zero A1B1 effect size, which we refer to as “stress test” SNPs (see further). To mitigate this, CC-GWAS also computes sample size independent weights based on infinite sample size; we refer to these as CC-GWAS_Exact_ weights. The CC-GWAS_Exact_ weights depend only on the population prevalences *K*_*A*_ and *K*_*B*_. From Eq 2 it follows that

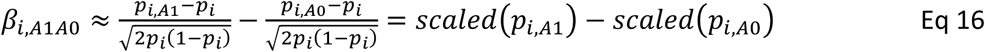

Multiplying with (1 − *K*_*A*_) and substituting *p*_*i,A*0_ based on Eq 5, gives

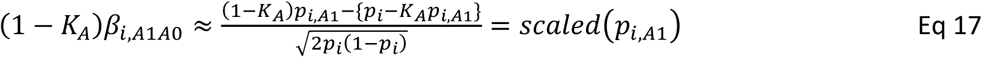

From this, *β*_*i,A*1*A*1_ follows as

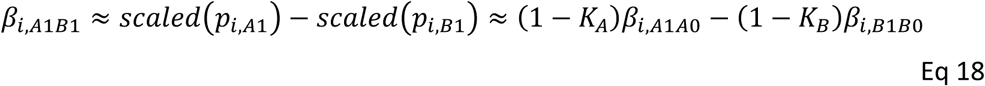

We note that simulations confirm that this approximation is justified for loci with 0.5 < *OR*< 2 for both *A* and *B* (Table S35). The CC-GWAS_Exact_ weights thus follow as

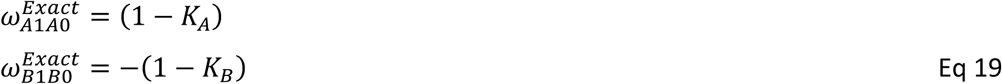

and

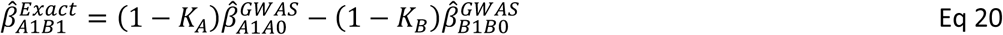

The p-values of the CC-GWAS_OLS_ component and CC-GWAS_Exact_ component are estimated as follows. First note that the standard error of 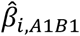 of the CC-GWAS_OLS_ component and CC-GWAS_Exact_ component at SNP *i* follow as

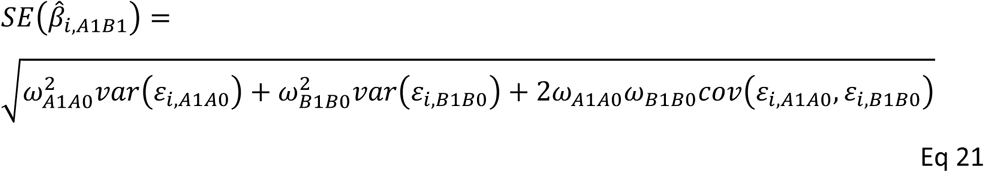

While noting that *ω*_*A*1*A*0_ > 0 and *ω*_*B*1*B*0_ < 0, this indicates that sample overlap of controls (introducing positive covariance between *ε*_*i,A*1*A*0_ and *ε*_*i,B*1*B*0_) will increase power of CC-GWAS. Betas and their standard errors give 0-values, from which p-values follow (assuming normally distributed error terms). CC-GWAS reports a SNP as statistically significant if it achieves 2 < 5 ∗ 10^−8^ using CC-GWAS_OLS_ weights *and* 2 < 10^−4^ using CC-GWAS_Exact_ weights (all statistical tests in this paper are two-sided), balancing power and type I error. We note that CC-GWAS is intended for comparing two *different* disorders with genetic correlation < 0.8. At larger genetic correlation, the anticipated number of stress test loci may increase posing a risk of per-study type I error > 0.05, and the CC-GWAS_OLS_ weights may become meaningless when the expected genetic distance between cases is close to 0.

When GWAS results are available for a direct case-case GWAS, 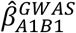, CC-GWAS can be extended to CC-GWAS+. The CC-GWAS+_OLS_ weights are defined as

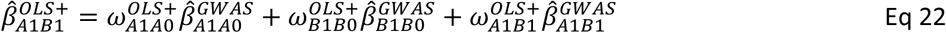

The CC-GWAS+_OLS_ weights follow analogue to Eq 11 for the CC-GWAS_OLS_ weights while noting that

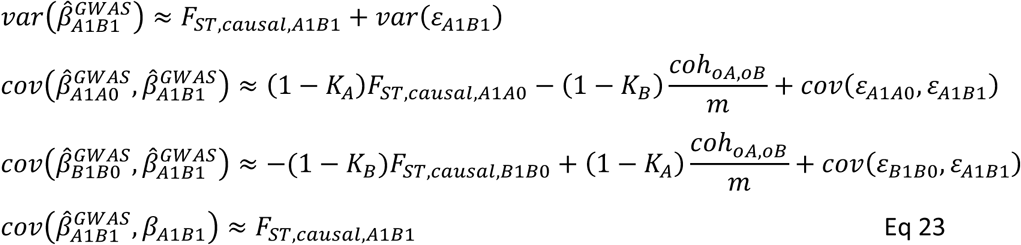

The standard error 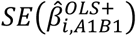 follows as

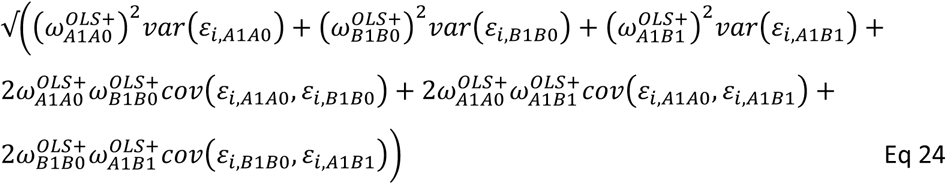

The CC-GWAS+_Exact_ component is simply defined as 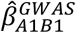 (i.e. 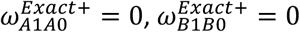 and 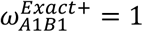). CC-GWAS+ reports a SNP as statistically significant if it achieves 2 < 5 ∗ 10^−8^ using CC-GWAS+_OLS_ weights *and* 2 < 10^−4^ using CC-GWAS+_Exact_ weights.

We now derive the expectation of the variance and covariance of the error terms of the betas. Assume the GWAS results are based on *N*_*A*1_ (resp. *N*_*B*1_) cases and *N*_*A*0_ (resp. *N*_*B*0_) controls of disorder A (resp. disorder B). First, while assuming small effect loci (typical for polygenic disorders), note that the variance of the allele frequency of SNP *i* in *X* (with *X* representing one of *A*1, *A*0, *B*1, *B*0) can be approximated by

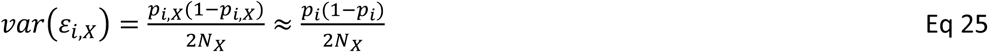

From Eq 2, we find

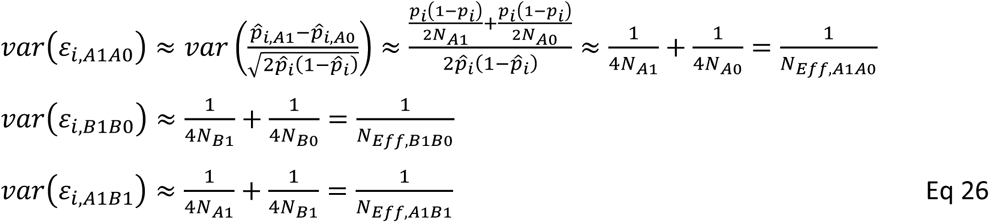

When assuming an overlap of *N*_*overlap,A*0*B*0_ of controls, the expectation of the covariance of the error can be derived as follows. First, note that the error term of the allele frequency in all controls *A*0 can be expressed in terms of the error terms of (*A*0, *in overlap*) and (*A*0, *not in overlap*)as

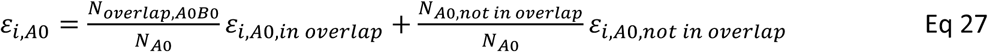

Thus, the covariance of error terms follows as

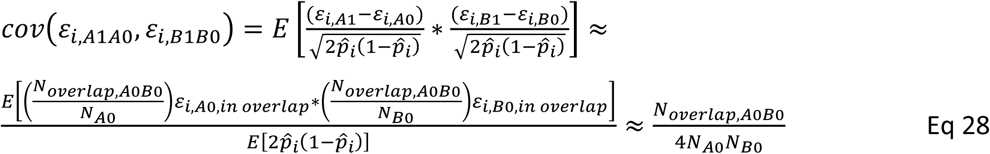

Now, assume of number of *N* _*A*1 *in A*1*B*1_ and *N* _*B*1 *in A*1*B*1_ are available for a direct case-case GWAS included in CC-GWAS+ (typically *N* _*A*1 *in A*1*B*1_ < *N* _*A*1_ and *N* _*B*1 *in A*1*B*1_ < *N* _*B*1_). This gives the following expected covariance of error terms

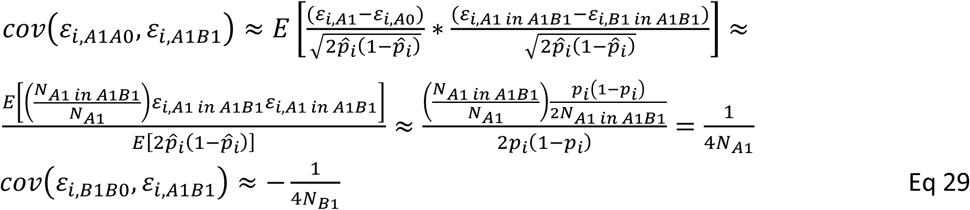

Thus, the expectations of the variance and the covariance of the error terms are given. We note that in practical application of CC-GWAS, the covariances of error terms are based on analytical computation and the intercept of cross-trait LD score regression (described in the section ‘*Application of CC-GWAS to empirical data sets*’).

In practice, case-control GWAS results are not presented on the standardized observed scale based on 50/50 case-control ascertainment (*β*_*i*_) but as odd ratios (*OR*_*i*_) from logistic regression. We apply two approaches to transpose results from logistic regression to *β*_*i*_. First, we assume for large sample sizes that the *z*-value from logistic regression is equal to the *z*-value from linear regression on the observed scale. With the expected variance of error-terms derived as 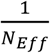 (Eq 26), we find

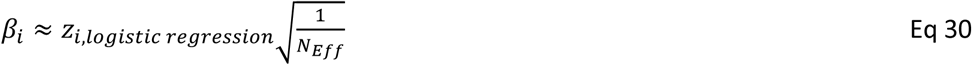

The second approach is based on equation 5 in the paper from from Lloyd-Jones et al.^79^, which reads for 50% cases as

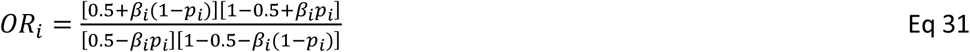

Rewriting gives

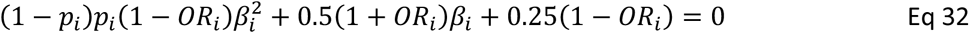

The solution from this quadratic equation with 0 < *β*_*i*_ < 2 for *OR*_*i*_ > 1, and −2 < *β*_*i*_ < 0 for *OR*_*i*_ < 1, gives an approximation of *β*. We confirm in simulation (see below) that both approaches approximate *β*_*i*_ well, but the first approach is slightly less noisy. In the CC-GWAS software, we therefore transform the betas with the first approach, and compare these to transformations with the second approach to provide a rough double-check of whether *N*_*Eff*_ has been defined accurately.

### Filtering criteria to exclude potential false positive associations due to differential tagging of a causal stress test SNP

CC-GWAS identifies and discards false positive associations that can arise due to differential tagging of a causal stress test SNP (Figure 3A). Specifically, CC-GWAS screens the 1MB region around every genome-wide significant candidate CC-GWAS SNP for evidence of a differentially linked stress test SNP, and conservatively filters the candidate CC-GWAS SNP when suggestive evidence of a differentially linked stress test SNP is detected. The criteria of the filtering step were motivated by extensive simulations (Table S7). Here we provide an intuitive overview of the criteria (details are provided in the Supplementary Note and Table S6).

For each candidate CC-GWAS SNP, filtering comprises of three sets of criteria (A, B, and C), and the SNP is discarded when at least one of the three sets of criteria is met. The criteria (A) are intended for intermediate sample sizes, the criteria (B) for relatively small sample sizes, and the criteria (C) for very large sample sizes.

In criteria (A), CC-GWAS considers, as a potential stress test SNP, the SNP in a 1Mb region (around the candidate CC-GWAS SNP) with the largest product of case-control z-scores across the two disorders. The candidate CC-GWAS SNP is filtered as potential false positive association when *all* of the following criteria A1, A2 and A3 are met:

A1. The potential stress test SNP is likely to have the same population allele frequencies among cases of the two disorders, reflected by a CC-GWAS_Exact_ p-value larger than 10^−4^.
A2. The potential stress test SNP is likely to be the causal SNP for both disorders, reflected by absolute case-control z-scores almost as large (allowing for sampling variance) as the largest absolute case-control z-scores in the region for both disorders.
A3. The case-control z-scores at the candidate CC-GWAS SNP and at the potential stress test SNP have a pattern concordant with differential tagging. (See the new Supplementary Note and Table S6 for details.)

The second set of criteria (B) is intended for when at least one case-control GWAS is underpowered, in which case the most likely potential stress test SNP is no longer the SNP considered in (A), due to sampling variance. Specifically, if the case-control GWAS of at least one disorder is underpowered (*N*_*eff*_ < 40*k*), CC-GWAS *additionally* considers, as a potential stress test SNP, the SNP in a 1Mb region with the largest maximum absolute case-control z-score across the two disorders. The CC-GWAS SNP is filtered as a potential false positive association when the following criterium B1 is met:

B1. The potential stress test SNP is likely to have the same population allele frequencies among cases, reflected by a CC-GWAS_Exact_ p-value larger than 10^−4^. (We note that we do not include criteria analogous to A2 and A3, because the error around the z-scores is too large. We also note that when the causal stress test SNP has small effects, GWAS samples with *N*_*eff*_ ≥ 40*k* may also be underpowered; however, the per-locus type I error rate is already < 10 ^−4^ before applying the CC-GWAS filtering step in this scenario; see Table S7.)

The third set of criteria (C) is primarily intended for when the case-control GWAS are very well powered and the causal stress test SNP is *not* genotyped/imputed. In this case, very subtle tagging differences between the potential stress test SNP in criteria (A) and the causal stress test SNP can lead to *p*_*Exact*_ < 10 ^−4^ (i.e. violation of criterium (A1) and thus wrongfully not filtering the candidate CC-GWAS SNP with criteria (A)). Therefore, the CC-GWAS SNP is filtered when the following criterium C1 is met (irrespective of sample size):

C1. The power of CC-GWAS is much lower than the power of the respective case-control analyses, reflected by a much smaller CC-GWAS_Exact_ z-score than the corresponding case-control z-scores.

We note that the filtering criteria assume one large-effect causal stress test SNP in the 1MB region. In theory, if there exists one *causal stress test SNP* of large effect and one large-effect *causal non-stress test SNP* (i.e. with non-zero case-case effect) of even larger effect in the same region, filtering of false positive association due to differential tagging of the causal stress test SNP may be less effective, due to incorrect specification of the potential causal stress test SNP (thus not meeting criterium A1). However, we believe this is exceedingly unlikely in practice, as this would require all of (i) the presence of a causal stress test SNP, (ii) differential tagging of the causal stress test SNP, and the presence of a causal non-stress test SNP of even larger effect in the same region. Furthermore, even in this case, there would be a false-positive association only at the SNP level, and not at the region level (since the existence of a causal non-stress test SNP implies a true case-case association in the region). Thus, we believe it is appropriate to assume only one large-effect causal stress test SNP in our filtering algorithm.

### Main CC-GWAS simulations to assess power and type I error

We simulated individual level data of *m* independent SNPs in line with ref.^78^ for disorders *A* and *B* as follows. Liability-scale effect sizes (*β* _*i,lA*_ and *β* _*i,lB*_) were drawn from a bivariate normal distribution with variances 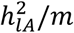 and 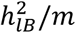 and 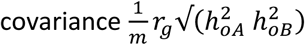. Effective allele frequencies (*EAF*) of *m* SNPs were drawn from a uniform distribution [0.01,0.5]. Individuals were simulated one-by-one by

1. Randomly assigning *m* genotypes *G*_*i*_ (i.e. 0, 1 or 2 effective alleles) with the probabilities given by the *EAFs* while assuming Hardy-Weinberg equilibrium
2. Defining genetic liabilities as 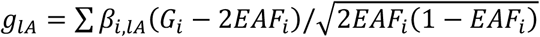, and 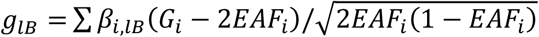
3. Defining liabilities as *l*_*A*_ = *g*_*lA*_ + *e*_*lA*_ and *l*_*B*_ = *g*_*lB*_ + *e*_*lB*_, with *e*_*lA*_ drawn from a standard normal distribution with variance 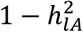, and *e*_*lB*_ drawn from a standard normal distribution with variance 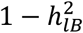. (Here we simulate *e*_*lA*_ and *e*_*lB*_ to be uncorrelated, but note that correlation of *e* _*lA*_ and *e* _*lB*_ does not impact simulation results.)
4. Defining disorder status as *A* = 1 (resp. *B* = 1) when *l*_*A*_ > *T*_*A*_ (resp. *l*_*B*_ > *T*_*B*_) with *T*_*A*_ (resp. *T*_*B*_) corresponding to a population prevalence of *K*_*A*_ (resp. *K*_*B*_)

Individuals were simulated until the required number of nonoverlapping cases and controls (*N*_*A*1_, *N*_*A*0_, *N*_*B*1_, *N*_*B*0_) were obtained. Subsequently, GWASs *A*1*A*0 and *B*1*B*0 were performed with logistic regression in Plink 1.9^80^, CC-GWAS was applied as described above, and the power of CC-GWAS, the CC-GWAS_OLS_ component, the CC-GWAS_Exact_ component and the delta method were recorded. A second set of 3 * *m* **null-null SNPs** with no effect on *lA*. and *lB* were included in the simulation to estimate the respective type I error rates of null-null SNPs. Then, we simulated 3 * *m* **stress test SNPs** as follows. We defined a stress test SNP to explain a proportion of *β*_*lA*_ of variance on the liability scale of *A*, and a proportion of *α*_*0A*_ on the observed scale via the standard transformarion^24,78^. The effect of the stress test SNP on the observed scale follows as 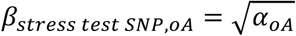, and the allele frequency in cases of A is approximated by (1 − *K*_*A*_) *β*_*stress test SNP, oA*_ (Eq 17) and the allele frequency in controls of A by − *K*_*A*_*β* _*stress test SNP,o A*_. The allele frequency in cases of B is equal to (1 − *K*_*A*_) *β*_*stress test SNP, oA*_, thus giving *β*_*stress test SNP, oB*_ = (1 − *K* _*A*_) *β*_*stress test SNP, oA*_/(1 − *K*_*B*_) (Eq 17). The allele frequency in controls is approximated by −*K* _*B*_*β*_*stress test SNP, oB*_. We simulated cases and controls in line with these allele frequencies. Thus, simulating 3 * *m* stress test SNPs allowed recording of type I error of stress test SNPs. Simulations were repeated 50 times.

The parameters of simulation were largely in line with those used in Figure 2, but in order to reduce computational time, sample sizes were reduced to *N*_*A*1_ = *N*_*A*0_ = *N*_*B*1_ = *N*_*B*0_ = 4,000, number of causal SNPs to *m* = 1,000, and required levels of significance were reduced to *p* < 0.01 for the CC-GWAS_OLS_ component and *p* < 0.05 for the CC-GWAS_Exact_ component. Three values of genetic correlation were simulated (0.2, 0.5 and 0.8). Simulation results are displayed in Table S2 and match analytical computations (described below). The concordance of simulation and analytical computations confirms that increasing sample sizes and decreasing p-value thresholds in analytical computations in Figure 2 is justified. We also simulated data with a different bivariate architecture, with the distribution of SNP effects in line with the general distribution applied in Frei et al.^70^: 1/3 of causal SNPs have an impact on disorder A only, 1/3 of SNPs have an impact on disorder B only, and 1/3 of SNPs have an impact both disorder A and disorder B (the correlation of these SNP effects specify the genome-wide genetic correlation, as in Frei et al.^70^; Table S5).

### CC-GWAS analytical computations to assess power and type I error

For analytical computations, we first consider one sets of weights to derive expected results for the CC-GWAS_OLS_ component, CC-GWAS_Exact_ component, and delta method respectively:

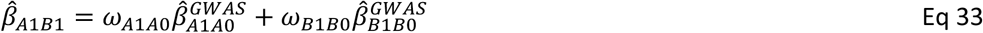

The variance of betas and error terms follow as

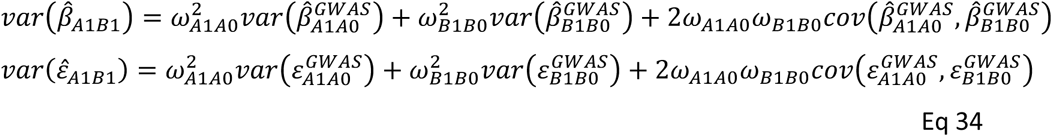

the analytical expectations of which have been derived above. The variance of 0-values at causal loci follows as 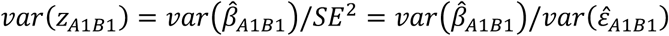, and the power as 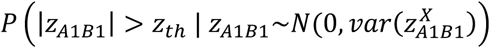 with *z*_*th*_ = *ϕ*^-1^(1 − *p*_*threshold*_/2) and *ϕ*^-1^ the standard normal quantile function (with *p*_*threshold*_ = 0.01 to compare to simulation and *p*_*threshold*_ = 5 * 10^−8^ in Figure 2). Type I error rate at null-null loci is well controlled (i.e. equals p-value) while noting that 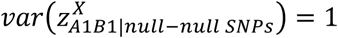. Type I error at stress test SNPs follows as *p*(|*z*_*A*1*B*1_| > *z*_*th*_ | *z* _*A*1*B*1_∼ *N* (*ω* _*A*1*A*0_ *β*_*stress test SNP, oA*_ + *ω* _*B*1*B*0_*β*_*stress test SNP, oB*_,1).

Now consider CC-GWAS combining the estimates of 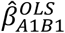 and 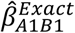 requiring 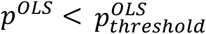 and 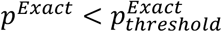 (with corresponding 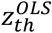 and 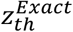) for significance (with thresholds set at 0.01 and 0.05 to compare to simulation, and 5 * 10^−8^ and 10^−4^ in Figure 2). The covariance of 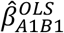 and 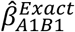 follows as

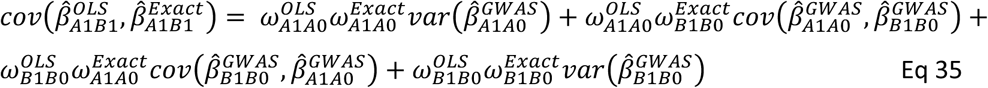

the analytical expectations of which are given in the above. The covariance of the error terms 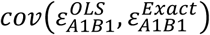 follow analogously, and the covariance of z-values follows at causal SNPs as 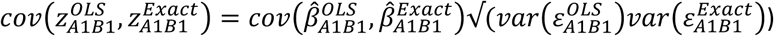. In combination with the expectations of 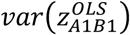 and 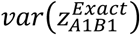, this defines the variances and covariance of the bivariate normal distribution 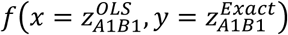 around mean (0,0) at causal SNPs. The power follows as

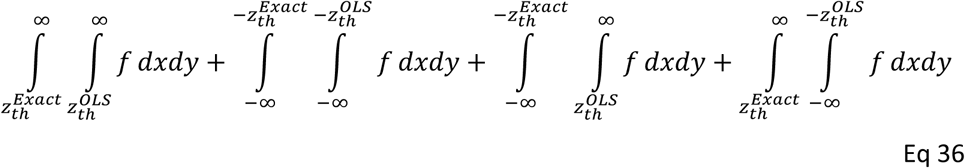

We note that the last two terms (i.e. with the CC-GWAS_OLS_ component and CC-GWAS_Exact_ component meeting the required level of significance at opposite sign) have negligible magnitudes. The type I error at null-null SNPs is found by substituting in the equation above the bivariate normal distribution *f′* with mean (0,0) and variance covariance of z-values at null-null SNPs, i.e. 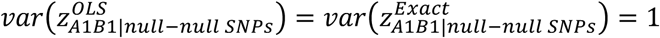 and 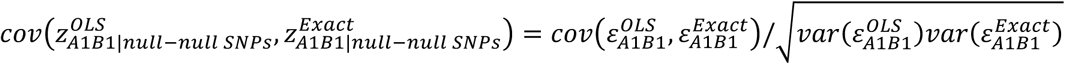. The type I error at stress test SNPs is found by substituting in the equation above the bivariate normal distribution *f〞* with mean (*α*_*A*1*A*0_*β*_*stress test SNP, oA*_ + *α*_*B*1*B*0_*β*_*stress test SNP, oB*_ (1 − *K*_*A*_)*β*_*stress test SNP, oA*_ + (1 − *K*_*B*_)*β*_*stress test SNP, oB*_) and the same variance covariance as of 0-values at null-null SNPs.

The difference in power between CC-GWAS and a direct case-case GWAS equals

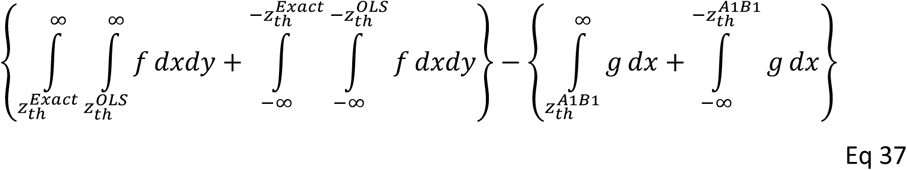

where 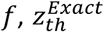 and 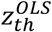 are defined as in Eq 36, and the function *g* (representing z-scores of the direct case-case GWAS) follows a normal distribution with mean 0 and variance 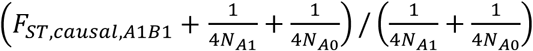 (Eq 23 and Eq 26). Note that the CC-GWAS_OLS_ component reflects the genome-wide significance threshold, thus 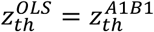.

In secondary analyses, we investigate the impact of overlap in controls. For simulation, we therefore exchanged double controls (*A* = 0 and *B* = 0) between those selected as *A* 0 and *B* 0 selected by chance (thereby preventing the impact of double screening of controls^81^). For analytical computations, we simply adjusted the covariance of error terms in line with the equation above. We also assessed CC-GWAS using the type S error rate, defined as the proportion of significantly identified loci (true positives) identified with the wrong sign^26,27^. We therefore extended the bivariate normal distribution *f* at causal loci to 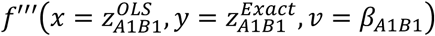 with an additional dimension covering the causal effects *β* _*A*1*B1*_, while noting that *var*(*β*_*A*1*B*1_) =*Fst*_*A*1*B*1_ and 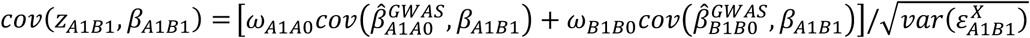 and mean (0,0,0). The type S error follows as

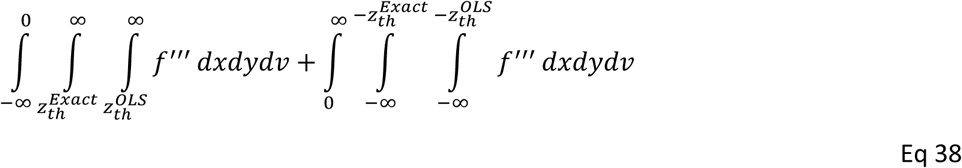

When a direct case-case GWAS is available, CC-GWAS can be extended to CC-GWAS+, the derivations of which follow analogue to the above and are not shown here. These analytical computations were confirmed with our Main simulations (Table S2).

### Simulation of false positive associations due to differential tagging of a causal stress test SNP

We performed extensive simulation to study the impact of potential false positive associations due to differential tagging of a causal stress test SNP. We used real LD patterns in two distinct populations: 25k British UK Biobank samples and 25k “other European” UK Biobank samples (defined as non-British and non-Irish); we note that the *F*_*ST*_ between these two populations is 0.0006^75^, which is greater than the range of *F*_*ST*_ values in the 3 CC-GWAS comparisons of psychiatric disorders for which in-sample allele frequencies were available to estimate *F*_*ST*_ (0.0001-0.0005; Table S37). An overview of LD (signed correlation) differences between 25k British UK Biobank samples and 25k “other European” UK Biobank samples is provided in Table S38. Other parameters were based on our main stress test SNP simulations (Figure 2C; section ‘*Main CC-GWAS simulations to assess power and type I error’*). We selected 10,000 independent SNPs on chromosome 1 as stress test SNPs, and simulated GWAS results^82^ for all 1000 Genome SNPs^33^ within a 100kb radius (the range that LD typically spans^28,29^) with MAF>0.01. The stress test loci included on average 417 SNPs (range 62-800).

Specifically, for each simulated region, we defined the causal stress test SNP (*K*1) with effect sizes *β*_*ST, A*1*A*0_ and *β*_*ST, B*1*B*0_. The average marginal effect size (i.e. marginal effect size with infinite sample size) for each other SNP *i* in the region was given by *β* _*i*_, _*A*1*A*0_= *β* _*ST, A*1*A*0_ * *r* _*ST, i|population A*_ and by *β* _*i*_, _*B*1*B*0_= *β* _*ST, B*1*B*0_ * *r* _*ST, i|population B*_, where *r* is the LD (signed correlation) with the stress test SNP in the respective population. The scaled allele frequencies in infinite sample size in cases and controls of disorder A and disorder B were approximated from Eq 18. The error terms of the scaled allele frequencies in *X* (with *X* representing one of *A*1, *A* 0, *B*1, *B*0) were drawn from a multivariate normal distribution 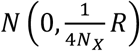 where R is the LD matrix (of signed correlations) in the respective population (see Eq 26 and ref.^82^). The simulated GWAS effect size estimates follow from Eq 26 as 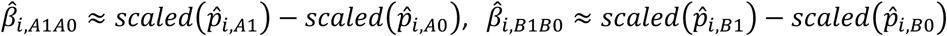 and 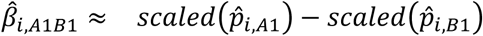. We verified that simulating GWAS summary statistics in this way (based on allele frequencies in cases and controls) yields similar variance/covariance structures of GWAS results as when simulating individual-level data based on real UK Biobank genotypes (data not shown). The advantage of simulating GWAS summary statistics is that this allows increasing the number of simulation runs and sample size dramatically compared to simulations based on individual-level data.

Based on the simulated GWAS results, we first applied CC-GWAS twice including the filtering step: once with the causal stress test SNP included in the GWAS results (CC-GWAS-causal-typed), and once excluding the causal stress test SNP (and all SNPs in perfect LD in both populations) from the GWAS results (CC-GWAS-causal-untyped). Subsequently, we applied CC-GWAS without the filtering step (CC-GWAS-nofilter), and reported the results from 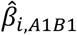 (Direct case-case GWAS). We repeated these analyses 10,000 times per parameter setting and report the per-locus type I error rate: the number of loci with at least one genome-wide significant tagging SNP divided by the number of loci tested. The simulation results are reported in Figure 3B and Table S7.

We performed several secondary simulation analyses. First, we also reported the per-tagging SNP type I error rate (the number of genome-wide significant tagging SNPs divided by the number of tagging SNPs tested) (results in Table S7). Second, we varied the proportion of liability variance explained by the stress test SNP in disorder A from 10 ^−3^ (as in the main simulations) to 10 ^−4^ and 10 ^−5^ (results in Table S7). Third, we repeated simulation for the specific parameter settings based on all 28 pairs of 8 psychiatric disorders analysed in this study (parameter settings in Table S14; results in Table S7). We thus simulated 102 parameter settings covering a wide range of real-world case-case comparisons: 3 different effect-sizes of the stress-test SNP x {6 settings for Figure 3 + 28 settings for the 8 psychiatric disorders}. Fourth, we investigated 16 perturbations of the filtering criteria (results in Table S8).

### Empirical data sets

We compared cases from SCZ^15^, BIP^16^, MDD^17^, ADHD^18^, AN^19^, ASD^20^, OCD^21^, and TS^22^ based on publicly available case-control GWAS results (see URLs). To further validate CC-GWAS we also compare cases from BC^30^, CD^63^, UC^63^ and RA^64^, based on the case-control GWAS results from samples genotyped on chips with genome-wide coverage (see URLs). Numbers of cases and controls are listed in Table 1 and Table S14. The transformation of odds ratios to the standardize betas on the observed scale (Eq 30) requires *N*_*eff*_ (Eq 26). For some of the disorders (BIP, MDD, AN and RA), *N*_*eff*_ was provided on a SNP-by-SNP basis in publicly available GWAS results. For other disorders (SCZ, ADHD, ASD, OCD, TS, BC, CD and UC), we approximated a genome-wide fixed *N*_*eff*_ by summing the *N*_*eff*_ of the contributing 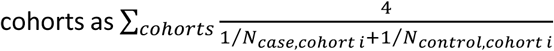. In quality control SNPs were removed with *MAF*< 0.01, *INFO* < 0.6, *N*_*eff*_ < 0.67 * *mAx*(*N*_*eff*_), duplicate SNP names, strand-ambiguous SNPs, and the MHC region (chr6:25,000,000-34,000,000) was excluded due to its compilated LD structure. All reported SNP names and chromosome positions are based on GRCh37/hg19. (In principle, applying a fixed *N*_*eff*_ for cohorts without SNP-by-SNP *N*_*eff*_ information could lead to inaccurate transformation of beta for some SNPs. Therefore, we reran CC-GWAS analyses for SCZ, BIP and MDD with fixed *N*_*eff*_ yielding nearly identical results to the primary analyses (with fixed *N*_*eff*_ for SCZ and SNP-by-SNP *N*_*eff*_ for BIP and MDD). This confirms that using fixed *N*_*eff*_ is appropriate.

### Application of CC-GWAS to breast cancer

To further assess robustness of CC-GWAS, we applied CC-GWAS to BC case-control GWAS results of 61,282 cases + 45,494 controls (OncoArray sample in ref.^30^) vs. BC case-control GWAS results 46,785 cases + 42,892 controls (iCOGs sample in ref.^30^). Input parameters of CC-GWAS are the population prevalences^83^, liability-scale heritabilities 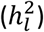^37–39^, genetic correlation (*r*_*g*_)^2^, the intercept from cross-trait LD score regression^2^ (used to model covariance of the error-terms), the sample size including overlap of controls (*N*_*overlap*_= 0; also used to model covariance of the error-terms; see below), and expectation of the number of independent causal SNPs (*m*). The number of independent causal SNPs was set at *m* = 7,500^31^ (see below for a detailed discussion of the assumed number of causal SNPs in applications of CC-GWAS). The resulting CC-GWAS_OLS_ weights and CC-GWAS_Exact_ weights are reported in Table S9.

### Application of CC-GWAS to psychiatric and other empirical data sets

Input parameters of CC-GWAS are the population prevalences (*K*), liability-scale heritabilities 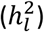, genetic correlation (*r*_*g*_), the intercept from cross-trait LD score regression^2^ (used to model covariance of the error-terms), the sample sizes including overlap of controls (also used to model covariance of the error-terms; see below), and expectation of the number of independent causal SNPs (*m*). Prevalences are displayed in Table 1 and Table S14 and were based on ref.^84^ for the eight psychiatric disorders, on ref.^85^ for UC and CD, and ref.^64^ for RA. Heritabilities were assessed with stratified LD score regression based on the baseline LD v2.0 model^37–39^, and transposed to liability-scale^24,78^. Genetic correlations were estimated with cross-trait LD score regression^2^. The number of causal SNPs was set at *m* = 10,000 for the psychiatric disorders, and *m* = 1,000 for CD, UC and RA based on ref.^32^.

We note that the intercept of cross-trait LD score regression (representing covariance of 0-values at null-null SNPs^6^) can analytically be derived as

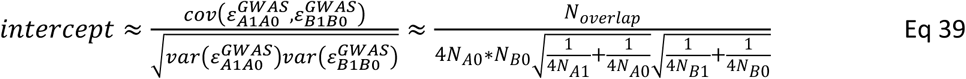

The intercepts estimated with cross-trait LD score regression^2^ were typically larger than those expected analytically based on Eq 39 (Table S39) based on sample-overlap. This could roughly be attributable to (i) underestimation of sample-overlap (ii) factors increasing the cross-trait LD score regression intercept other than covariance of error terms based on sample-overlap. For example, Yengo et al. showed that shared population stratification may increase the intercept^86^. In addition, it is known that attenuation bias of univariate LD score regression can increase the univariate LD score intercept^40^, and we hypothesize this phenomenon may also increase the cross-trait LD score regression intercept. Importantly, we note that overestimation of the covariance of error terms will underestimate the standard error of CC-GWAS results (Eq 21) thereby risking increased false positive rate. Therefore, in CC-GWAS we model the covariance of error terms based on the minimum of the intercept from cross-trait LD score regression and the expected intercept based on Eq 39.

Based on the listed input parameters, CC-GWAS (software provided in R^87^; see URL section) was applied one disorder pair at a time. CC-GWAS first estimates *F*_*ST,causal*_ and plots *A*1, *A*0, *B*1, and *B*0 as described above (and displayed in Figure 1 and Figure S13). Second, CC-GWAS transforms case-control input GWAS results (on the per-allele *OR* scale) to the standardized observed scale based on 50/50 case-control ascertainment in two ways as described above (via *N*_*eff*_ and via the equation from Lloyd-Jones et al.^79^). The transformations were in concordance with correlation between betas of both approaches > 0.985 and only slight differences in magnitude with relative differences > 0.9 and < 1.1. Third, CC-GWAS_OLS_ weights and CC-GWAS_Exact_ weights (displayed in Table 1 and Table S14) were computed as described. Fourth, based on these weights, CC-GWAS_OLS_ estimates 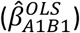 and CC-GWAS_Exact_ estimates 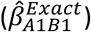 were computed with accompanying standard errors and p-values as described above. Fifths and finally, CC-GWAS reports SNPs as statistically significant that achieve *P*<5×10^−8^ using CC-GWAS_OLS_ weights *and P*<10^−4^ using CC-GWAS_Exact_ weights.

CC-GWAS results were clumped in line with ref.^15^ based using 1000 Genomes data^33^ as LD reference panel with Plink 1.9^80^ (--clump-p1 5e-8 --clump-p2 5e-8 --clump-r2 0.1 --clump-kb 3000; see URLs) (Table S12). Loci within 250kb of each other after the first clumping step were collapsed. We defined *CC-GWAS-specific* loci as loci for which none of the genome-wide significant SNPs have an *r*^2^>0.8 with any of the genome-wide significant SNPs in the input case-control GWAS results (Table S12). We chose this value because we think it is unlikely that a CC-GWAS locus would statistically result from a significant case-control locus for which all significant SNPs have r^2^≤0.8 with all significant SNPs in the CC-GWAS locus. An overview of the number of CC-GWAS loci is given in Table 1 and Table S14, and details are reported in Table 2 and Table S13.

Secondary analyses included a different clumping strategy in line with ref.^40^ with Plink 1.9 (--clump-p1 5e-8 --clump-p2 5e-8 --clump-r2 0.01 --clump-kb 5000) while subsequently collapsing loci within 100kb of each other. In another set of secondary analyses input case-control summary statistics were corrected for the respective intercepts of stratified LD score regression (similar to Turley et al.^6^) by dividing the input case-control standard errors by 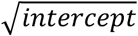 and adjusting the *z*-values and *p*-values accordingly. We believe this correction may be overly conservative, because some increase of the intercept may be expected due to attenuation bias of imperfect matching of LD patterns^40^. Nevertheless, we verified as follows that CC-GWAS provides proportionally biased results when applied on biased input GWAS results (i.e. CC-GWAS does not introduce an additional layer of bias). First, in simulation we multiplied the standard errors with *c*_*A*1*A*0_ = *c*_*B*1*B*0_ = 0.9 and verified that the increase in 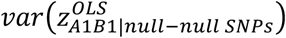 was proportional to the increase in 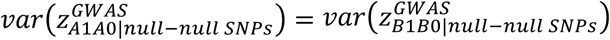 under various scenarios (Table S18). Second, we note this bias can also be analytically derived as

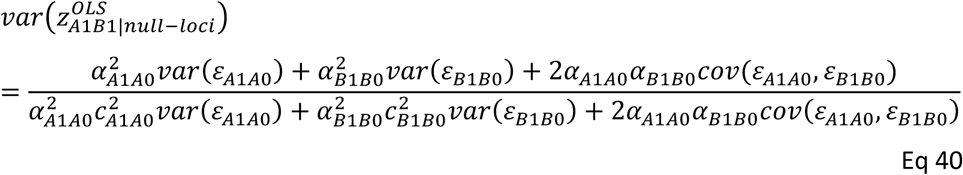

as confirmed with simulation (Table S18); note that for unbiased (*c*_*A*1*A*0_ = *c*_*B*1*B*0_ = 1) case-control GWAS results 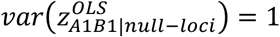 as expected. In another set of secondary analyses, CC-GWAS of SCZ vs. MDD was extended to include the direct case-case GWAS from ref.^4^ including 23,585 SCZ cases and 15,270 BIP cases (see URL section).

### The assumed number of causal SNPs in applications of CC-GWAS

Our primary recommendation is to specify *m* based on published estimates of genome-wide polygenicity, such as the effective number of independently associated causal SNPs^32^ or the total number of independently associated SNPs^31,70,88,89^. These values generally range from 1,000 for relatively sparse traits (e.g. autoimmune diseases) to 10,000 for highly polygenic traits (e.g. psychiatric disorders). When estimates of genome-wide polygenicity are not available, our recommendation is to specify *m*=1,000 for traits that are expected to have relatively sparse architectures (e.g. autoimmune diseases), *m*=10,000 for traits that are expected to have highly polygenic architectures (e.g. psychiatric disorders), and *m*=5,000 for traits with no clear expectation. When comparing disorders with different levels of polygenicity, our recommendation is to specify *m* based on the expected average across both disorders.

### SMR and HEIDI analyses

We used the SMR test for colocalization^35^ to identify CC-GWAS loci with significant associations between gene expression effect sizes in *cis* and CC-GWAS_OLS_ case-case effect sizes. We tested cis-eQTL effects in 13 GTEx v7 brain tissues^41^ (Amygdala, Anterior cingulate cortex, Caudate basal ganglia, Cerebellar Hemisphere, Cerebellum, Cortex, Frontal Cortex, Hippocampus, Hypothalamus, Nucleus accumbens basal ganglia, Putamen basal ganglia, Spinal cord cervical c-1, and Substantia nigra; see URLs), and a meta-analysis of eQTL effects in brain tissues^42^ (see URLs). In line with standard application of SMR^35^, we tested probes of genes with significant eQTL associations, with the lead eQTL SNP within 1MB of the lead CC-GWAS SNP. SMR analyses were performed on 2MB cis windows around the tested probe. The threshold of significance was adjusted per tested disorder-pair by dividing 0.05 by the respective number of probes tested (Table S23). We used the HEIDI test for heterogeneity^35^ to exclude loci with evidence of linkage effects (2 < 0.05).

### Replication data sets

For replication analyses, we used CC-GWAS discovery results of additional analyses of SCZ vs. MDD based on GWAS results from Ripke et al.^14^ and Wray et al.^61^ (Table S25; see URLs). To obtain independent replication data, we applied MetaSubtract^62^ separately for SCZ (results (i) Pardinas et al.^15^ – results (ii) Ripke et al.^14^) and for MDD (results (i) Howard et al.^17^ – results (ii) Wray et al.^61^). MetaSubtract^62^ recovers these results by subtracting results (ii) from results (i) as follows:

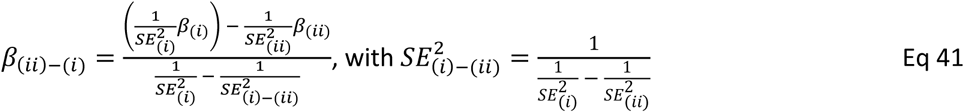

Replication GWAS results were thus obtained for 7,035 cases and 21,187 controls (*N*_*eff*_ = 22,369) for SCZ and 111,175 cases and 216,289 controls (*N*_*eff*_ = 242,275) for MDD, with information on 4,105,296 SNPs for SCZ and MDD (see Table S25 for sample sizes in discovery analyses). Independence of replication results and discovery results were confirmed with cross-trait LD score regression intercepts of 0.037 for SCZ and −0.004 for MDD. We corrected the replication results for the univariate LD score regression intercepts (by multiplying SE with 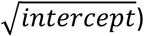), as the intercepts appeared slightly inflated (1.25 (attenuation ratio 0.58) for SCZ and 1.12 (attenuation ratio 0.25) for MDD). We note this correction may have been overly conservative, because if the discovery data are not exact subsets of the full data, we anticipate that MetaSubtract replication results would be conservative, as independent signals from the discovery data would be subtracted from the full data when producing the replication data.

For further replication analyses, we used CC-GWAS discovery results from the three comparisons of CD^63^, UC^63^ and RA^64^ (Table S25). For CD^63^ and UC^63^, two sets of GWAS results are publicly available (see URLs): (a) results of 1000 genomes Phase 1 imputed SNPs from individuals genotyped on genotyping chips with genome-wide coverage, and (b) meta-analyses results of (a) with additional samples genotyped with the Immunochip covering 196,524 SNPs without genome-wide coverage^63^. Our CC-GWAS discovery results were based on GWAS results (a). For replication, we needed the Immunochip only results, which we obtained by subtracting (a) from (b) with MetaSubtract^62^. These Immonochip only results were thus obtained for 101,482 SNPs after QC for CD and 101,792 SNPs for UC, and were based on 14,594 cases and 26,715 controls (*N*_*eff*_ = 37,752) for CD and 10,679 cases and 26,715 controls (*N*_*eff*_ = 30,517) for UC. For RA, the genome-wide GWAS results and Immunochip results are separately available (see URLs) providing replication results of 830,956 SNPs (note that Okada et al.^64^ imputed Immunochip SNP data) of 5,486 cases and 14,556 controls (*N*_*eff*_= 15,274). (Note that LD score regression could not be applied on the Immunochip results from CD, UC and RA, because the Immunochip does not provide the genome-wide coverage required for LD score regression^37^. Further note that scaling of the SE based on the univariate LD score regression intercept only impacts the number of significant replicated loci, but not the regression slope in Figure 5.) All sample sizes for discovery and replication analyses are reported in Table S25.

In a final set of replication analyses, we sought to perform replication analyses using independent replication data that was obtained without requiring the use of MetaSubtract, and compared RA vs BC (of low biological interest, but useful for assessing the robustness of the CC-GWAS method). For the CC-GWAS discovery analysis, we compared RA GWAS (8,875 cases + 29,367 controls) vs. a meta-analysis of BC OncoArray sample^30^ (61,282 cases + 45,494 controls) and BC iCOGs sample^30^ (46,785 cases + 42,892 controls). For the CC-GWAS replication analysis, we compared RA immunochip (5,486 cases + 14,556 controls) vs. BC GWAS sample^30^ (14,910 cases + 17,588 controls).

### Application of CC-GWAS to replication data sets

For replication of SCZ vs. MDD, CD vs. UC, CD vs. RA and UC vs. RA we computed CC-GWAS_OLS_ *A*1*B*1 effect based on the CC-GWAS_OLS_ weights from the respective discovery results (Table S25). (We applied CC-GWAS_OLS_ weights from the discovery analyses rather than re-estimating the CC-GWAS_OLS_ weights based on the replication GWAS results, because CC-GWAS_OLS_ weights are sample size dependent.) For SCZ vs. MDD, the covariance of the error terms was negligible (cross-trait LD score regression intercept of 0.012) while the covariance could not be estimated for the autoimmune disease pairs (as LD score regression cannot be applied on Immunochip data) and was thus set to 0. Because covariance of the error terms decreases the standard error of CC-GWAS_OLS_ estimates (Eq 21), setting the covariance to 0 may lead to conservative bias with respect to significance in replication, but it does not impact the magnitude of the CC-GWAS_OLS_ effect size estimates themselves nor the regression slopes displayed in Figure 5.

## Acknowledgements

We thank A. Schoech, L. O’Connor, O. Weissbrod, S. Gazal, D. Ruderfer, B.W.J.H. Penninx, N.R. Wray, K. Kendler, J. Smoller, W. van Rheenen, and the members of the Cross-Disorder Group of the Psychiatric Genomics Consortium for helpful discussions. For information about sample-overlap, we thank S. Ripke, V. Trubetskoy and the PGC working groups of Schizophrenia, Bipolar Disorder, Major Depressive Disorder, Attention Deficit Hyperactivity Disorder, Eating Disorders, Autism Spectrum Disorder, and OCD & Tourette Syndrome. The breast cancer genome-wide association analyses were supported by the Government of Canada through Genome Canada and the Canadian Institutes of Health Research, the ‘Ministère de l’Économie, de la Science et de l’Innovation du Québec’ through Genome Québec and grant PSR-SIIRI-701, The National Institutes of Health (U19 CA148065, X01HG007492), Cancer Research UK (C1287/A10118, C1287/A16563, C1287/A10710) and The European Union (HEALTH-F2-2009-223175 and H2020 633784 and 634935). All studies and funders are listed in Michailidou et al (Nature, 2017). This research was funded by NIH grants R01 HG006399, R37 MH107649, R01 MH101244, R01 CA222147, and NWO Veni grant (91619152) to W.J.P. This research was conducted using the UK Biobank Resources under application 10438.

## Author contribution

W.J.P. and A.L.P. designed experiments, W.J.P. performed experiments and analysed data. W.J.P. and A.L.P. wrote the manuscript.

## Competing interests

The authors declare no competing interests.

## Notes

### Competing Interest Statement

The authors have declared no competing interest.

## References

1. Lee, S. H. et al. Genetic relationship between five psychiatric disorders estimated from genome-wide SNPs. Nat. Genet. 45, 984–94 (2013).

2. Bulik-Sullivan, B. et al. An atlas of genetic correlations across human diseases and traits. Nat. Genet. (2015). doi: 10.1038/ng.3406

3. Lee, P. H. et al. Genomic Relationships, Novel Loci, and Pleiotropic Mechanisms across Eight Psychiatric Disorders. Cell 179, 1469-1482.e11 (2019).

4. Ruderfer, D. M. et al. Genomic Dissection of Bipolar Disorder and Schizophrenia, Including 28 Subphenotypes. Cell 173, 1705-1715.e16 (2018).

5. Pasaniuc, B. & Price, A. L. Dissecting the genetics of complex traits using summary association statistics. Nat. Rev. Genet. 18, 117–127 (2017).

6. Turley, P. et al. Multi-trait analysis of genome-wide association summary statistics using MTAG. Nat. Genet. 50, 229–237 (2018).

7. Qi, G. & Chatterjee, N. Heritability informed power optimization (HIPO) leads to enhanced detection of genetic associations across multiple traits. PLoS Genet. 14, e1007549 (2018).

8. Baselmans, B. M. L. et al. Multivariate genome-wide analyses of the well-being spectrum. Nat. Genet. 51, 445–451 (2019).

9. Nieuwboer, H. A., Pool, R., Dolan, C. V., Boomsma, D. I. & Nivard, M. G. GWIS: Genome-Wide Inferred Statistics for Functions of Multiple Phenotypes. Am. J. Hum. Genet. 99, 917–927 (2016).

10. Bhattacharjee, S. et al. A subset-based approach improves power and interpretation for the combined analysis of genetic association studies of heterogeneous traits. Am. J. Hum. Genet. 90, 821–35 (2012).

11. Han, B. & Eskin, E. Interpreting meta-analyses of genome-wide association studies. PLoS Genet. 8, e1002555 (2012).

12. Zhu, Z. et al. Causal associations between risk factors and common diseases inferred from GWAS summary data. Nat. Commun. 9, 224 (2018).

13. Byrne, E. M. et al. Conditional GWAS analysis identifies putative disorder-specific SNPs for psychiatric disorders. bioRxiv 592899 (2019). doi: 10.1101/592899

14. Ripke, S. et al. Biological insights from 108 schizophrenia-associated genetic loci. Nature 511, 421–7 (2014).

15. Pardiñas, A. F. et al. Common schizophrenia alleles are enriched in mutation-intolerant genes and in regions under strong background selection. Nat. Genet. 50, 381–389 (2018).

16. Stahl, E. A. et al. Genome-wide association study identifies 30 loci associated with bipolar disorder. Nat. Genet. 51, 793–803 (2019).

17. Howard, D. M. et al. Genome-wide meta-analysis of depression identifies 102 independent variants and highlights the importance of the prefrontal brain regions. Nat. Neurosci. 22, 343–352 (2019).

18. Demontis, D. et al. Discovery of the first genome-wide significant risk loci for attention deficit/hyperactivity disorder. Nat. Genet. 51, 63–75 (2019).

19. Watson, H. J. et al. Genome-wide association study identifies eight risk loci and implicates metabo-psychiatric origins for anorexia nervosa. Nat. Genet. 51, 1207–1214 (2019).

20. Grove, J. et al. Identification of common genetic risk variants for autism spectrum disorder. Nat. Genet. 51, 431–444 (2019).

21. International Obsessive Compulsive Disorder Foundation Genetics Collaborative (IOCDF-GC) and OCD Collaborative Genetics Association Studies (OCGAS). Revealing the complex genetic architecture of obsessive-compulsive disorder using meta-analysis. Mol. Psychiatry 23, 1181–1188 (2018).

22. Yu, D. et al. Interrogating the Genetic Determinants of Tourette’s Syndrome and Other Tic Disorders Through Genome-Wide Association Studies. Am. J. Psychiatry 176, 217–227 (2019).

23. Yang, J., Wray, N. R. & Visscher, P. M. Comparing apples and oranges: equating the power of case-control and quantitative trait association studies. Genet. Epidemiol. 34, 254–7 (2010).

24. Lee, S. H., Wray, N. R., Goddard, M. E. & Visscher, P. M. Estimating Missing Heritability for Disease from Genome-wide Association Studies. Am. J. Hum. Genet. 88, 294–305 (2011).

25. Pe’er, I., Yelensky, R., Altshuler, D. & Daly, M. J. Estimation of the multiple testing burden for genomewide association studies of nearly all common variants. Genet. Epidemiol. 32, 381–385 (2008).

26. Gelman, A. & Tuerlinckx, F. Type S error rates for classical and Bayesian single and multiple comparison procedures. Comput. Stat. 15, 373–390 (2000).

27. Stephens, M. False discovery rates: a new deal. Biostatistics 18, 275–294 (2017).

28. Reich, D. E. et al. Linkage disequilibrium in the human genome. Nature (2001). doi: 10.1038/35075590

29. Slatkin, M. Linkage disequilibrium - Understanding the evolutionary past and mapping the medical future. Nature Reviews Genetics (2008). doi: 10.1038/nrg2361

30. Michailidou, K. et al. Association analysis identifies 65 new breast cancer risk loci. Nature 551, 92–94 (2017).

31. Zhang, Y. D. et al. Assessment of polygenic architecture and risk prediction based on common variants across fourteen cancers. Nat. Commun. (2020).

32. O’Connor, L. J. et al. Extreme Polygenicity of Complex Traits Is Explained by Negative Selection. Am. J. Hum. Genet. 105, 456–476 (2019).

33. Genomes Project Consortium et al. A global reference for human genetic variation. Nature 526, 68–74 (2015).

34. Buniello, A. et al. The NHGRI-EBI GWAS Catalog of published genome-wide association studies, targeted arrays and summary statistics 2019. Nucleic Acids Res. 47, D1005–D1012 (2019).

35. Zhu, Z. et al. Integration of summary data from GWAS and eQTL studies predicts complex trait gene targets. Nat. Genet. 48, (2016).

36. Bulik-Sullivan, B. K. et al. LD Score regression distinguishes confounding from polygenicity in genome-wide association studies. Nat. Genet. 47, 291–5 (2015).

37. Finucane, H. K. et al. Partitioning heritability by functional annotation using genome-wide association summary statistics. Nat. Genet. (2015). doi: 10.1038/ng.3404

38. Gazal, S. et al. Linkage disequilibrium–dependent architecture of human complex traits shows action of negative selection. Nat. Genet. 49, 1421–1427 (2017).

39. Gazal, S., Marquez-Luna, C., Finucane, H. K. & Price, A. L. Reconciling S-LDSC and LDAK functional enrichment estimates. Nat. Genet. 51, 1202–1204 (2019).

40. Loh, P.-R., Kichaev, G., Gazal, S., Schoech, A. P. & Price, A. L. Mixed-model association for biobank-scale datasets. Nat. Genet. 50, 906–908 (2018).

41. GTEx Consortium et al. Genetic effects on gene expression across human tissues. Nature 550, 204–213 (2017).

42. Qi, T. et al. Identifying gene targets for brain-related traits using transcriptomic and methylomic data from blood. Nat. Commun. 9, 2282 (2018).

43. O’Leary, N. A. et al. Reference sequence (RefSeq) database at NCBI: current status, taxonomic expansion, and functional annotation. Nucleic Acids Res. 44, D733–45 (2016).

44. Hoefs, S. J. G. et al. NDUFA2 complex I mutation leads to Leigh disease. Am. J. Hum. Genet. 82, 1306–15 (2008).

45. Li, Z. et al. Genome-wide association analysis identifies 30 new susceptibility loci for schizophrenia. Nat. Genet. 49, 1576–1583 (2017).

46. Ikeda, M. et al. Genome-Wide Association Study Detected Novel Susceptibility Genes for Schizophrenia and Shared Trans-Populations/Diseases Genetic Effect. Schizophr. Bull. 45, 824–834 (2019).

47. Lam, M. et al. Comparative genetic architectures of schizophrenia in East Asian and European populations. Nat. Genet. 51, 1670–1678 (2019).

48. Lam, M. et al. Pleiotropic Meta-Analysis of Cognition, Education, and Schizophrenia Differentiates Roles of Early Neurodevelopmental and Adult Synaptic Pathways. Am. J. Hum. Genet. 105, 334–350 (2019).

49. Nagel, M. et al. Meta-analysis of genome-wide association studies for neuroticism in 449,484 individuals identifies novel genetic loci and pathways. Nat. Genet. 50, 920–927 (2018).

50. Lee, J. J. et al. Gene discovery and polygenic prediction from a genome-wide association study of educational attainment in 1.1 million individuals. Nat. Genet. 50, 1112–1121 (2018).

51. Savage, J. E. et al. Genome-wide association meta-analysis in 269,867 individuals identifies new genetic and functional links to intelligence. Nat. Genet. 50, 912–919 (2018).

52. Giri, A. et al. Trans-ethnic association study of blood pressure determinants in over 750,000 individuals. Nat. Genet. 51, 51–62 (2019).

53. Evangelou, E. et al. Genetic analysis of over 1 million people identifies 535 new loci associated with blood pressure traits. Nat. Genet. 50, 1412–1425 (2018).

54. The UniProt Consortium. UniProt: the universal protein knowledgebase. Nucleic Acids Res. 45, D158–D169 (2017).

55. Okbay, A. et al. Genome-wide association study identifies 74 loci associated with educational attainment. Nature 533, 539–542 (2016).

56. Hill, W. D. et al. A combined analysis of genetically correlated traits identifies 187 loci and a role for neurogenesis and myelination in intelligence. Mol. Psychiatry 24, 169–181 (2019).

57. Davies, G. et al. Study of 300,486 individuals identifies 148 independent genetic loci influencing general cognitive function. Nat. Commun. 9, 2098 (2018).

58. Moore, D. L., Apara, A. & Goldberg, J. L. Krüppel-like transcription factors in the nervous system: novel players in neurite outgrowth and axon regeneration. Mol. Cell. Neurosci. 47, 233–43 (2011).

59. Sekar, A. et al. Schizophrenia risk from complex variation of complement component 4. Nature 530, 177–83 (2016).

60. Yanagi, M. et al. Expression of Kruppel-like factor 5 gene in human brain and association of the gene with the susceptibility to schizophrenia. Schizophr. Res. 100, 291–301 (2008).

61. Wray, N. R. et al. Genome-wide association analyses identify 44 risk variants and refine the genetic architecture of major depression. Nat. Genet. 50, 668–681 (2018).

62. Nolte, I. M. et al. Missing heritability: is the gap closing? An analysis of 32 complex traits in the Lifelines Cohort Study. Eur. J. Hum. Genet. 25, 877–885 (2017).

63. Liu, J. Z. et al. Association analyses identify 38 susceptibility loci for inflammatory bowel disease and highlight shared genetic risk across populations. Nat. Genet. 47, 979–986 (2015).

64. Okada, Y. et al. Genetics of rheumatoid arthritis contributes to biology and drug discovery. Nature 506, 376–81 (2014).

65. Marigorta, U. M. & Navarro, A. High trans-ethnic replicability of GWAS results implies common causal variants. PLoS Genet. 9, e1003566 (2013).

66. Palmer, C. & Pe’er, I. Statistical correction of the Winner’s Curse explains replication variability in quantitative trait genome-wide association studies. PLOS Genet. 13, e1006916 (2017).

67. Benjamini, Y. & Hochberg, Y. Controlling the False Discovery Rate: A Practical and Powerful Approach to Multiple Testing. J. R. Stat. Soc. Ser. B 57, 289–300 (1995).

68. Nelson, C. P. et al. Association analyses based on false discovery rate implicate new loci for coronary artery disease. Nat. Genet. 49, 1385–1391 (2017).

69. Kanai, M. et al. Genetic analysis of quantitative traits in the Japanese population links cell types to complex human diseases. Nat. Genet. 50, 390–400 (2018).

70. Frei, O. et al. Bivariate causal mixture model quantifies polygenic overlap between complex traits beyond genetic correlation. Nat. Commun. 10, 2417 (2019).

71. Knevel, R. et al. Using genetics to prioritize diagnoses for rheumatology outpatients with inflammatory arthritis. Sci. Transl. Med. 12, eaay1548 (2020).

72. Traglia, M. et al. Genetic Mechanisms Leading to Sex Differences Across Common Diseases and Anthropometric Traits. Genetics 205, 979–992 (2017).

73. Khramtsova, E. A., Davis, L. K. & Stranger, B. E. The role of sex in the genomics of human complex traits. Nat. Rev. Genet. 20, 173–190 (2019).

74. Aschard, H., Vilhjálmsson, B. J., Joshi, A. D., Price, A. L. & Kraft, P. Adjusting for Heritable Covariates Can Bias Effect Estimates in Genome-Wide Association Studies. Am. J. Hum. Genet. 96, 329–339 (2015).

75. Bhatia, G., Patterson, N., Sankararaman, S. & Price, A. L. Estimating and interpreting FST: The impact of rare variants. Genome Res. 23, 1514–1521 (2013).

76. Price, A. L. et al. Discerning the ancestry of European Americans in genetic association studies. PLoS Genet. 4, e236 (2008).

77. Milne, R. L. et al. Identification of ten variants associated with risk of estrogen-receptor-negative breast cancer. Nat. Genet. 49, 1767–1778 (2017).

78. Golan, D., Lander, E. S. & Rosset, S. Measuring missing heritability: Inferring the contribution of common variants. Proc. Natl. Acad. Sci. U. S. A. 111, E5272–81 (2014).

79. Lloyd-Jones, L. R., Robinson, M. R., Yang, J. & Visscher, P. M. Transformation of Summary Statistics from Linear Mixed Model Association on All-or-None Traits to Odds Ratio. Genetics 208, 1397–1408 (2018).

80. Chang, C. C. et al. Second-generation PLINK: rising to the challenge of larger and richer datasets. Gigascience 4, 7 (2015).

81. van Rheenen, W., Peyrot, W. J., Schork, A. J., Lee, S. H. & Wray, N. R. Genetic correlations of polygenic disease traits: from theory to practice. Nat. Rev. Genet. 20, 567–581 (2019).

82. Conneely, K. N. & Boehnke, M. So many correlated tests, so little time! Rapid adjustment of P values for multiple correlated tests. Am. J. Hum. Genet. (2007). doi: 10.1086/522036

83. Siegel, R., Ma, J., Zou, Z. & Jemal, A. Cancer statistics, 2014. CA. Cancer J. Clin. 64, 9–29 (2014).

84. Sullivan, P. F. & Geschwind, D. H. Defining the Genetic, Genomic, Cellular, and Diagnostic Architectures of Psychiatric Disorders. Cell 177, 162–183 (2019).

85. Molodecky, N. A. et al. Increasing incidence and prevalence of the inflammatory bowel diseases with time, based on systematic review. Gastroenterology 142, 46-54.e42; quiz e30 (2012).

86. Yengo, L., Yang, J. & Visscher, P. M. Expectation of the intercept from bivariate LD score regression in the presence of population stratification. bioRxiv 310565 (2018). doi: 10.1101/310565

87. R Core Team. R: A Language and Environment for Statistical Computing. (2018).

88. Zhang, Y., Qi, G., Park, J.-H. & Chatterjee, N. Estimation of complex effect-size distributions using summary-level statistics from genome-wide association studies across 32 complex traits. Nat. Genet. 50, 1318–1326 (2018).

89. Zeng, J. et al. Signatures of negative selection in the genetic architecture of human complex traits. Nat. Genet. 50, 746–753 (2018).

