## Supplementary Information for "Identifying loci with different allele frequencies among cases of eight psychiatric disorders using CC-GWAS"

### Supplementary Note

#### *1. Filtering step to exclude false positive associations due to differential tagging of a causal stress test SNP*

(This section repeats content from the Methods section, to ensure that it is self-contained.) CC-GWAS identifies and discards false positive associations that can arise due to differential tagging of a causal stress test SNP (with the same allele frequency among cases of both disorders), e.g. due to subtle differences in ancestry between the input case-control studies (Figure 3A). Specifically, CC-GWAS screens the 1MB region around every genome-wide significant candidate CC-GWAS SNP for evidence of a differentially linked stress test SNP, and conservatively filters the candidate CC-GWAS SNP when suggestive evidence of a differentially linked stress test SNP is detected. The filtering criteria were motivated by extensive simulations (see below; Table S7). A list of the filtering criteria is provided in Table S6.

For each candidate CC-GWAS SNP, filtering comprises of three sets of criteria (A, B, and C), and the SNP is discarded when at least one of the three sets of criteria is met. The criteria (A) are intended for intermediate sample sizes, the criteria (B) for relatively small sample sizes, and the criteria (C) for very large sample sizes. In the first set of criteria (A), CC-GWAS considers, as a potential stress test SNP, the SNP in a 1Mb region (around the candidate CC-GWAS SNP) with the largest product of case-control z-scores across the two disorders, i.e. the SNP $max.zAzB$ with $zA_{max.zAzB} * zB_{max.zAzB} = \max_{1MB}\{zA * zB\}$ (a causal stress test will generally have the maximum $|zA|$ and maximum $|zB|$ value, at same sign of effect). The candidate CC-GWAS SNP is filtered as potential false positive association when *all* of the following criteria A1, A2 and A3 are met:

A1. The SNP $max.zAzB$ is likely to have the same population allele frequencies among cases of the two disorders, reflected by a CC-GWAS$_{Exact}$ p-value larger than $10^{-4}$.

A2. The SNP $max.zAzB$ is likely to be the causal SNP for both disorders, reflected by absolute case-control z-scores almost as large (allowing for sampling variance) as the largest absolute case-control z-scores in the region for both disorders:

- $|zA_{max.zAzB}| - |zA_{max.zA}| > -1$ ***AND*** $|zB_{max.zAzB}| - |zB_{max.zB}| > -1$, where $max.zA$ denotes the SNP with $|zA_{max.zA}| = max_{1MB}\{|zA|\}$, and $max.zB$ the SNP with $|zB_{max.zB}| = max_{1MB}\{|zB|\}$.

A3. The case-control z-scores at the candidate CC-GWAS SNP ($ccgwas$) and at SNP $max.zAzB$ have a pattern concordant with differential tagging.

- $|(zA_{ccgwas}/zA_{max.zAzB}) - (zB_{ccgwas}/zB_{max.zAzB})| < 1$

Note that these ratios of z-scores roughly correspond to the correlation (LD) between the SNP $ccgwas$ and SNP $max.zAzB$ in the respective populations (if SNP $max.zAzB$ is indeed the causal SNP in the region). If this criterion is not met, the case-control z-scores at SNP $ccgwas$ and at SNP $max.zAzB$ have a pattern that is *not* concordant with differential tagging. The threshold of 1 accounts for differences in LD as well as sampling variance; a difference larger than 1 would require other explanations (such as e.g. SNP $max.zAzB$ is not the causal SNP in the region, or SNP $max.zAzB$ is a causal SNP independent from SNP $ccgwas$).

The second set of criteria (B) is intended for when at least one case-control GWAS is underpowered, in which case the most likely potential stress test SNP is no longer the SNP $max.zAzB$, due to sampling variance. Specifically, if the case-control GWAS of at least one disorder is underpowered ($N_{eff} < 40k$), CC-GWAS additionally considers, as a potential stress test SNP, the SNP in a 1Mb region with the largest maximum absolute case-control z-score across the two disorders, i.e. the SNP $max.z$ with $max\{|zA_{max.z}|, |zB_{max.z}|\} = max\{\max_{1MB}\{|zA|\}, \max_{1MB}\{|zB|\}\}$ (as this is the most likely causal SNP in this region). The CC-GWAS SNP is filtered as a potential false positive association when the following criterium B1 is met:

B1. The SNP $max.z$ is likely to have the same population allele frequencies among cases, reflected by a CC-GWAS$_{\text{Exact}}$ p-value larger than $10^{-4}$. (We note that we do not include criteria analogous to A2 and A3, because the error around the z-scores is too large. We also note that when the causal stress test SNP has small effects, GWAS samples with $N_{eff} \geq 40k$ may also be underpowered; however, the per-locus type I error rate is already $< 10^{-4}$ before applying the CC-GWAS filtering step in this scenario; see Table S7.)

The third set of criteria (C) is primarily intended for when the case-control GWAS are very well powered and the causal stress test SNP is *not* genotyped/imputed. In this case, very subtle tagging differences between the potential stress test SNP in criteria (A) and the causal stress test SNP can lead to $p_{Exact} < 10^{-4}$ (i.e. violation of A1, thus wrongfully not excluding the candidate CC-GWAS SNP). Therefore, the CC-GWAS SNP is filtered when the following criterium C1 is met (irrespective of sample size):

C1. The power of CC-GWAS is much lower than the power of the respective case-control analyses, reflected by a much smaller CC-GWAS$_{Exact}$ z-score of SNP $ccgwas$ than the corresponding case-control z-scores.

- $\left|z_{ccgwas}^{Exact}\right| < 0.5 * |zA_{max.zA}|$ ***AND*** $\left|z_{ccgwas}^{Exact}\right| < 0.5 * |zB_{max.zB}|$

The candidate CC-GWAS SNP is excluded when at least one of the criteria (A), (B) and (C) is met.

#### *2. Perturbing criteria in filtering step to exclude false positive associations due to differential tagging of a causal stress test SNP*

The filtering criteria were designed to adequately control type I error due to differential tagging of a causal stress test SNP, while minimizing unnecessary filtering of true positive associations. However, the criteria and thresholds applied are ad hoc and somewhat arbitrary. Therefore, in analyses of both simulated and empirical data, we conducted secondary analyses to assess the impact of perturbing each of the filtering criteria. Table S8 reports results of perturbation on filtering in simulated data, empirical analyses of breast cancer (BC vs. BC), and empirical analyses of psychiatric disorders. We note that, for each of the criteria, a more stringent threshold filters *fewer* loci, potentially resulting in *higher* power and/or *higher* type I error. For the simulated data we report the type I error, and for the empirical data we report the number of filtered loci.

Criterium (A1) is intended to check if the potential stress test SNP is likely to have the same allele frequency among cases (reflected by a CC-GWAS$_{Exact}$ p-value larger than $10^{-4}$). When varying the threshold in criterium (A1) from $10^{-4}$ to [$10^{-3}$,$10^{-7}$], type I error in simulations was generally well controlled but slightly increased for the more stringent threshold $10^{-3}$, empirical BC vs. BC analyses did not change (2/2 were filtered), and empirical analyses of psychiatric disorders changed little from 7/321 loci (for default threshold of $10^{-4}$) to 7/321 loci (for threshold of $10^{-3}$) up to 14/321 loci (for threshold of $10^{-7}$). We believe a threshold of $10^{-4}$ is adequate, because stress test SNPs cannot be numerous (see Discussion in main manuscript). Note that this threshold is also in line with the threshold applied in the CC-GWAS$_{Exact}$ component of CC-GWAS (which aims to protect against type I error at the stress test SNPs themselves, as opposed to SNPs that differentially tag them). Criterium (A2) is intended to check if the potential stress test SNP is likely to be the causal SNP in the region for both disorders (reflected by having a z-score at most −1 smaller than the maximum case-control z-score for both disorders). When varying the threshold in criterium (A2) from −1 to [−0.5,−3], type I error in simulations was well controlled throughout, empirical BC vs. BC analyses did not change (2/2 were filtered), and empirical analyses of psychiatric disorders changed little from 7/321 loci (for default threshold of −1) to 7/321 loci (for threshold of −0.5) up to 11/321 loci (for threshold of −3).

Threshold (A3) is intended to check that the candidate CC-GWAS SNP and the potential stress test SNP have case-control z-scores with a pattern concordant with differential tagging (see above and Table S8 for details).

When varying the threshold in criterium (A3) from 1 to [0.5,2], type I error in simulations was generally well controlled but slightly increased for the more stringent threshold of 0.5, empirical BC vs. BC analyses did not change (2/2 were filtered), and empirical analyses of psychiatric disorders changed little from 7/321 loci (for default threshold of 1) to 7/321 loci (for threshold of 0.5) up to 12/321 loci (for threshold of 2).

Criterium (B1) is intended for when at least one case-control GWAS is underpowered ($N_{eff}$ < 40k). When varying the $N_{eff}$ threshold underlying criterium (B1) from 40k to [5k,80k], type I error in simulations increased to $1.2x10^{-3}$ and $4.4x10^{-4}$ (for threshold 5k) for parameters in line with SCZ vs. OCD resp. SCZ vs. TS (the two pairs of psychiatric disorders from which candidate CC-GWAS loci were filtered in empirical analyses based on criterium (B), with 5k<$N_{eff}$<40k for one of the disorders) and was well controlled for the other thresholds, empirical BC vs. BC analyses did not change (2/2 were filtered), and empirical analyses of psychiatric disorders changed modestly from 7/321 loci (for default threshold of 40k) to 0/321 loci (for threshold of 5k) up to 35/321 loci (for threshold of 80k). These simulation results illustrate the necessity to include criterium (B1) in our filtering, because a threshold 5k is equivalent to excluding criterium (B1) for the simulated parameter settings. Furthermore, extensive simulations for a wide range of $N_{eff}$ (for default threshold of 40k) suggests that the threshold of 40k adequately controls type I error (Table S7), and a threshold of 80k may thus be overly conservative. (Note that when the causal stress test SNP has small effects, GWAS samples with $N_{eff} \geq 40k$ may also be underpowered; however, the per-locus type I error rate is already $< 10^{-4}$ before applying the CC-GWAS filtering step in this scenario; Table S7.) Criterium (B1) is intended to check if the potential stress test SNP is likely to have the same allele frequency among cases (reflected by a CC-$GWAS_{Exact}$ p-value larger than $10^{-4}$). When varying the p-value threshold in criterium (B1) from $10^{-4}$ to [$10^{-5}$,$10^{-3}$], type I error in simulations increased to $1.7x10^{-4}$ and $9.9x10^{-5}$ (for threshold $10^{-3}$) for parameters in line with SCZ vs. OCD resp. SCZ vs. TS (the two pairs of psychiatric disorders from which candidate CC-GWAS loci were filtered in empirical analyses based on criterium (B)) and was well controlled for the other thresholds, empirical BC vs. BC analyses did not change (2/2 were filtered), and empirical analyses of psychiatric disorders changed modestly from 7/321 loci (for default threshold of $10^{-4}$) to 16/321 loci (for threshold of $10^{-5}$) down to 3/321 loci (for threshold of $10^{-3}$). We believe a threshold of $10^{-4}$ is adequate, because stress test SNPs cannot be numerous (see Discussion in main manuscript).

Criterium (C1) is primarily intended for when the case-control GWAS are very well powered and the causal stress test SNP is *not* genotyped/imputed. When varying the z-score ratio threshold in criterium (C1) from 0.5 to [0.25,0.75], type I error in simulations was generally well controlled (although slightly increased (per-locus type I error of $3.6 \times 10^{-4}$) for threshold 0.25 at the simulation parameters of Figure 3 with 5x sample size multiplier with the causal stress test SNP untyped), empirical BC vs. BC analyses did not change (2/2 were filtered), and empirical analyses of psychiatric disorders changed little from 7/321 loci (for default threshold of 0.5) to 7/321 loci (for threshold of 0.25) up to 8/321 loci (for threshold of 0.75).

Finally, we investigated the impact of applying criteria (A), (B) or (C) individually (with default thresholds), instead of applying all of the criteria. When applying only criteria (A), type I error in simulations increased to $1.2 \times 10^{-3}$ and $4.4 \times 10^{-4}$ for parameters in line with SCZ vs. OCD resp. SCZ vs. TS (the two pairs of psychiatric disorders from which candidate CC-GWAS loci were filtered in empirical analyses based on criterium (B)) and type I error in simulations increased for simulation parameters of Figure 3 when the causal stress test SNP is *not* genotyped/imputed (in particular for 5x sample size multiplier, due to the absence of criterium (C)), empirical BC vs. BC analyses did not change (2/2 were filtered), and none of the loci in the empirical analyses of psychiatric disorders loci (0/321) were filtered. Thus, criteria (A) did not provide appropriate type I error control when applied without criteria (B) and (C). When applying only criterium (B), type I error in simulations increased for most simulation parameters (as most disorder pairs had $N_{eff}$>40k for both disorders), none of the empirical BC vs. BC loci (0/2) were filtered (as sample-sizes were >40k), and 7 of the loci in the empirical analyses of psychiatric disorders (7/321) were filtered. Thus, criterium (B) did not provide appropriate type I error control when applied without criteria (A) and (C). When applying only criterium (C), type I error in simulations increased to $1.2 \times 10^{-3}$ and $4.4 \times 10^{-4}$ for parameters in line with SCZ vs. OCD resp. SCZ vs. TS (the two pairs of psychiatric disorders from which candidate CC-GWAS loci were filtered in empirical analyses based on criterium (B)), both empirical BC vs. BC loci (2/2) were filtered, and none of the loci in the empirical analyses of psychiatric disorders loci (0/321) were filtered. Thus, criterium (C) did not provide appropriate type I error control when applied without criteria (A) and (B). We conclude that alle of the criteria (A), (B) or (C) are necessary to adequately control type I error due to differential tagging of a causal stress test SNP for a wide range of possible disorder pairs.

#### *3. Comparison of CC-GWAS vs. simple meta-analysis based method*

An alternative approach for identifying SNPs with disorder-specific effects is to identify all SNPs that have a genome-wide significant effect in at least one of the two disorders and do not have a genome-wide significant effect in a meta-analysis of the two disorders. We assessed two different versions of

Method MA: Method MA1, which outputs SNPs with p<5x10$^{-8}$ for one or both disorders and meta-analysis of both disorder GWAS p≥5x10$^{-8}$; and Method MA2, which outputs SNPs with p<2.5x10$^{-8}$ for one or both disorders and meta-analysis of both disorder GWAS p≥2.5x10$^{-8}$. We compute power, type I error at null-null SNPs, and type I error at stress test SNPs analogue to our computations for CC-GWAS. More specifically, consider the multivariate normal distribution $f\left(x = z_{A1A0}, y = z_{B1B0}, v = z_{meta-analysis}\right)$, where $z_{meta-analysis}$ follows from a fixed-effect meta-analysis as $\left\{\omega_1\hat{\beta}_{A1A0} + \omega_2\hat{\beta}_{B1B0}\right\}/\sqrt{(\omega_1 SE_{A1A0})^2 + (\omega_2 SE_{B1B0})^2}$ assuming no sample-overlap between both case-control samples. While assuming effect sizes following a bivariate normal distribution, the elements of the variance-covariance matrix for the causal component of the z-scores, $\Sigma_{causal}$, equal

- $var(causal\ comp.of\ z_{A1A0}) = F_{ST,causal,A1A0}/SE_{A1A0}$ (see Eq 12)
- $var(causal\ comp.of\ z_{B1B0}) = F_{ST,causal,B1B0}/SE_{B1B0}$ (see Eq 12)
- $var\left(causal\ comp.of\ z_{meta-analysis}\right) = \left\{\omega_1 F_{ST,causal,A1A0} + \omega_2 F_{ST,causal,B1B0} + 2\omega_1\omega_2\frac{coh_{oA,oB}}{m}\right\}/\sqrt{(\omega_1 SE_{A1A0})^2 + (\omega_2 SE_{B1B0})^2}$ (see Eq 4 and Eq 13)
- $cov(causal\ comp.of\ z_{A1A0}, causal\ comp.of\ z_{B1B0}) = \left(\frac{coh_{oA,oB}}{m}\right)/(SE_{A1A0}SE_{B1B0})$
- $cov\left(causal\ comp.of\ z_{A1A0}, causal\ comp.of\ z_{meta-analysis}\right) = \left(\omega_1 F_{ST,causal,A1A0} + \omega_2\left(\frac{coh_{oA,oB}}{m}\right)\right)/\left(SE_{A1A0}\sqrt{(\omega_1 SE_{A1A0})^2 + (\omega_2 SE_{B1B0})^2}\right)$ (see Eq 13)
- $cov\left(causal\ comp.of\ z_{B1B0}, causal\ comp.of\ z_{meta-analysis}\right) = \left(\omega_2 F_{ST,causal,B1B0} + \omega_1\left(\frac{coh_{oA,oB}}{m}\right)\right)/\left(SE_{B1B0}\sqrt{(\omega_1 SE_{A1A0})^2 + (\omega_2 SE_{B1B0})^2}\right)$ (see Eq 13)

The elements of the variance-covariance matrix for the error component of the z-scores, $\Sigma_{error}$, equal

- $var(error\ comp.of\ z_{A1A0}) = 1$
- $var(error\ comp.of\ z_{B1B0}) = 1$
- $var\left(error\ comp.of\ z_{meta-analysis}\right) = 1$
- $cov(error\ comp.of\ z_{A1A0}, error\ comp.of\ z_{B1B0}) = 0$ (assuming no sample-overlap)
- $cov\left(error\ comp.of\ z_{A1A0}, error\ comp.of\ z_{meta-analysis}\right) = \frac{(\omega_1/N_{eff,A1A0})}{SE_{A1A0}\sqrt{(\omega_1 SE_{A1A0})^2 + (\omega_2 SE_{B1B0})^2}}$
- $cov\left(error\ comp.of\ z_{B1B0}, error\ comp.of\ z_{meta-analysis}\right) = \frac{(\omega_2/N_{eff,B1B0})}{SE_{B1B0}\sqrt{(\omega_1 SE_{A1A0})^2 + (\omega_2 SE_{B1B0})^2}}$

The power of Method MA follows from function $f_{causal}$ with mean (0,0,0) and variance-covariance $\Sigma_{causal} + \Sigma_{error}$ as

$$\int_{-\infty}^{-z_{th}} \int_{-\infty}^{\infty} \int_{-z_{th}}^{z_{th}} f''' \, dxdydv + \int_{z_{th}}^{\infty} \int_{-\infty}^{\infty} \int_{-z_{th}}^{z_{th}} f''' \, dxdydv + \int_{-\infty}^{\infty} \int_{-\infty}^{-z_{th}} \int_{-z_{th}}^{z_{th}} f''' \, dxdydv + \int_{-\infty}^{\infty} \int_{z_{th}}^{\infty} \int_{-z_{th}}^{z_{th}} f''' \, dxdydv$$

where $z_{th}$ corresponds to the respective significance thresholds in Method MA1 and Methods MA2. The type I error at null-null SNPs of Method MA follows by substituting $f_{causal}$ with $f_{null-null}$ with mean (0,0,0) and variance-covariance $\Sigma_{error}$. The type I error at stress test SNPs of Method MA follows by substituting $f_{causal}$ with $f_{stress\ test}$ with mean corresponding with the case-control effects of the stress test SNP specified (see Figure 2) and variance-covariance $\Sigma_{error}$. These analytical computations were confirmed with simulation of individual level data in line with Table S2 (data not shown). Results of comparing CC-GWAS with Method MA1 and Methods MA2 are presented in Figure S17 and Figure S18 and their captions. Overall, we do not recommend the meta-analysis method, because it has lower power than CC-GWAS.

### Supplementary Table captions

**Table S1. Numerical results of power and type I error of CC-GWAS.** This Table reports the numerical results of Figure 2. We report (A) the power to detect SNPs with effect sizes following a bivariate normal distribution, (B) the type I error rate for loci with no effect on A1A0 or B1B0 ("null-null" SNPs) and (C) the type I error rate for SNPs with the same allele frequency in A1 vs. B1 that explain 0.10% of variance in A1 vs. A0 and 0.29% of variance in B1 vs. B0 ("stress test" SNPs), for each of four methods: CC-GWAS, the CC-GWAS$_{OLS}$ component, the CC-GWAS$_{Exact}$ component, and a naïve Delta method (see text). Default parameter settings are in line with Figure 2. These results are confirmed with simulation results in Table S2.

**Table S2. Power and type I error of CC-GWAS: analytical computations vs. simulation.** This Table accompanies Figure 2. We report analytical computations and simulation results of (A) the power to detect SNPs with effect sizes following a bivariate normal distribution, (B) the type I error rate for loci with no effect on A1A0 or B1B0 ("null-null" SNPs), (C) the type I error rate for SNPs (with MAF randomly drawn from [0.01-0.5]) with the same allele frequency in A1 vs. B1 that explain 0.10% of variance in A1 vs. A0 and 0.29% of variance in B1 vs. B0 ("stress test" SNPs) and (D) the type I error rate for the "same stress" test SNPs with MAF fixed at 0.01 (with odds ratios of approximately 1.6 in disorder A and 1.8 in disorder B), for each of five methods: CC-GWAS, the CC-GWAS$_{OLS}$ component, the CC-GWAS$_{Exact}$ component, the Delta method, and CC-GWAS+ (see Methods). Default parameter settings are: $h^2$=0.2, prevalence $K$=0.01, and sample size 4,000 cases + 4,000 controls for disorder A; liability-scale $h^2$=0.1, prevalence $K$=0.15, and sample size 4,000 cases + 4,000 controls for disorder B; $m$=1,000 causal SNPs for each disorder; and genetic correlation $r_g$=0.5 between disorders. Levels of significance were set as $p < 0.01$ for the CC-GWAS$_{OLS}$ weights and $p < 0.05$ for the CC-GWAS$_{Exact}$ weights. We report the mean (standard error) of 50 simulation runs (see Methods). The simulations in Panel A, Panel B and Panel C match the corresponding analytical computations, thereby justifying the analytical computations used in Figure 2. Furthermore, the type I error at stress test SNPs with MAF=0.01 (Panel D) was similarly well controlled as the type I error at stress test SNPs with MAF ranging 0.01-0.5 (Panel C), and in line with the specified level for significance in the CC-GWAS$_{Exact}$ component. (We note that the type I error at null-null SNPs in Panel B is not impacted by low MAF, provided that the case-control z scores are unbiased.)

**Table S3. Type S error of CC-GWAS.** This Table is an extension of Figure 2 and Table S1, and reports the type S error, defined as the proportion of significantly identified loci (true positives) identified with the wrong sign, for each of four methods: CC-GWAS, the CC-GWAS$_{OLS}$ component, the CC-GWAS$_{Exact}$ component, and a naïve Delta method (see text). Default parameter settings are in line with Figure 2.

**Table S4. CC-GWAS+ vs. MTAG.** When GWAS results of a direct case-case comparison (A1B1) are available, MTAG[1] can be applied to increase power of case-case analyses with the case-control (A1A0 and B1B0) GWAS results as helper traits. This Table presents a comparison of CC-GWAS+ and MTAG based on simulation of individual level data as in Table S2. Note that the required levels of significance were set as $p < 0.01$ for the CC-GWAS+$_{OLS}$ component (and for MTAG) and $p < 0.05$ for the CC-GWAS+$_{Exact}$ component. We reached five conclusions. First, although the magnitudes of CC-GWAS+$_{OLS}$ weights and MTAG weights are different (Panel A), they are roughly proportional (Panel B). This is because both CC-GWAS+$_{OLS}$ and MTAG aim to minimize the difference between estimated and true case-case effect sizes. Second, the power of CC-GWAS+$_{OLS}$ is slightly larger than the power of MTAG

(Panel C). This is because standard errors are defined differently. CC-GWAS+ defines its standard error as $\sqrt{var\big((\omega_1\beta_{A1A0}+\omega_2\beta_{B1B0}+\omega_3\beta_{A1B1})-(\omega_1\hat{\varepsilon}_{A1A0}+\omega_2\hat{\varepsilon}_{B1B0}+\omega_3\hat{\varepsilon}_{A1B1})\big)}$ (see Methods), which is the sampling variance of the estimator. On the other hand, MTAG defines its standard error as $\sqrt{var\left(\beta_{A1B1}-\big(\omega_1\hat{\beta}_{A1A0}+\omega_2\hat{\beta}_{B1B0}+\omega_3\hat{\beta}_{A1B1}\big)\right)}$ (Equation at left bottom of first page of Methods in ref.[1]). Thus, MTAG additionally models the variance of $\beta_{A1B1}-(\omega_1\beta_{A1A0}+\omega_2\beta_{B1B0}+\omega_3\beta_{A1B1})$ resulting in a larger standard error. Third, the power of CC-GWAS+ (combining the CC-GWAS+$_{OLS}$ component and the CC-GWAS+$_{Exact}$ component) is comparable to the power of MTAG (Panel C). Fourth, in line with the above, the type I error of MTAG is more conservative (<0.007) at null-null SNPs than the already adequate type I error of CC-GWAS+$_{OLS}$ (<0.01; Panel D). Fifth, and most importantly, the type I error at stress test SNP is much larger for MTAG (>0.1) than for CC-GWAS+ (<0.05; Panel E). In conclusion, when GWAS results of a direct case-case comparison are available, we advise the use of CC-GWAS+ over MTAG.

**Table S5. Applying CC-GWAS on data from a different bivariate genetic architecture.** The CC-GWAS+$_{OLS}$ component assumes that SNP effects for both disorders follow a bivariate normal distribution. Here we present results of CC-GWAS applied on simulated data violating this assumption, with the distribution of SNP effects in line with the general distribution applied in Frei et al.[2]: 1/3 of causal SNPs have an impact on disorder A only, 1/3 of SNPs have an impact on disorder B only, and 1/3 of SNPs have an impact both disorder A and disorder B (the correlation of these SNP effects specify the genome-wide genetic correlation, as in Frei et al.[2]). All other simulation parameters are in line with the simulations in Table S2, and the reported input/theory values represent the analytically expected results (i.e. exactly in line with Table S2). The average power across all causal SNPs is remarkably similar to the analytically expected power, although some differences exist for the three subsets of causal SNPs. Note that this Table reports no type I error at null-null SNPs and no type I error at stress test SNPs, because these SNPs are not impacted by this different bivariate genetic architecture (and do not differ from results in Table S2).

**Table S6. Filtering criteria of false positive associations due to differential tagging of a causal stress test SNP.** This Table reports an overview of the filtering criteria to identify and discard false positive associations that can arise due to differential tagging of a causal stress test SNP (with the same allele frequency in cases of both disorders), e.g. due to subtle differences in ancestry between the input case-control studies. See the Supplementary Note for more details.

**Table S7. Simulation of false positive associations due to differential tagging of a causal stress test SNP.** This Table accompanies Figure 3B, and presents simulation results for 34 different disorder pairs (6 pairs displayed in Figure 3B, and 28 pairs based on real-life comparisons of 8 psychiatric disorder pairs; see Table S14 for the CC-GWAS$_{OLS}$ and CC-GWAS$_{Exact}$ weights) and three different causal stress test SNPs explaining respectively a proportion of $10^{-3}$, $10^{-4}$ and $10^{-5}$ of liability variance in disorder A. We report the per tagging SNP type I error rate and the per locus type I error rate (the number of loci with at least one genome-wide significant tagging SNP divided by the number of loci tested) for CC-GWAS (with filtering) in the scenario where the causal stress test SNP is genotyped/imputed; CC-GWAS (with filtering) in the scenario where the causal stress test SNP is *not* genotyped/imputed; CC-GWAS with no filtering. See the Method section for more details.

**Table S8. Perturbing filtering criteria of false positive associations due to differential tagging of a causal stress test SNP.** This Table reports the impact of perturbating the filtering criteria on (i) the type I error of simulated loci filtered for a stress test SNP explaining 0.1% of liability variance in A and a selection of relevant parameter settings. We present simulating results of parameters in line with Figure 3, the three comparisons of SCZ, BIP, MDD, and of SCZ vs. OCD and SCZ vs. TS (the two pairs of psychiatric disorders from which candidate CC-GWAS loci were filtered), (ii) the empirical loci filtered in the BC vs. BC analyses, and (iii) the empirical loci filtered among the 8 psychiatric disorders. We note that, for each of the criteria, a more stringent threshold filters *fewer* loci, potentially resulting in *higher* power and/or *higher* type I error. For the simulated data we report the type I error, and for the empirical data we report the number of filtered loci. See the Supplementary Note for a detailed description of the results.

**Table S9. CC-GWAS analyses of breast cancer vs. breast cancer.**
For the BC (OncoArray sample in ref.[3]) vs. BC (iCOGs sample in ref.[3]) analyses, we report the case-control sample sizes, #SNPs, the most likely prevalence ($K$)[4], liability-scale heritability estimated using stratified LD score regression[5–7] ($h^2$), genetic correlation estimated using cross-trait LD score regression[8] ($r_g$), CC-GWAS$_{\text{OLS}}$ weights (based on the most likely prevalences), number of independent genome-wide significant loci for each case-control comparison, number of independent genome-significant CC-GWAS loci, and number of independent genome-significant CC-GWAS loci that are CC-GWAS-specific. CC-GWAS$_{\text{Exact}}$ weights are equal to $(1 - K_A)$ for disorder A and − $(1 - K_B)$ for disorder B. Notably, CC-GWAS identified two independent genome-wide significant candidate loci (containing a total of 5 SNPs) prior to filtering for differential tagging of stress test SNPs. All SNPs in these two loci very clearly met the filtering criteria (Table S10).

**Table S10. Empirical loci excluded by the filter of potential false positive associations due to differential tagging of a causal stress test SNP.** This Table reports details of the empirical candidate loci that were filtered from the analyses of breast cancer vs. breast cancer, and psychiatric disorders. We report the specific criteria and values on the basis of which these loci were excluded. Although we report only the candidate CC-GWAS lead SNP here, some candidate loci contained more than 1 candidate SNP; these candidate SNPs were all excluded based on the same criteria as the respective lead SNP.

**Tables S11. Genetic distance between cases and/or controls of eight psychiatric disorders and three autoimmune diseases.** This Table reports the numerical results of Figure 1 and Figure S11. We report $F_{ST,causal}$ for eight psychiatric disorders and three autoimmune diseases. The quantity $F_{st,causal}$ is derived based on the respective population prevalences, SNP-based heritabilities, genetic correlations, and number of assumed independent causal SNPs ($m$). The equation of $F_{st,causal}$ has $m$ in the denominator, thus $m * F_{st,causal}$ (reported in Figure 1 and Figure S11) is independent of $m$. SCZ, schizophrenia; BIP, bipolar disorder; MDD, major depressive disorder; ADHD, attention deficit/hyperactivity disorder; ANO, anorexia nervosa; ASD, autism spectrum disorder; OCD, obsessive–compulsive disorder; TS, Tourette's Syndrome and Other Tic Disorders; CD, Crohn's disorder; UC, ulcerative colitis; RA, rheumatoid arthritis.

**Table S12. Overview of definitions of independent loci used throughout the manuscript.** We report the two definitions of independent loci used throughout the manuscript for: (i) clumping of CC-GWAS results, (ii) comparing CC-GWAS loci to input case-control results to define CC-GWAS-specific loci, (iii) comparing CC-GWAS loci to CC-GWAS results from other disorder pairs, (iv) comparing CC-GWAS results to results from other studies.

**Table S13. List of 313 CC-GWAS loci for eight psychiatric disorders.** This Table is an extension of Table 2. For each CC-GWAS locus, we report the lead CC-GWAS SNP and its chromosome, physical position, and reference allele frequency, the locus name, the respective case-control effect sizes and *p*-values, the CC-GWAS$_{OLS}$ case-case effect size and *p*-value, the CC-GWAS$_{Exact}$ case-case effect size and *p*-value, indicator of CC-GWAS-specific loci, overlap with other CC-GWAS loci, indicator of independent loci (of the 28 pairs comparing all 8 psychiatric disorders, of the 3 pairs comparing SCZ, BIP and MDD, of the 10 pairs comparing SCZ, BIP, MDD, ADHD and ASD analyzed by Byrne et al.[9]), number of genes with significant SMR colocalization results (for CC-GWAS-specific loci only), overlap with Lee et al.[10], overlap with Byrne et al.[9], and overlap with results in the GWAS Catalog[11] displayed as "first author, year (PMID)". Effect sizes are reported on the standardized observed scale based on 50/50 case-control ascertainment. [a]denotes loci that have not been reported previously[11]. [b]denotes locus names based on (most) significant SMR results (only applicable for CC-GWAS specific loci). [c]denotes locus names based on exonic lead SNPs (applicable for all CC-GWAS specific). Remaining locus names are based on nearest gene, and do not refer to any inferred biological function. We note that this Table excludes 7 loci filtered as potential false positive associations due to differential tagging of a causal stress test SNP (Table S10), and 1 locus excluded based on specifying a range of disorder prevalence values (as opposed to specifying only the most likely disorder prevalence). SCZ, schizophrenia; BIP, bipolar disorder; MDD, major depressive disorder; ADHD, attention deficit/hyperactivity disorder; ANO, anorexia nervosa; ASD, autism spectrum disorder; OCD, obsessive–compulsive disorder; TS, Tourette's Syndrome and Other Tic Disorders.

**Table S14. Summary of CC-GWAS results for eight psychiatric disorders for the main analyses and at different Exact p-value thresholds.** This Table is an extension of Table 1. For each pair of eight psychiatric disorders, we report the case-control sample sizes, #SNPs, prevalence ($K$)[4], liability-scale heritability estimated using stratified LD score regression[5–7] ($h^2$), genetic correlation estimated using cross-trait LD score regression[8] ($r_g$), CC-GWAS$_{OLS}$ weights, number of independent genome-wide significant loci for each case-control comparison, number of independent genome-significant CC-GWAS loci, number of independent genome-significant CC-GWAS loci that are CC-GWAS-specific, number of independent genome-significant when applying the CC-GWAS$_{OLS}$ component only, number of independent genome-significant CC-GWAS loci when applying different thresholds of significance for the CC-GWAS$_{Exact}$ component (note that CC-GWAS is based on the threshold p<$10^{-4}$). CC-GWAS$_{Exact}$ weights are equal to $(1 - K_A)$ for disorder A and – $(1 - K_B)$ for disorder B. We note that this Table reports the number of loci before filtering potential false positive associations due differential tagging of a causal stress test SNP, and without specifying a range of disorder prevalence values (these filters had minimal effect in our main analyses). SCZ, schizophrenia; BIP, bipolar disorder; MDD, major depressive disorder; ADHD, attention deficit/hyperactivity disorder; ANO, anorexia nervosa; ASD, autism spectrum disorder; OCD, obsessive–compulsive disorder; TS, Tourette's Syndrome and Other Tic Disorders.

**Table S15. Summary of CC-GWAS results for eight psychiatric disorders with alternative clumping strategy.** The results in Table 1 and Table S14 were clumped by clumping correlated SNPs ($r^2$≥0.1) in 3MB windows and collapsing remaining SNPs within 250kb in line with ref.[12]; the results in this Table are clumped by clumping correlated SNPs ($r^2$≥0.01) in 5MB windows and collapsing remaining SNPs within 100kb in line with ref.[13]. For each pair of eight psychiatric disorders, we report the case-control sample sizes, #SNPs, prevalence $(K)$[4], liability-scale heritability estimated using stratified LD score regression[5–7] $(h^2)$, genetic correlation estimated using cross-trait LD score regression[8] $(r_g)$, CC-GWAS$_{OLS}$ weights, number of independent genome-wide significant loci for each case-control comparison, number of independent genome-significant CC-GWAS loci, and number of independent genome-significant CC-GWAS loci that are CC-GWAS-specific. CC-GWAS$_{Exact}$ weights are equal to $(1-K_A)$ for disorder A and $-(1-K_B)$ for disorder B. Note that the heritability estimates, genetic correlation estimates and CC-GWAS$_{OLS}$ weights are identical to those reported in Table 1 and Table S14. We note that this Table reports the number of loci before filtering potential false positive associations due to differential tagging of a causal stress test SNP, and without specifying a range of disorder prevalence values (these filters had minimal effect in our main analyses). SCZ, schizophrenia; BIP, bipolar disorder; MDD, major depressive disorder; ADHD, attention deficit/hyperactivity disorder; ANO, anorexia nervosa; ASD, autism spectrum disorder; OCD, obsessive–compulsive disorder; TS, Tourette's Syndrome and Other Tic Disorders.

**Table S16. Summary of CC-GWAS results for eight psychiatric disorders when correcting for the LD score regression intercept.** The results are analogue to Table 1 and Table S14, but with input GWAS results corrected for their LD score regression intercept in line with Turley et al.[1] (see Methods). For each pair of eight psychiatric disorders, we report the case-control sample sizes, #SNPs, prevalence $(K)$[4], liability-scale heritability estimated using stratified LD score regression[5–7] $(h^2)$, genetic correlation estimated using cross-trait LD score regression[8] $(r_g)$, CC-GWAS$_{OLS}$ weights, number of independent genome-wide significant loci for each case-control comparison, number of independent genome-significant CC-GWAS loci, and number of independent genome-significant CC-GWAS loci that are CC-GWAS-specific. CC-GWAS$_{Exact}$ weights are equal to $(1-K_A)$ for disorder A and $-(1-K_B)$ for disorder B. Note that compared to the results reported in Table 1 and Table S14, the heritability estimates are smaller, that CC-GWAS$_{OLS}$ weights are smaller (reflecting lower expected signal to noise ratio based on the lower heritability estimates), and the numbers of independent loci detected are proportionally smaller for the case-control comparisons as for the case-case comparisons. However, we believe this correction is overly conservative, as S-LDSC intercept attenuation ratios[13] were relatively small (Table S17), implying little evidence of confounding. We note that this Table reports the number of loci before filtering potential false positive associations due differential tagging of a causal stress test SNP, and without specifying a range of disorder prevalence values (these filters had minimal effect in our main analyses). SCZ, schizophrenia; BIP, bipolar disorder; MDD, major depressive disorder; ADHD, attention deficit/hyperactivity disorder; ANO, anorexia nervosa; ASD, autism spectrum disorder; OCD, obsessive–compulsive disorder; TS, Tourette's Syndrome and Other Tic Disorders.

**Table S17. Stratified LD score regression results of the eight psychiatric disorders (including the attenuation ratios).** For each of the eight psychiatric disorders, we report the number of cases, number of controls, the prevalence $(K)$[4], liability-scale heritability $(h2\_liab)$ and observed-scale heritability $(h2\_liab)$ estimated using stratified LD score regression[5–7], the intercept of stratified LD score regression, the *mean X2* and the attenuation ratio[13]. SCZ, schizophrenia; BIP, bipolar disorder; MDD, major depressive disorder; ADHD, attention deficit/hyperactivity disorder; ANO, anorexia nervosa; ASD, autism spectrum disorder; OCD, obsessive–compulsive disorder; TS, Tourette's Syndrome and Other Tic Disorders.

**Table S18. Expected inflation of CC-GWAS results based on inflated input GWAS results.** This Table reports analytical computations and simulation results at null-null SNPs when the input case-control GWAS results would be biased (with decreased standard error and increased variance of $z$-values, i.e. $> 1$). We report the genetic correlation between disorder A and disorder B used for simulation, which input case-control GWAS results were set to be inflated (by multiplying the standard error of the respective GWAS result with 0.9), the variance of $z$-values of biased case-control GWAS results, and the variance of $z$-values of CC-GWAS and CC-GWAS+ results. Note that the z-values of CC-GWAS have comparable (or smaller) variance as the biased input case-control GWAS results. Parameter are in line with Table S2: $h^2$=0.2, prevalence $K$=0.01, and sample size 4,000 cases + 4,000 controls for disorder A; liability-scale $h^2$=0.1, prevalence $K$=0.15, and sample size 4,000 cases + 4,000 controls for disorder B; $m$=1,000 causal SNPs for each disorder; and genetic correlation $r_g$=0.5 between disorders. We report the mean (standard error) of 50 simulation runs (see Methods).

**Table S19. Summary of CC-GWAS results for eight psychiatric disorders when applying non-stratified LD score regression.** The results are analogue to Table 1 and Table S14, but with heritability estimates based on non-stratified LD score regression[14]. For each pair of eight psychiatric disorders, we report the case-control sample sizes, #SNPs, prevalence $(K)$[4], liability-scale heritability estimated using stratified LD score regression[5–7] $(h^2)$, genetic correlation estimated using cross-trait LD score regression[8] $(r_g)$, CC-GWAS$_{OLS}$ weights, number of independent genome-wide significant loci for each case-control comparison, number of independent genome-significant CC-GWAS loci, and number of independent genome-significant CC-GWAS loci that are CC-GWAS-specific. CC-GWAS$_{Exact}$ weights are equal to $(1-K_A)$ for disorder A and $-(1-K_B)$ for disorder B. Note that compared to the results reported in Table 1 and Table S14, the heritability estimates are smaller (as expected[7]), and that CC-GWAS$_{OLS}$ weights are smaller (reflecting lower expected signal to noise ratio based on the lower heritability estimates), while the numbers of independent loci detected are comparable. We note that this Table reports the number of loci before filtering potential false positive associations due differential tagging of a causal stress test SNP, and without specifying a range of disorder prevalence values (these filters had minimal effect in our main analyses). SCZ, schizophrenia; BIP, bipolar disorder; MDD, major depressive disorder; ADHD, attention deficit/hyperactivity disorder; ANO, anorexia nervosa; ASD, autism spectrum disorder; OCD, obsessive–compulsive disorder; TS, Tourette's Syndrome and Other Tic Disorders.

**Table S20. Summary of CC-GWAS results for eight psychiatric disorders under different assumptions about the number of causal SNPs.** The results are analogue to Table 1 and Table S14, but assuming half $(m = 5{,}000)$ and double $(m = 20{,}000)$ the number independent causal SNPs respectively. For each pair of eight psychiatric disorders, we report the case-control sample sizes, #SNPs, prevalence $(K)$[4], liability-scale heritability estimated using stratified LD score regression[5–7] $(h^2)$, genetic correlation estimated using cross-trait LD score regression[8] $(r_g)$, number of assumed causal SNPs, CC-GWAS$_{OLS}$ weights, number of independent genome-wide significant loci for each case-control comparison, number of independent genome-significant CC-GWAS loci, and number of independent genome-significant CC-GWAS loci that are CC-GWAS-specific. CC-GWAS$_{\text{Exact}}$ weights are equal to $(1 - K_A)$ for disorder A and – $(1 - K_B)$ for disorder B. Note that compared to the results reported in Table 1 and Table S14, the CC-GWAS$_{\text{OLS}}$ weights are larger for $m = 5{,}000$ (reflecting larger expected signal to noise ratio) and smaller for $m = 20{,}000$ (reflecting smaller expected signal to noise ratio). We note that this Table reports the number of loci before filtering potential false positive associations due differential tagging of a causal stress test SNP, and without specifying a range of disorder prevalence values (these filters had minimal effect in our main analyses). SCZ, schizophrenia; BIP, bipolar disorder; MDD, major depressive disorder; ADHD, attention deficit/hyperactivity disorder; ANO, anorexia nervosa; ASD, autism spectrum disorder; OCD, obsessive–compulsive disorder; TS, Tourette's Syndrome and Other Tic Disorders.

**Table S21. CC-GWAS results of SCZ, BIP and MDD when varying the specified population prevalence for the CC-GWAS$_{\text{OLS}}$ weights.** We performed CC-GWAS analyses varying the specified disorder prevalences for the CC-GWAS$_{\text{OLS}}$ weights for SCZ (0.4%-1.0%)[4,12], BIP (0.5%-2.0%)[15], and MDD (16%-30%)[4,16]. We report the specified prevalences in the main analyses (see Table 1), the adjusted prevalences in the secondary analyses, the CC-GWAS$_{\text{OLS}}$ weights in the main analyses (see Table 1), the CC-GWAS$_{\text{OLS}}$ weights in the secondary analyses, the number of significant loci in the main analyses (see Table 1), the number of loci in the secondary analyses, the number of overlapping loci between the main analyses and secondary analyses based on the lead SNPs, and the number of overlapping loci between the main analyses and secondary analyses based on overlap of any pair of significant SNP in the loci. CC-GWAS$_{\text{Exact}}$ weights are equal to $(1 - K_A)$ for disorder A and – $(1 - K_B)$ for disorder B. Overall, results changed little when varying the disorder prevalences for the CC-GWAS$_{\text{OLS}}$ weights.

**Table S22. List of 9 CC-GWAS+ loci for SCZ vs. BIP.** We extended CC-GWAS to CC-GWAS+ by including results of a direct case-case comparison based on 23,585 SCZ cases and 15,270 BIP cases[17] to the case-control comparison of SCZ[12] (40,675 cases and 64,643 controls) and BIP[15] (20,352 cases and 31,358). We analyzed 4,537,755 SNPs, and assumed 10,000 independent causal SNPs[18]. The CC-GWAS+$_{\text{OLS}}$ weights were 0.35 for SCZ case-control, -0.25 for BIP case-control and 0.22 SCZ case-BIP case, and the CC-GWAS+$_{\text{Exact}}$ plus weights were 0, 0 and 1 respectively (see Methods). For each CC-GWAS+ locus, we report the lead CC-GWAS+ SNP and its chromosome, physical position, and reference allele frequency, the nearest gene, the respective case-control effect sizes and *p*-values, the direct case-case comparison effect size and *p*-value, the CC-GWAS+$_{\text{OLS}}$ case-case effect size and *p*-value, the CC-GWAS+$_{\text{Exact}}$ case-case effect size and *p*-value, and an indicator of CC-GWAS-specific loci. SCZ, schizophrenia; BIP, bipolar disorder.

**Table S23. SMR results of CC-GWAS-specific loci for eight psychiatric disorders.** For each significant SMR colocalization result, we report the disorder pair and CC-GWAS-specific locus, the number of SMR probes tested for that disorder pair, the SMR significance p-value threshold for that pair, the gene, the tissue, and the outcome of the SMR software (the probe ID and its chromosome and base pair position; the top SNP with the strongest eQTL signal for that probe and its chromosome, base pair position, allele A1 (=effect allele), allele A2 and A1-allele frequency; the beta, standard error and p-value of CC-GWAS result at the top eQTL SNP; the beta, standard error and p-value of the eQTL result at the top eQTL SNP; the beta, standard error and p-value of SMR analyses; the p-value of the HEIDI test, and the number of SNPs tested in the HEIDI test). SCZ, schizophrenia; BIP, bipolar disorder; MDD, major depressive disorder; ADHD, attention deficit/hyperactivity disorder; ANO, anorexia nervosa;

ASD, autism spectrum disorder; OCD, obsessive–compulsive disorder; TS, Tourette's Syndrome and Other Tic Disorders.

**Table S24. Case-control and case-case effect sizes at the *KLF16* and *KLF6* CC-GWAS loci**. For the CC-GWAS lead SNPs at the SCZ vs. BIP *KLF16* locus and SCZ vs. MDD *KLF* 6 locus, we show the chromosome, base pair position, beta and p-values of the SCZ, BIP and MDD case-control effect sizes, and beta and p-values of the CC-GWAS$_{OLS}$ case-case effect sizes of SCZ vs. BIP, SCZ vs. MDD and BIP vs. MDD. SCZ, schizophrenia; BIP, bipolar disorder; MDD, major depressive disorder.

**Table S25. Summary of CC-GWAS results in discovery data sets of replication analyses for SCZ vs. MDD, the three autoimmune disorders and RA vs BC.** The first part of this Table (Discovery data) mimics Table 1. For each pair of SCZ vs. MDD, CD vs. UC, CD vs. RA, UC vs. RA and BC vs. RA for the discovery data, we report the case-control sample sizes, #SNPs, prevalence ($K$)[4], liability-scale heritability estimated using stratified LD score regression[5–7] ($h^2$), genetic correlation estimated using cross-trait LD score regression[8] ($r_g$), CC-GWAS$_{OLS}$ weights, number of independent genome-wide significant loci for each case-control comparison, number of independent genome-significant CC-GWAS loci, and number of independent genome-significant CC-GWAS loci that are CC-GWAS-specific. For the replication data, we report the case-control sample sizes, and the number of loci from the discovery data with matching replication data in the case-control analyses and case-case analyses. SCZ, schizophrenia; MDD, major depressive disorder; CD, Crohn's disorder; UC, ulcerative colitis; RA, rheumatoid arthritis; BC, breast cancer.

**Table S26. List of 119 CC-GWAS loci analyzed in replication analyses.** This Table reports the numerical results of Figure 5. For each CC-GWAS locus used in replication analyses, we report the respective disorders, the lead SNP in the discovery analyses, the SNP used for replication with its chromosome and base-pair position (selected as the most significant discovery SNP in the locus with matching replication data), the discovery set effect sizes and p-values for the case-control comparison and the CC-GWAS$_{OLS}$ and CC-GWAS$_{Exact}$ case-case comparisons, and the replication effect sizes and p-values for the case-control comparison and the CC-GWAS$_{OLS}$ and CC-GWAS$_{Exact}$ case-case comparisons. An overview of these results is reported in Table S25. SCZ, schizophrenia; MDD, major depressive disorder; CD, Crohn's disorder; UC, ulcerative colitis; RA, rheumatoid arthritis.

**Table S27. List of 186 case-control loci analyzed in within-disorder replication analyses.** This Table reports numerical values of Figure S15. For each case-control locus used for within disorder replication analyses, we report the respective disorder, the SNP used for replication with its chromosome and base-pair position (selected as the most significant discovery SNP in the locus with matching replication data), the discovery set case-control effect size and p-value, and the replication case-control effect size and p-value. An overview of these results is reported in Table S25. SCZ, schizophrenia; MDD, major depressive disorder; CD, Crohn's disorder; UC, ulcerative colitis; RA, rheumatoid arthritis.

**Table S28. CC-GWAS SCZ vs. BIP results at the 2 loci identified by Ruderfer et al.** For the two SCZ vs. BIP loci identified by Ruderfer et al.[17], this Table reports results of the 2 Ruderfer et al. lead SNPs (these SNPs were not included CC-GWAS in CC-GWAS analyses, because no SCZ case-control[12] results were available for these SNPs), and the Ruderfer et al. and CC-GWAS results at the SNP within 100kb with the smallest Ruderfer et al. p-value that was also analyzed with CC-GWAS. Note that these results present no independent replication, because the data of Ruderfer et al. were included in the CC-GWAS analyses. Results were somewhat different (as would be expected due to different sample sets and different methods), but generally consistent. (Note that a CC-GWAS effect-size $< 0$ corresponds to and $OR < 1$.)

**Table S29. Ruderfer et al. results at 12 CC-GWAS SCZ vs. BIP loci.** For the lead SNPs of the 12 CC-GWAS SCZ vs. BIP loci, this Table reports the Ruderfer et al.[17] results of a direct SCZ vs. BIP case-case comparison. Note that these results present no independent replication, because the data of Ruderfer et al. were included in the CC-GWAS analyses. Results were somewhat different (as would be expected due to different sample sets and different methods), but generally consistent. (Note that a CC-GWAS effect-size $< 0$ corresponds to and $OR < 1$, and a CC-GWAS effect-size $> 0$ corresponds to and $OR > 1$.)

**Table S30. Expectation of CC-GWAS type I error in the presence of population stratification impacting allele frequency.** We report the type I error at loci with varying differences in population allele frequencies across the case-control data of A1A0 and the case-control data of B1B0. The A1A0 case-control results are fixed in pop0 (with affective allele frequency $EAF = 0.5$) and compared to B1B0 case-control results from pop0 ($EAF = 0.5$, i.e. from the same population), pop1 ($EAF = 0.45$, reflecting an allele frequency difference with pop0 typical of differences within a continental population[19]), pop2 ($EAF = 0.4$), pop3 ($EAF = 0.3$, reflecting an allele frequency difference with pop0 typical for differences between continental populations[20]), and pop4 ($EAF = 0.1$). We report the disorder, its population prevalence (*K*); the effective allele frequency in the population, in cases and in controls; the case-control effect sizes expressed as per allele OR, variance explained on the observed scale with 50/50 case ascertainment, and as standardized beta on the observed scale with 50/50 case ascertainment; and the type I error of CC-GWAS and a direct case-case comparison (GWAS A1B1). Panel A shows the expectation for null-null SNPs, and Panel B shows the expectation for stress test SNPs. The parameters are in line with Figure 2: $h^2$=0.2, prevalence *K*=0.01, and sample size 100,000 cases + 100,000 controls for disorder A; liability-scale $h^2$=0.1, prevalence *K*=0.15, and sample size 100,000 cases + 100,000 controls for disorder B; *m*=5,000 causal SNPs for each disorder; and genetic correlation $r_g$=0.5 between disorders. Stress test SNPs have the same allele frequency in A1 vs. B1 (when in the same population) and explain 0.10% of liability variance in A1 vs. A0 and 0.29% of liability variance in B1 vs. B0, as in Figure 2.

**Table S31. False discovery rate correction (FDR) of CC-GWAS results.** We report (in rows) the total number of SNPs tested, the corresponding FDR p-value threshold for an FRD of 0.05, and the number of significant SNPs for this threshold, for (in columns) the average per disorder pair over the 28 pairs of eight psychiatric disorders (i.e. average of per pair FDR), the total of the three comparisons of SCZ, BIP and MDD (i.e. global FDR for these three disorders), and the total of the 28 pairs of eight psychiatric disorders (i.e. global FDR for the eight disorders). The FDR was computed with the Benjamini-Hochberg procedure[21]. Note that global FDR is more stringent than the per pair FDR (except for the global FDR of SCZ, BIP and MDD which reflect more powerful analyses), and note that all FDR p-value thresholds are >> $5x10^{-8}$, indicating that our presented results are conservatively corrected with respect to an FDR of 0.05. SCZ, schizophrenia; BIP, bipolar disorder; MDD, major depressive disorder.

**Table S32. Distribution of CC-GWAS p-values compared to case-control p-values for SCZ, BIP and MDD.** We report the proportion of loci with relatively large genome-wide significant p-values ($1.7x10^{-8}$ < P< $5x10^{-8}$) in case-control analyses and in CC-GWAS analyses of SCZ, BIP and MDD. (The threshold of $1.7x10^{-8}$ is chosen as $5x10^{-8}/3$.) We note that these proportions are similar for all CC-GWAS loci and case-control analyses, as this is a general property of polygenic architectures. The p-values of the CC-GWAS-specific loci are larger than for all CC-GWAS loci, because CC-GWAS loci with very small p-values are likely to also have significant case-control association (thus not being CC-GWAS-specific).

**Table S33. Overview of covariates applied in case-control analyses of eight psychiatric disorders.** None of the eight case-control GWAS was corrected for a genetically correlated covariate, suggesting the covariates applied will not increase type I error at stress test SNP. Note that studies corrected for different numbers of principal components, but this has no impact on results (assuming the studies adequately corrected for population stratification).

**Table S34. Simulation results of CC-GWAS applied on subtype data.** CC-GWAS was designed to compare two disorders (with different definitions of controls and potential overlap of cases), but it is also of interest to compare subtypes within a disorder (with same definitions of controls and no overlap of cases). We compared analytical expectation of CC-GWAS results (falsely assuming results are based on two disorder) to simulation results of CC-GWAS applied on subtype data. We report (A) the power to detect SNPs with effect sizes following a bivariate normal distribution and (B) the type I error rate for loci with no effect on A1A0 or B1B0 ("null-null" SNPs) for each of five methods: CC-GWAS, the CC-$GWAS_{OLS}$ component, the CC-$GWAS_{Exact}$ component, the Delta method, and CC-GWAS+ (see Methods). Subtype data were simulated by first simulating two disorders in line with the main simulations (see Methods and Table S2), and subsequently defining subtype-controls as double-controls for both disorders, subtype-A as A cases, and subtype-B as B cases that are controls for A. Default parameter settings are in line with the main simulations in Table S2: $h^2$=0.2, prevalence $K$=0.01, and sample size 4,000 cases + 4,000 controls for disorder A; liability-scale $h^2$=0.1, prevalence $K$=0.15, and sample size 4,000 cases + 4,000 controls for disorder B; $m$=1,000 causal SNPs for each disorder; and genetic correlation $r_g$=0.5 between disorders. Levels of significance were set as $p < 0.01$ for the CC-$GWAS_{OLS}$ weights and $p < 0.05$ for the CC-$GWAS_{Exact}$ weights. We report the mean (standard error) of 50 simulation runs. Note that the analytical expectations and simulation results correspond reasonably, suggesting that CC-GWAS can also be applied on data of subtypes. Further note that stress test SNPs in this scenario are SNPs with the exact same effect on both subtypes (contrary to when comparing two different disorders, the controls are exactly the same), and that the CC-$GWAS_{Exact}$ component should thus be replaced with the delta method (i.e. with weights 1 and -1 for the subtypeA-control and subtypeB-control comparisons).

**Table S35. Comparison of different approaches to approximate standardized observed-scale case-control and case-case effect sizes.** We compare several approaches to obtain effect sizes based on simulated individual level data. Simulation was performed based on the liability scale as described in the Method section. For each parameter setting, we report the liability variance explained by the SNP; the prevalence of A ($K$); the effective allele frequency (MAF); the per allele odds ratio (OR); the standardized observed case-control effect size based 50/50 case ascertainment obtained from (i) theory as the square root of the variance explained on the observed scale, based on the standard transformation of variance explained on the liability scale[22,23], (ii) linear regression (golden standard),

(iii) via the allele frequency in cases ($p_{i,A1}$) and controls ($p_{i,A0}$) as $\frac{p_{i,A1}-p_{i,A0}}{\sqrt{2p_i(1-p_i)}}$, (iv) via $N_{eff}$ as $z_{i,logistic\ regression}\sqrt{\frac{1}{N_{Eff}}}$, and (v) via the OR based on equation 5 in the paper from Lloyd-Jones et al.[24] (see Methods); the correlation of these betas with the betas obtained from linear regression; and the standardized observed case-case effect size based 50/50 ascertainment obtained via theory, via linear regression (golden standard), and via the CC-GWAS$_{\text{Exact}}$ component as $(1-K_A)\hat{\beta}_{i,A1A0} - (1-K_B)\hat{\beta}_{i,B1B0}$. For all parameter settings, SNPs are simulated to explain 0.1% of variance in disorder B with a population prevalence of $K_B = 0.05$, and sample size were set at 4,000 cases + 8,000 controls for disorder A and 3,000 cases + 2,000 controls for disorder A (these somewhat random sample sizes were chosen to confirm validity of approximations of beta also for non-balanced sample sizes). Simulations were repeated 300 times for each parameter setting, and the mean (standard error) respectively correlation across these 300 iterations are reported. Note that the A1A0 beta via $N_{eff}$ and beta via OR both provide appropriate approximation of beta via linear regression, but that the beta via $N_{eff}$ has a slightly large correlation with beta via logistic regression. Therefore, CC-GWAS applies the transformation via Neff as primary transformation and the transformation via OR to provide a rough double-check of whether $N_{Eff}$ has been defined accurately.

**Table S36. $F_{ST,causal}$: analytical computations vs. simulation.** We report analytical computations and simulation results of $F_{ST,causal}$ for the comparisons A1A0, B1B0, A1B1, A1B0, A0B1 and A0B0. Simulation parameter settings are: $h^2$=0.2, prevalence $K$=0.01, and sample size 4,000 cases + 4,000 controls for disorder A; liability-scale $h^2$=0.1, prevalence $K$=0.15, and sample size 4,000 cases + 4,000 controls for disorder B; $m$=1,000 causal SNPs for each disorder; and genetic correlation $r_g$=0.5 between disorders. Panel A reports simulations with 1,000 SNPs with effects following a bivariate normal distribution (see Methods). Panel B reports simulation based on 1,000 SNPs with subsets of SNPs: (i) with the exact same effects on both disorders from a uniform distribution, (ii) with effects on disorder A only from the same uniform distribution, (iii) with effects on disorder B only from the same uniform distribution, and (iv) no effect on either disorder (note that the number of SNPs in (i)-(iv) define the $r_g$ as indicated). The results from Panel B indicate that the equations of $F_{ST,causal}$ require less stringent assumptions than CC-GWAS.

**Table S37. $F_{ST}$ between GWAS datasets of psychiatric disorders.** This Table reports the $F_{ST}$ for the three pairs of psychiatric disorders with in-sample allele frequencies available. (Note we refer here to $F_{ST}$ as applied in population genetics[20]; not to be confused with $F_{ST,causal}$). We only report comparisons that were used in any of our (primary or secondary) analyses. All $F_{ST}$ values were smaller than 0.0006, suggesting that our simulations of tagging differences based on 25k British UK Biobank samples and 25k “other European” UK Biobank samples adequately represented our empirical analyses.

**Table S38. Tagging differences between 25k British UK Biobank samples and 25k “other European” UK Biobank samples.** For 68,986,554 SNP-pairs from 1000 Genomes on chromosome 1 with MAF>0.01 within 100kb of each other, we report the absolute values of the LD (signed correlation) differences between 25k British UK Biobank samples and 25k “other European” UK Biobank samples. The 68,986,554 SNP-pairs were selected based on taking all SNP-pairs in 2,485 consecutive 100kb windows (the range that LD typically spans[25,26]) on chromosome 1. We conclude that most SNP-pairs show high concordance of LD between both populations, but that some SNP-pairs also show moderate difference in LD (despite the fact that both populations are from the same continent). This illustrates the necessity to include in CC-GWAS the filtering step to exclude false positive associations due to differential tagging of a causal stress test SNP.

**Table S39. Intercept of cross-trait LD score regression vs. analytical expectation based on sample-overlap of controls.** For each pair of eight psychiatric disorders, we report the case-control sample sizes, number of overlapping controls, intercept from cross-trait LD score regression[8], and analytically expected intercept based on sample overlap (see Methods). The cross-trait LD score regression intercept is used to model the covariance of error terms. Overestimation of the covariance of error terms can lead to inflated type I error of CC-GWAS results. Therefore, CC-GWAS is based on the minimum of the intercept from cross-trait LD score regression and the analytical expectation. The information on sample-overlap was kindly provided in personal communication with Dr. S. Ripke and Dr. V. Trubetskoy. SCZ, schizophrenia; BIP, bipolar disorder; MDD, major depressive disorder; ADHD, attention deficit/hyperactivity disorder; ANO, anorexia nervosa; ASD, autism spectrum disorder; OCD, obsessive–compulsive disorder; TS, Tourette's Syndrome and Other Tic Disorders.

### Supplementary Figures

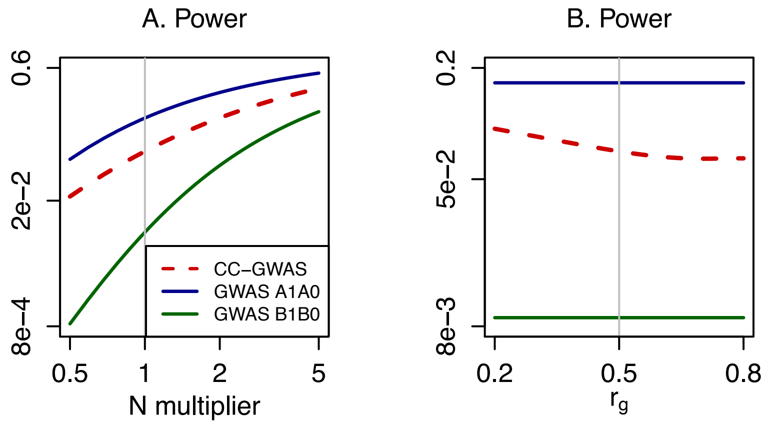

**Figure S1. Power of CC-GWAS and input case-control GWAS.** Analogue to main Figure 2 and based on the same default parameter values. We report the power to detect SNPs with effect sizes following a bivariate normal distribution, for each of three methods: CC-GWAS, input case-control GWAS A1A0, and input case-control GWAS B1B0. Note that the power of CC-GWAS lays in between the power of GWAS A1A0 and GWAS B1B0.

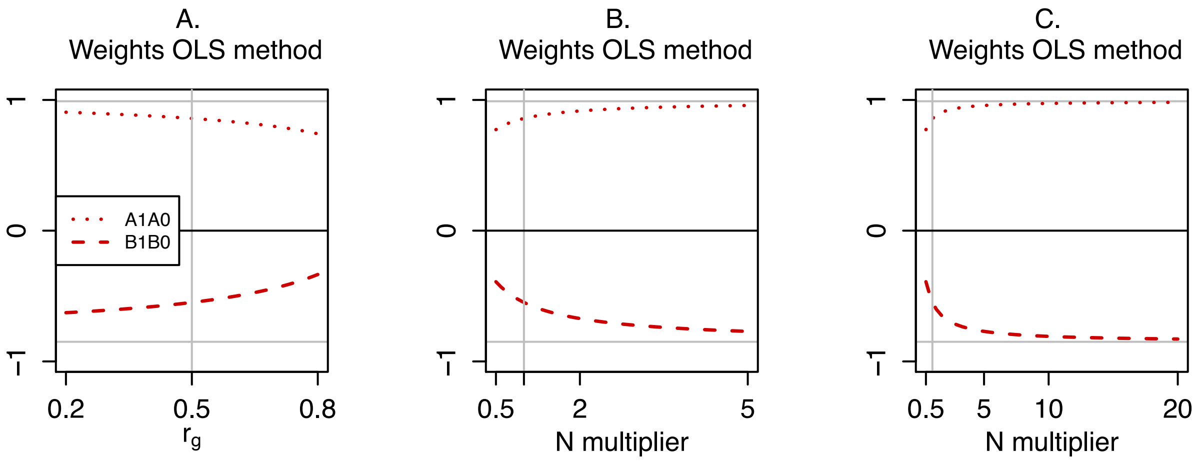

**Figure S2. CC-GWAS$_{OLS}$ weights of CC-GWAS based on analytical computations.** Analogue to main Figure 2 and based on the same default parameter values. We report the CC-GWAS$_{OLS}$ weights for the A1A0 input GWAS results (dotted line) and for the B1B0 input GWAS results (dashed line). The corresponding CC-GWAS$_{Exact}$ weights are displayed as horizontal grey lines ($1 - K_A = 0.99$ for A1A0 and $-(1 - K_B) = -0.85$ for B1B0). Panel A corresponds to the second row of Figure 2, and Panel B corresponds to the first row of Figure 2. Panel C illustrates that CC-GWAS$_{OLS}$ weights converge to the CC-GWAS$_{Exact}$ weight for even further increasing sample size.

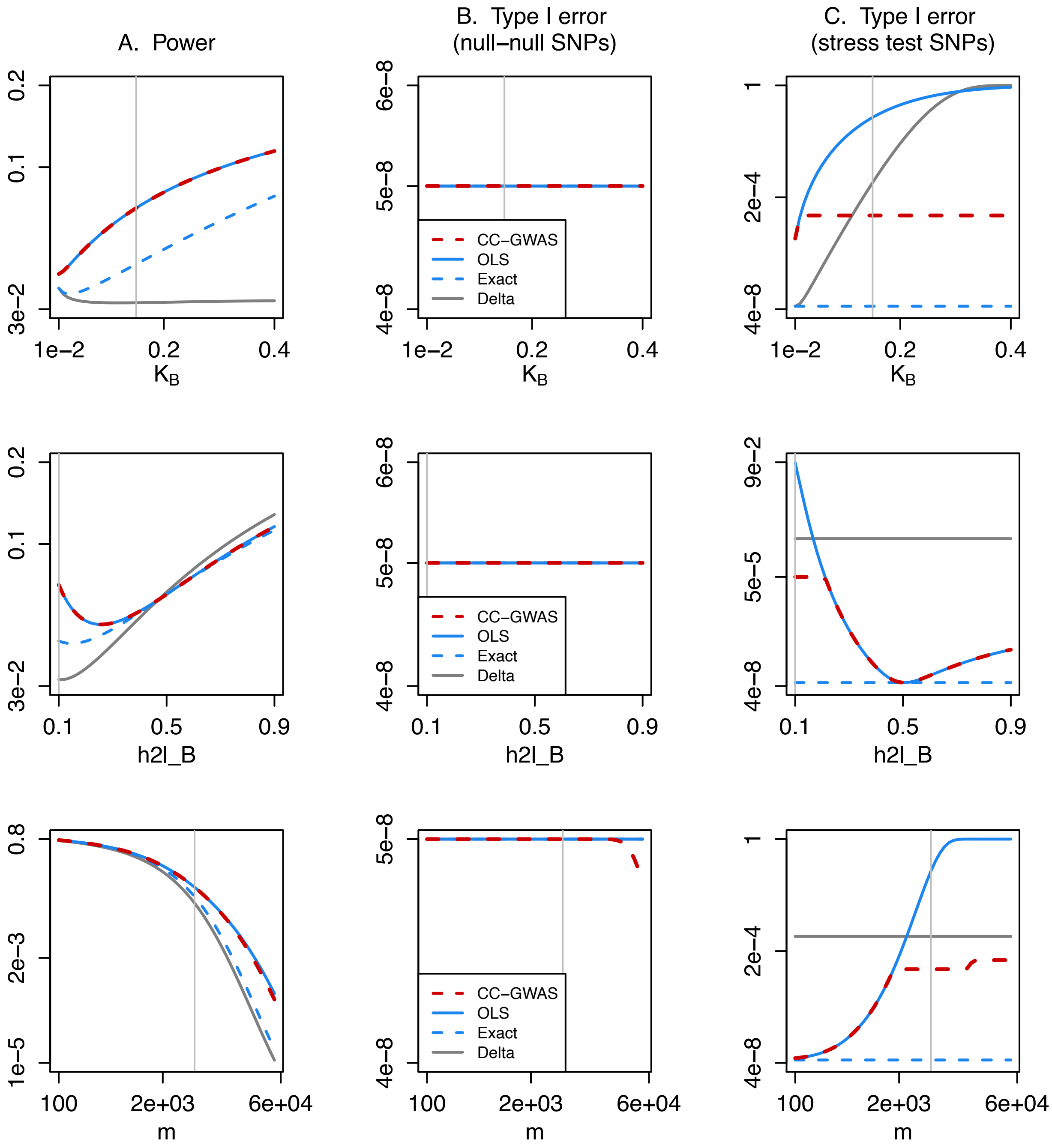

**Figure S3. Power and type I error of CC-GWAS for varying prevalence, heritability and number of causal loci.** This Figure is analogue to main Figure 2 and based on the same default parameter values, with varying values of the prevalence of disorder B ($K_B$), heritability of B ($h^2_{l,B}$) and number of causal loci ($m$). We report (A) the power to detect SNPs with effect sizes following a bivariate normal distribution, (B) the type I error rate for loci with no effect on A1A0 or B1B0 ("null-null" SNPs) and (C) the type I error rate for SNPs with the same allele frequency in A1 vs. B1 that explain 0.10% of variance in A1 vs. A0 and 0.29% of variance in B1 vs. B0 ("stress test" SNPs), for each of four methods: CC-GWAS, the CC-GWAS$_{OLS}$ component, the CC-GWAS$_{Exact}$ component, and a naïve Delta method (see main text).

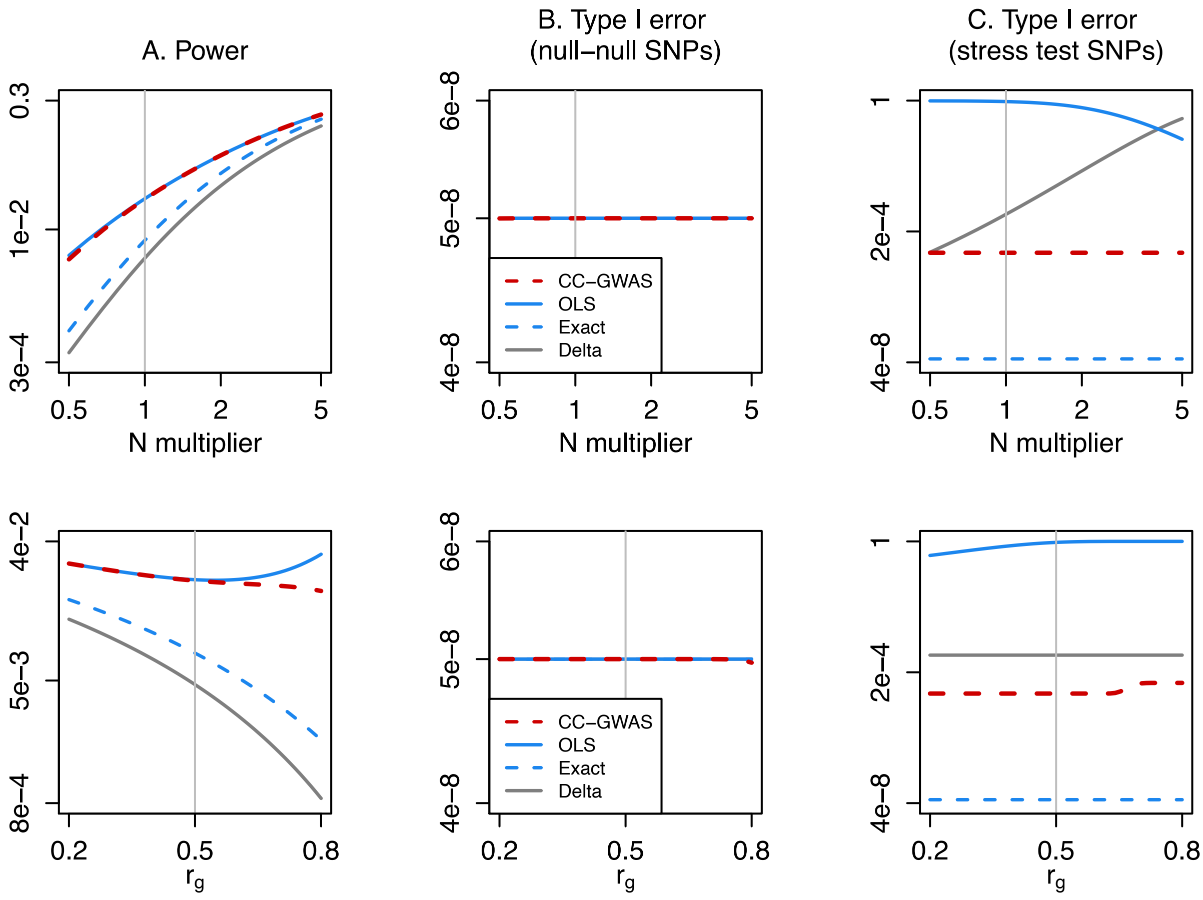

**Figure S4. Power and type I error of CC-GWAS for 10,000 causal loci.** This Figure is analogue to main Figure 2 and based on the same parameter values, except for the number of causal loci $m = 10{,}000$. We report (A) the power to detect SNPs with effect sizes following a bivariate normal distribution, (B) the type I error rate for loci with no effect on A1A0 or B1B0 ("null-null" SNPs) and (C) the type I error rate for SNPs with the same allele frequency in A1 vs. B1 that explain 0.10% of variance in A1 vs. A0 and 0.29% of variance in B1 vs. B0 ("stress test" SNPs), for each of four methods: CC-GWAS, the CC-GWAS$_{OLS}$ component, the CC-GWAS$_{Exact}$ component, and a naïve Delta method (see main text).

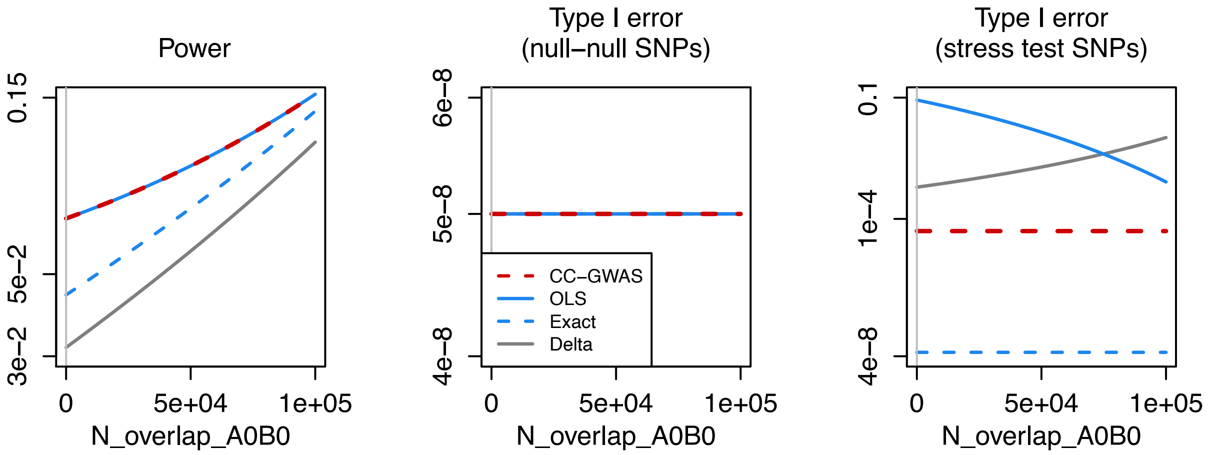

**Figure S5. Power and type I error of CC-GWAS with overlap of controls.** This Figure is analogue to main Figure 2 and based on the same parameter values, with increasing number of overlapping controls (note that overlap of 1e+5 corresponds to full overlap of controls; Figure 2 is based on no overlap of controls). We report (A) the power to detect SNPs with effect sizes following a bivariate normal distribution, (B) the type I error rate for loci with no effect on A1A0 or B1B0 (“null-null” SNPs) and (C) the type I error rate for SNPs with the same allele frequency in A1 vs. B1 that explain 0.10% of variance in A1 vs. A0 and 0.29% of variance in B1 vs. B0 (“stress test” SNPs), for each of four methods: CC-GWAS, the CC-GWAS$_{OLS}$ component, the CC-GWAS$_{Exact}$ component, and a naïve Delta method (see main text).

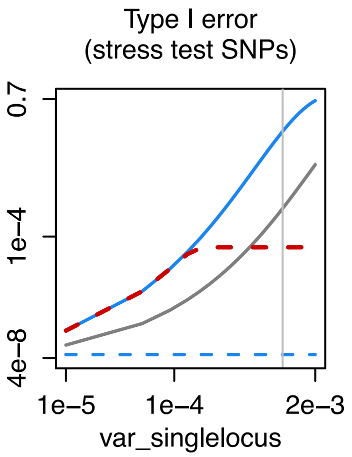

**Figure S6. Type I error at stress test SNPs.** This Figure is analogue to main Figure 2C and based on the same parameter values, with varying proportions of variance explained by the stress test SNPs in the liability of A1A0 (and corresponding varying variance explained on B1B0 such that the allele frequency in A1 equals the allele frequency in B1). We report the type I error rate for these stress test for each of four methods: CC-GWAS, the CC-GWAS$_{OLS}$ component, the CC-GWAS$_{Exact}$ component, and a naïve Delta method (see main text).

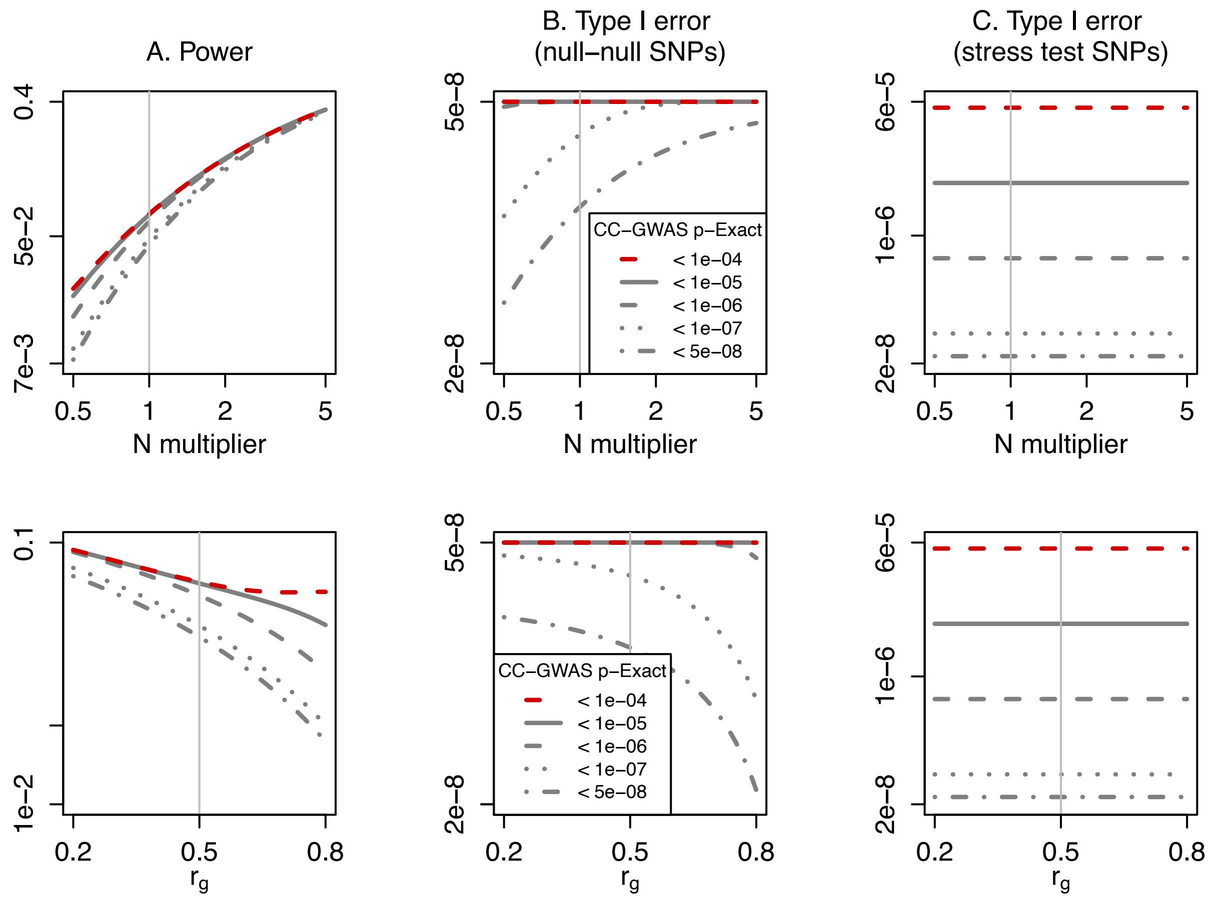

**Figure S7. Power and type I error of CC-GWAS at different Exact p-value thresholds.** This Figure is analogue to main Figure 2 and based on the same parameter values. We report (A) the power to detect SNPs with effect sizes following a bivariate normal distribution, (B) the type I error rate for loci with no effect on A1A0 or B1B0 ("null-null" SNPs) and (C) the type I error rate for SNPs with the same allele frequency in A1 vs. B1 that explain 0.10% of variance in A1 vs. A0 and 0.29% of variance in B1 vs. B0 ("stress test" SNPs), for each of five methods: CC-GWAS with Exact threshold $p < 10^{-4}$ (this is CC-GWAS as presented in Figure 2 and applied throughout this study), CC-GWAS with Exact threshold $p < 10^{-4}$, CC-GWAS with Exact threshold $p < 10^{-5}$, CC-GWAS with Exact threshold $p < 10^{-6}$, CC-GWAS with Exact threshold $p < 10^{-7}$, and CC-GWAS with Exact threshold $p < 5 * 10^{-8}$.

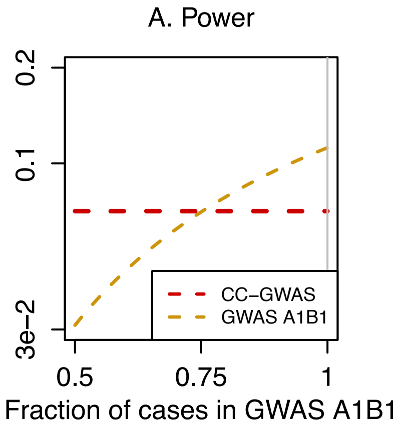

**Figure S8. Power of CC-GWAS and a direct case-case comparison (GWAS A1B1).** Analogue to main Figure 2 and based on the same default parameter values. We report the power to detect SNPs with effect sizes following a bivariate normal distribution, for each of two methods: CC-GWAS, and a direct case-case comparison GWAS A1B1. We vary the fraction of cases from the case-control comparison included in GWAS A1B1 (at fraction 1 all cases are included in GWAS A1B1; note that all cases are included in CC-GWAS throughout as CC-GWAS is based on the case-control GWAS results).

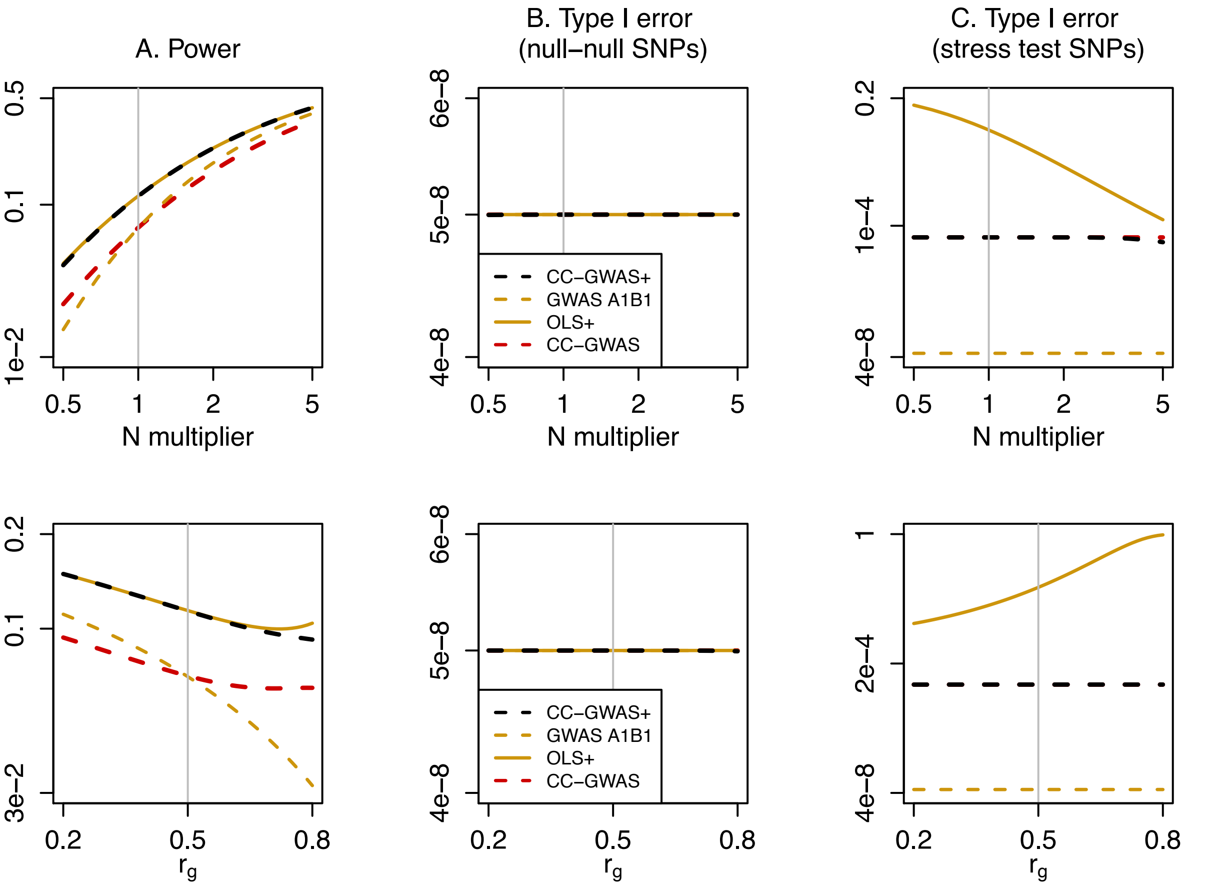

**Figure S9. Power and type I error of CC-GWAS+.** In CC-GWAS+, the the CC-GWAS$_{OLS}$ component is extended to the CC-GWAS+$_{OLS}$ component (see Methods) by including a direct case-case comparison (GWAS A1B1) resulting in weights of 0.35 for A1A0, -0.15 for B1B0 and 0.55 for A1B1 (for the same default parameter settings as in Figure 2, and assuming 75% of cases from the case-control comparisons were included in GWAS A1B1). The CC-GWAS$_{Exact}$ component is replaced with the CC-GWAS+$_{Exact}$ component (the GWAS A1B1 results, i.e. weights of 0, 0, and 1). CC-GWAS+ reports a SNP as statistically significant if it achieves $P < 5 * 10^{-8}$ using CC-GWAS+$_{OLS}$ weights *and* $P < 10^{-4}$ using CC-GWAS+$_{Exact}$ weights. We report (A) the power to detect SNPs with effect sizes following a bivariate normal distribution, (B) the type I error rate for loci with no effect on A1A0 or B1B0 (“null-null” SNPs) and (C) the type I error rate for SNPs with the same allele frequency in A1 vs. B1 that explain 0.10% of variance in A1 vs. A0 and 0.29% of variance in B1 vs. B0 (“stress test” SNPs), for each of four methods: CC-GWAS+, the direct case-case comparison GWAS A1B1, the CC-GWAS+$_{OLS}$ component, and CC-GWAS. Note that the CC-GWAS lines correspond exactly with Figure 2, because the same default parameters were used. For Panel B and Panel C the CC-GWAS+ lines (black dashed) overlay the CC-GWAS lines (red dashed).

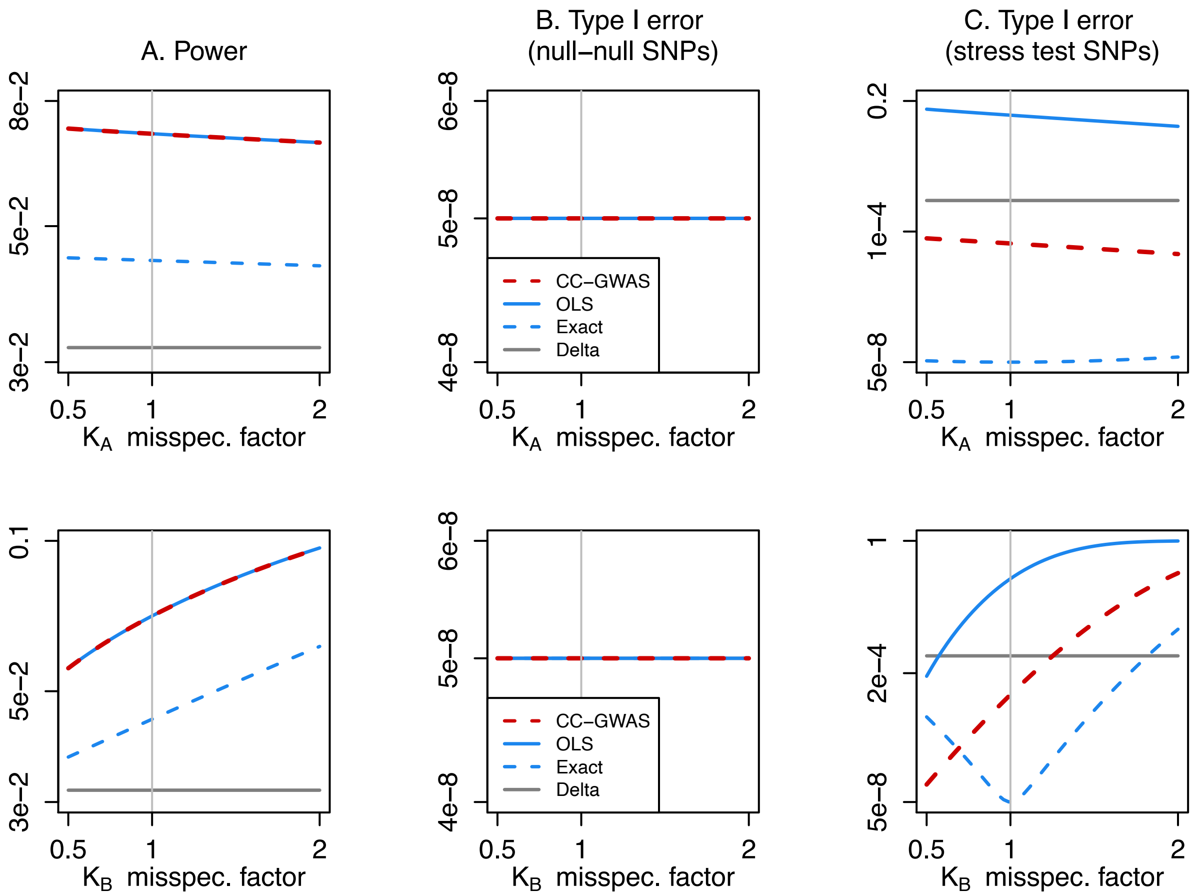

**Figure S10. Power and type I error of CC-GWAS when misspecifying the disorder prevalence.** This Figure is analogue to main Figure 2 and based on the same parameter values, with misspecification of the disorder prevalence in disorder A ($K_A$) and disorder B ($K_B$). We report (A) the power to detect SNPs with effect sizes following a bivariate normal distribution, (B) the type I error rate for loci with no effect on A1A0 or B1B0 ("null-null" SNPs) and (C) the type I error rate for SNPs with the same allele frequency in A1 vs. B1 that explain 0.10% of variance in A1 vs. A0 and 0.29% of variance in B1 vs. B0 ("stress test" SNPs), for each of four methods: CC-GWAS, the CC-GWAS$_{OLS}$ component, the CC-GWAS$_{Exact}$ component, and a naïve Delta method (see main text). The power of CC-GWAS either decreases or increases, the type I error rate at stress test SNPs respectively decreases or increases (i.e. same direction as change in power), and the type I error at null-null SNPs does not change.

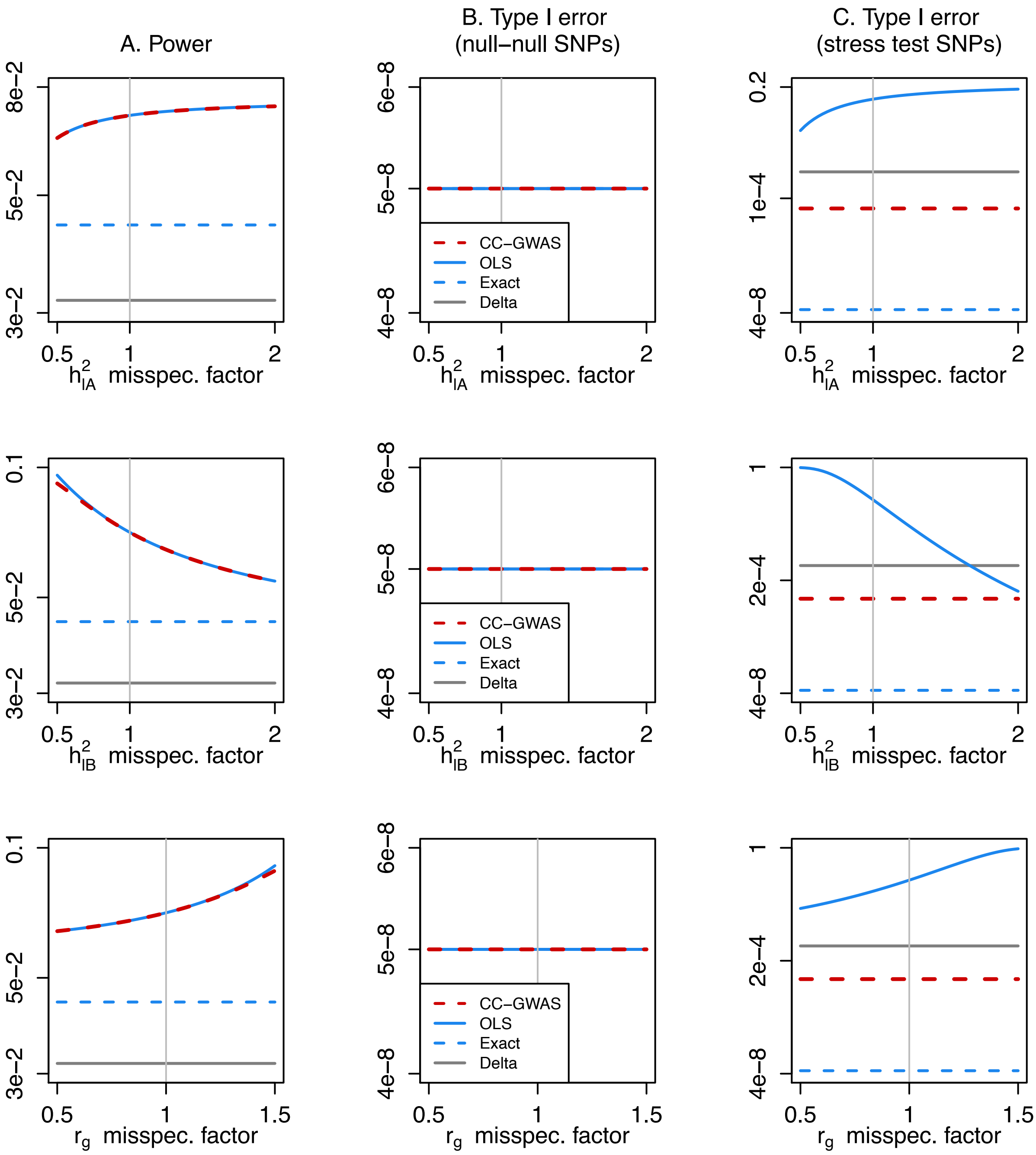

**Figure S11. Power and type I error of CC-GWAS when misspecifying the heritability or genetic correlation.** This Figure is analogue to main Figure 2 and based on the same parameter values, with misspecification of the heritability ($h^2_l$) and genetic correlation ($r_g$). We report (A) the power to detect SNPs with effect sizes following a bivariate normal distribution, (B) the type I error rate for loci with no effect on A1A0 or B1B0 ("null-null" SNPs) and (C) the type I error rate for SNPs with the same allele frequency in A1 vs. B1 that explain 0.10% of variance in A1 vs. A0 and 0.29% of variance in B1 vs. B0 ("stress test" SNPs), for each of four methods: CC-GWAS, the CC-GWAS$_{OLS}$ component, the CC-GWAS$_{Exact}$ component, and a naïve Delta method (see main text). Misspecification has little impact on power (defined as the total number of loci detected; it may also impact the number of CC-GWAS-specific loci), and no impact on type I error of CC-GWAS at null-null SNPs at stress test SNPs.

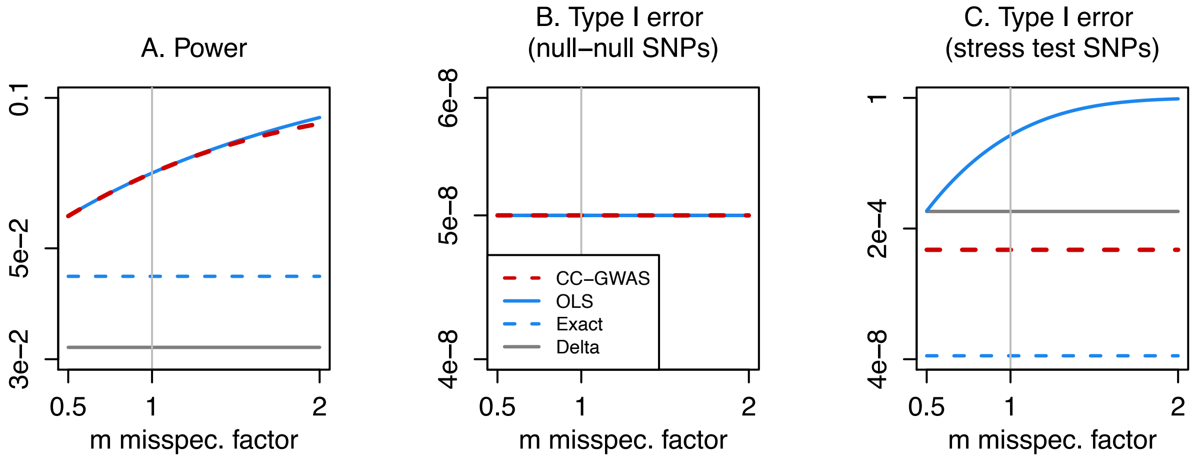

**Figure S12. Power and type I error of CC-GWAS when misspecifying the assumed number of causal SNPs.** This Figure is analogue to main Figure 2 and based on the same parameter values, with misspecification of the assumed number of causal SNPs. We report (A) the power to detect SNPs with effect sizes following a bivariate normal distribution, (B) the type I error rate for loci with no effect on A1A0 or B1B0 ("null-null" SNPs) and (C) the type I error rate for SNPs with the same allele frequency in A1 vs. B1 that explain 0.10% of variance in A1 vs. A0 and 0.29% of variance in B1 vs. B0 ("stress test" SNPs), for each of four methods: CC-GWAS, the CC-GWAS$_{OLS}$ component, the CC-GWAS$_{Exact}$ component, and a naïve Delta method (see main text). Misspecification has modest impact on power (defined as the total number of loci detected; it may also impact the number of CC-GWAS-specific loci), and no impact on type I error of CC-GWAS at null-null SNPs at stress test SNPs.

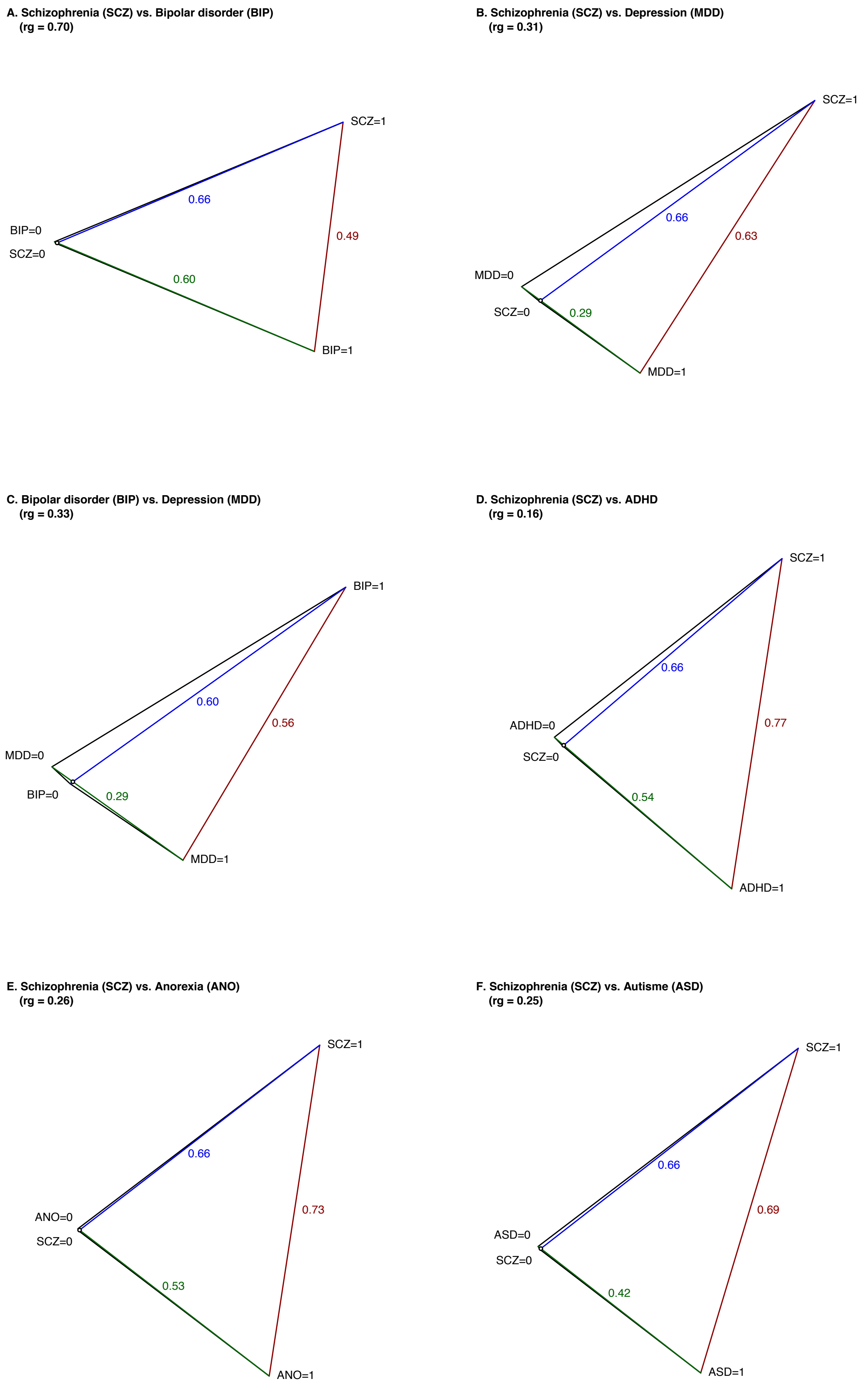

**Figure S13. Genetic distance between cases and/or controls from 31 disorder pairs.**

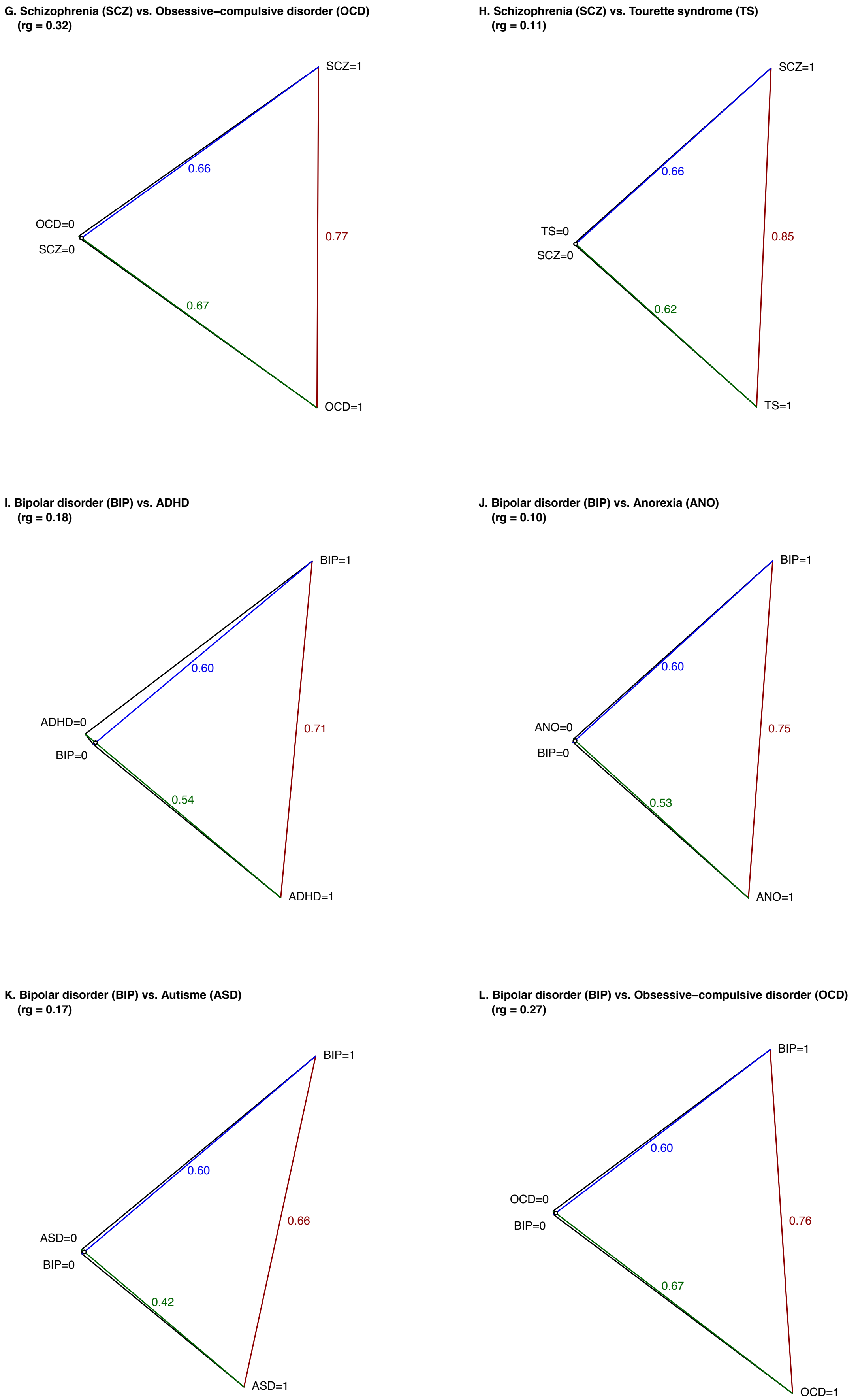

**Figure S13 continued.**

**M. Bipolar disorder (BIP) vs. Tourette syndrome (TS)**
**(rg = 0.08)**

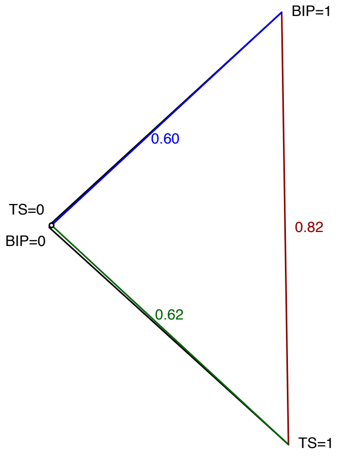

**N. Depression (MDD) vs. ADHD**
**(rg = 0.44)**

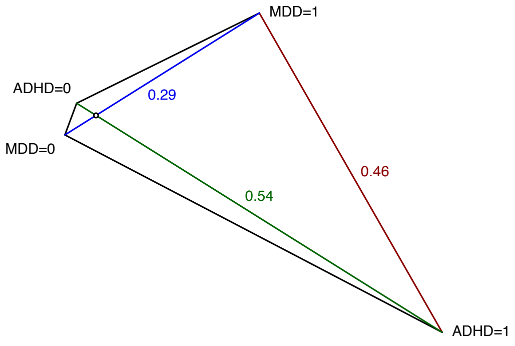

**O. Depression (MDD) vs. Anorexia (ANO)**
**(rg = 0.28)**

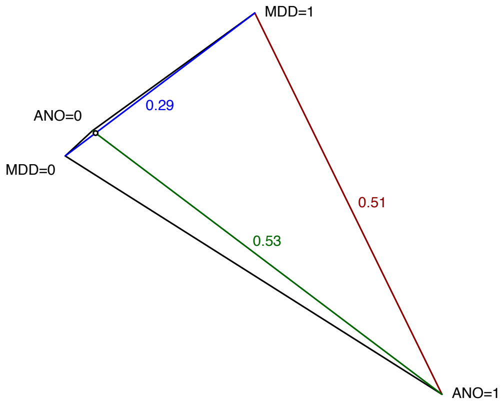

**P. Depression (MDD) vs. Autisme (ASD)**
**(rg = 0.34)**

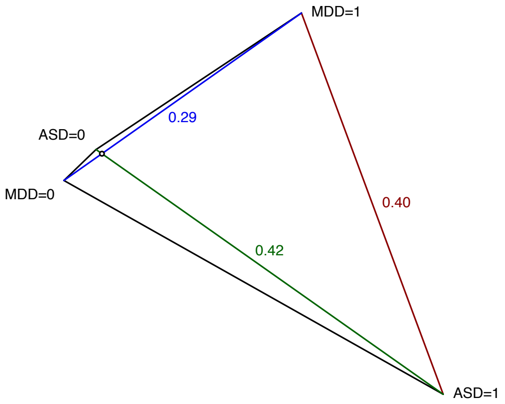

**Q. Depression (MDD) vs. Obsessive–compulsive disorder (OCD)**
**(rg = 0.25)**

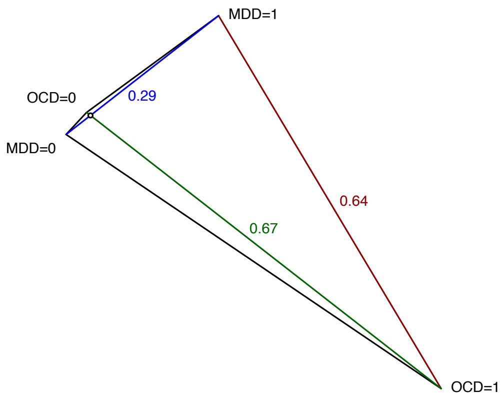

**R. Depression (MDD) vs. Tourette syndrome (TS)**
**(rg = 0.23)**

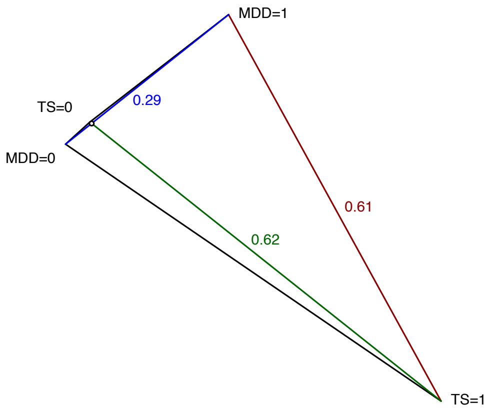

**Figure S13 continued.**

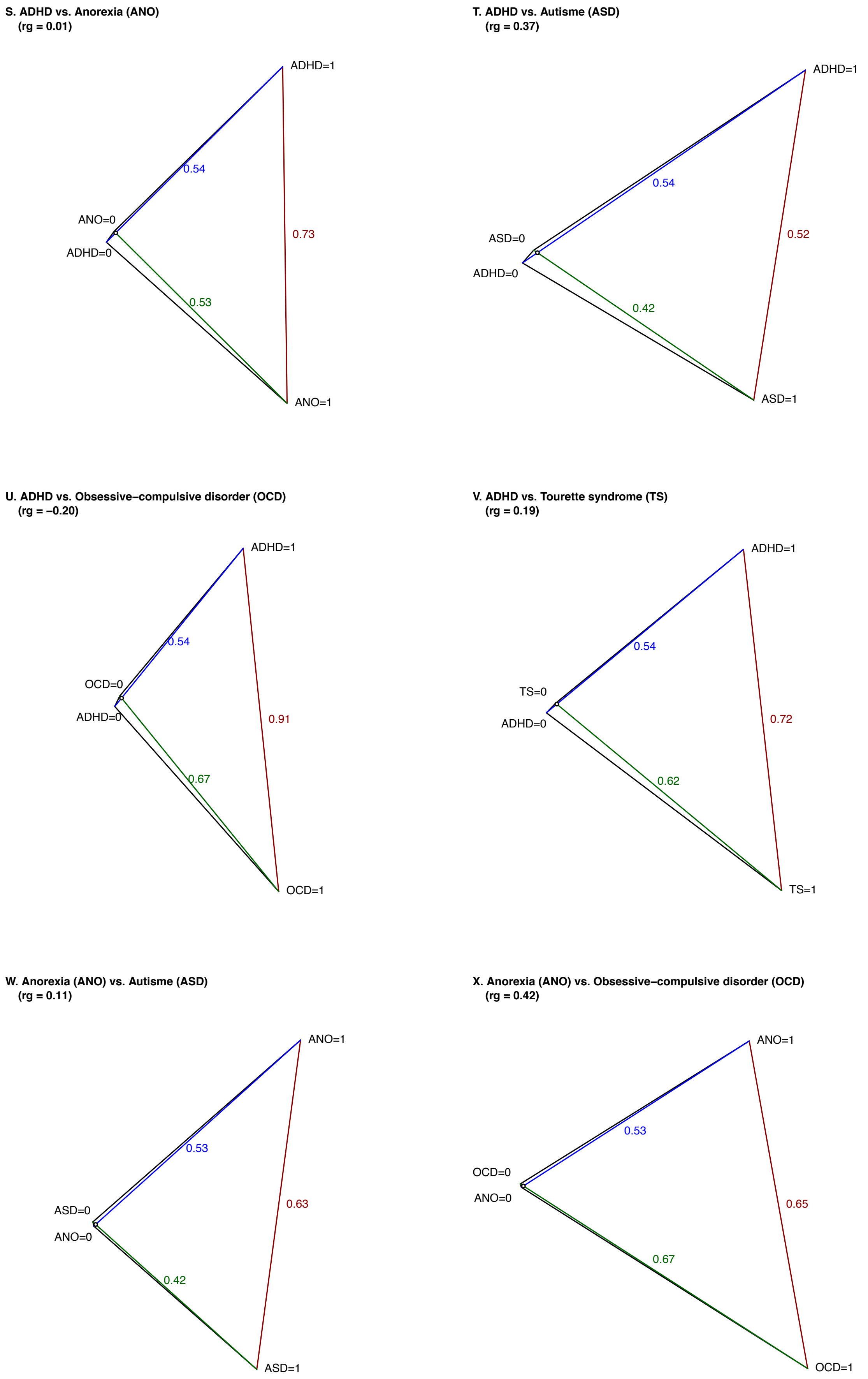

**Figure S13 continued.**

**Y. Anorexia (ANO) vs. Tourette syndrome (TS)**
**(rg = 0.08)**

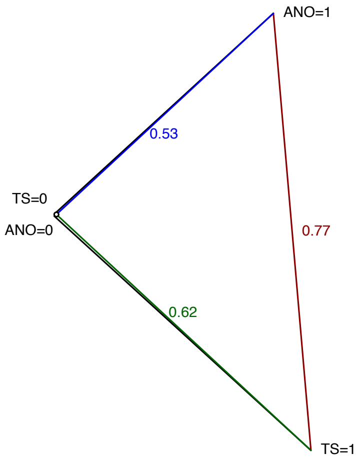

**Z. Autisme (ASD) vs. Obsessive−compulsive disorder (OCD)**
**(rg = 0.10)**

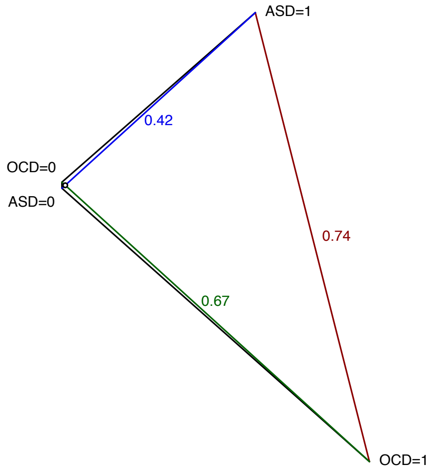

**AA. Autisme (ASD) vs. Tourette syndrome (TS)**
**(rg = 0.16)**

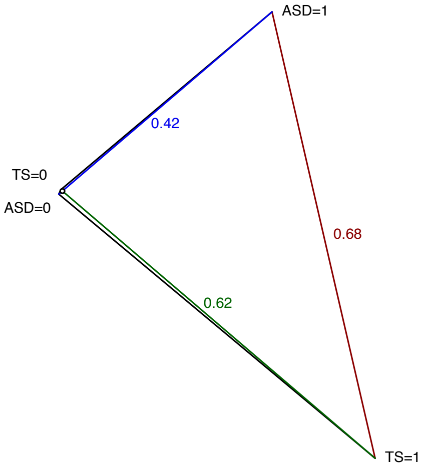

**AB. Obsessive−compulsive disorder (OCD) vs. Tourette syndrome (TS)**
**(rg = 0.50)**

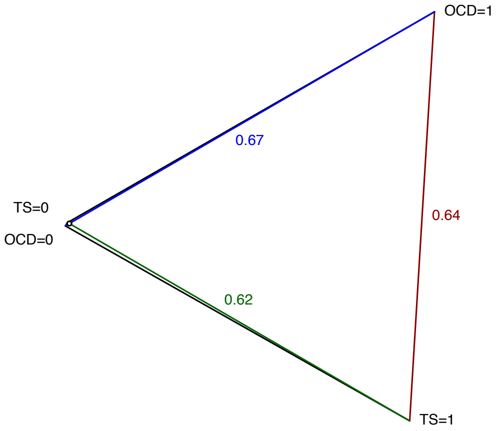

**AC. Crohn's disease (CD) vs. Ulcerative Colitis (UC)**
**(rg = 0.67)**

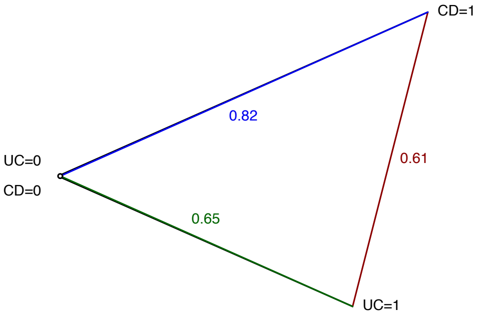

**AD. Crohn's disease (CD) vs. Rheumatoid Artritis (RA)**
**(rg = 0.09)**

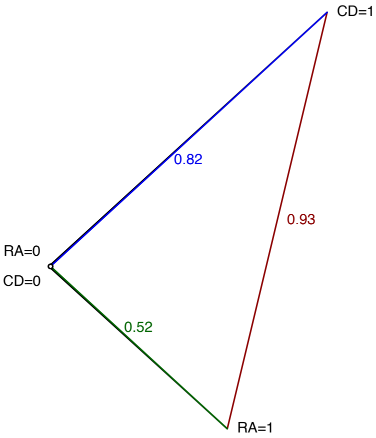

**Figure S13 continued.**

AE. Ulcerative Colitis (UC) vs. Rheumatoid Artritis (RA)
(rg = 0.04)

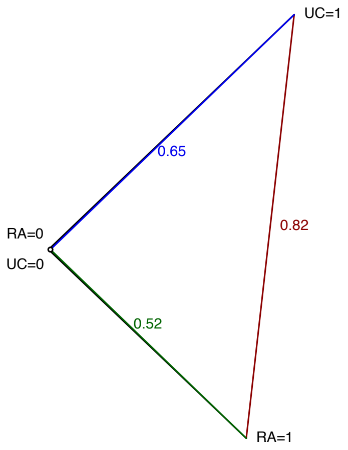

**Figure S13 continued. Genetic distance between cases and/or controls from 31 disorder pairs.** We report genetic distances for the 28 pairs of eight psychiatric disorders and 3 pairs of three autoimmune disorders. Genetic distances are displayed as $\sqrt{m * F_{ST,causal}}$, where $m$ is the number of independent causal variants and the square root facilitates 2-dimensional visualization. The quantity $m * F_{ST,causal}$ is derived based on the respective population prevalences, SNP-based heritabilities and genetic correlations (reported in Table 1 and Table S14). The cosine of the angle between the lines A1-A0 and B1-B0 is equal to the genetic correlation between disorder A and disorder B (see Methods). We note that $m * F_{ST,causal}$ is independent of $m$ when other parameters are fixed, because the equation for $F_{ST,causal}$ has $m$ in the denominator (see Methods). Numerical results are reported in Table S11. SCZ, schizophrenia; BIP, bipolar disorder; MDD, major depressive disorder; ADHD, attention deficit/hyperactivity disorder; ANO, anorexia nervosa; ASD, autism spectrum disorder; OCD, obsessive–compulsive disorder; TS, Tourette's Syndrome and Other Tic Disorders; CD, Crohn's disorder; UC, ulcerative colitis; RA, rheumatoid arthritis.

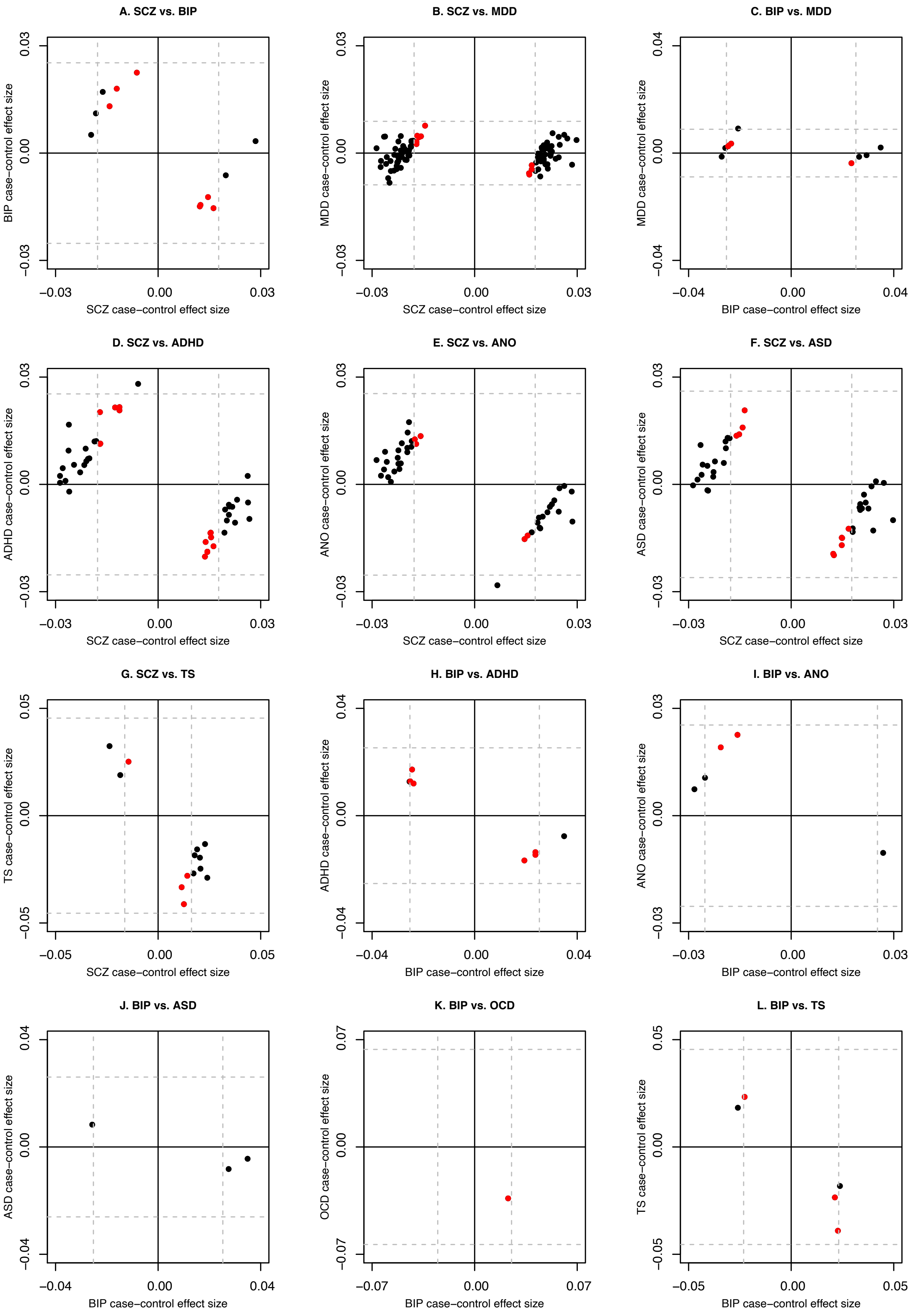

**Figure S14. Case-control effect sizes for CC-GWAS loci for 27 disorder pairs.**

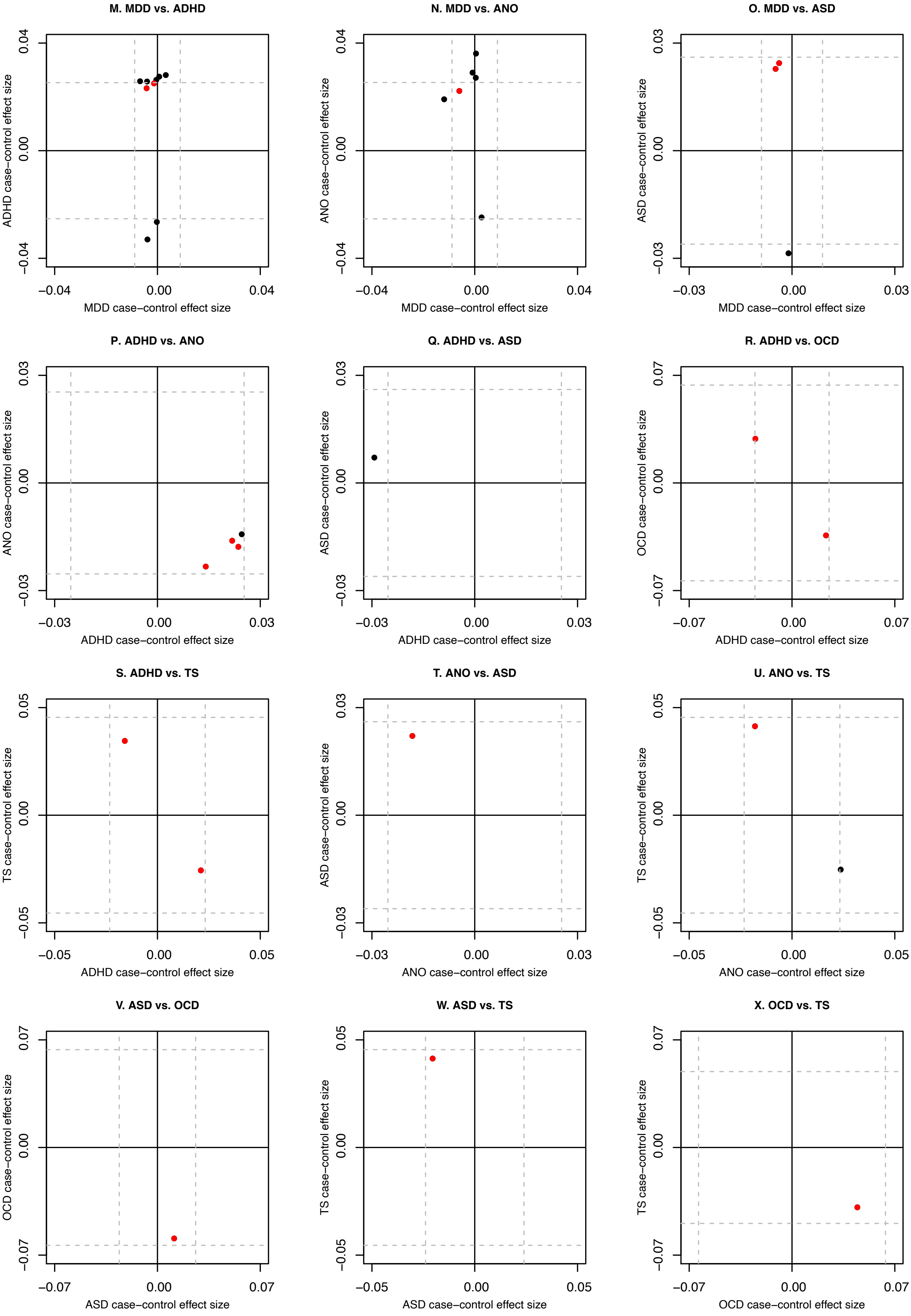

**Figure S14 continued.**

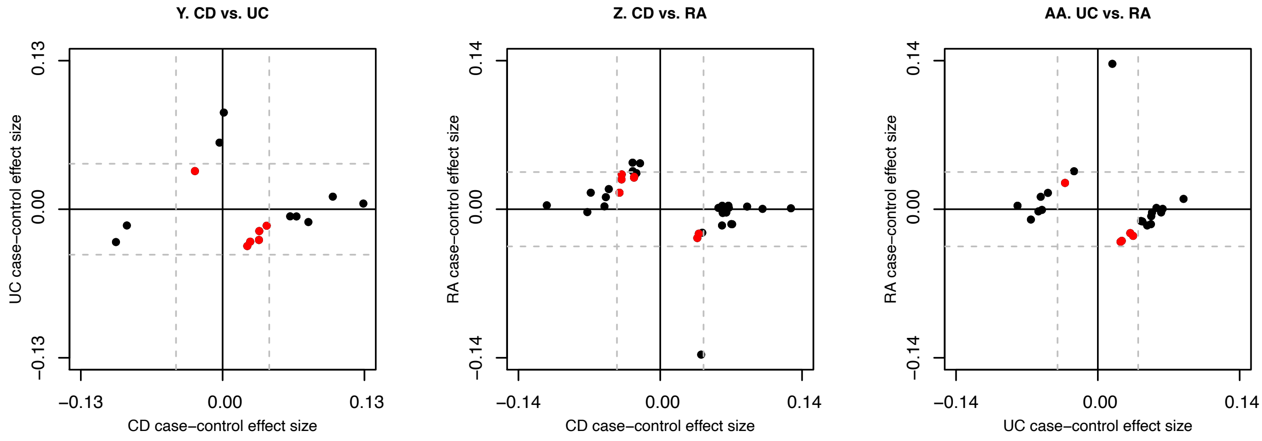

**Figure S14 continued. Case-control effect sizes for CC-GWAS loci for 27 disorder pairs.** This Figure is analogue to main Figure 4. We report the respective case-control effect sizes for lead SNPs at CC-GWAS loci for 27 disorder pairs of the eight psychiatric disorders and three autoimmune disorders. (Note that no CC-GWAS loci were found for 4 of 31 disorder pairs; Table 4). Effect sizes are reported on the standardized observed scale based on 50/50 case-control ascertainment. Red points denote CC-GWAS-specific loci, and black points denote remaining loci. Dashed lines denote effect-size thresholds for genome-wide significance. All red points (denoting lead SNPs for CC-GWAS-specific loci) lie inside the dashed lines for both disorders. SCZ, schizophrenia; BIP, bipolar disorder; MDD, major depressive disorder; ADHD, attention deficit/hyperactivity disorder; ANO, anorexia nervosa; ASD, autism spectrum disorder; OCD, obsessive–compulsive disorder; TS, Tourette's Syndrome and Other Tic Disorders; CD, Crohn's disorder; UC, ulcerative colitis; RA, rheumatoid arthritis.

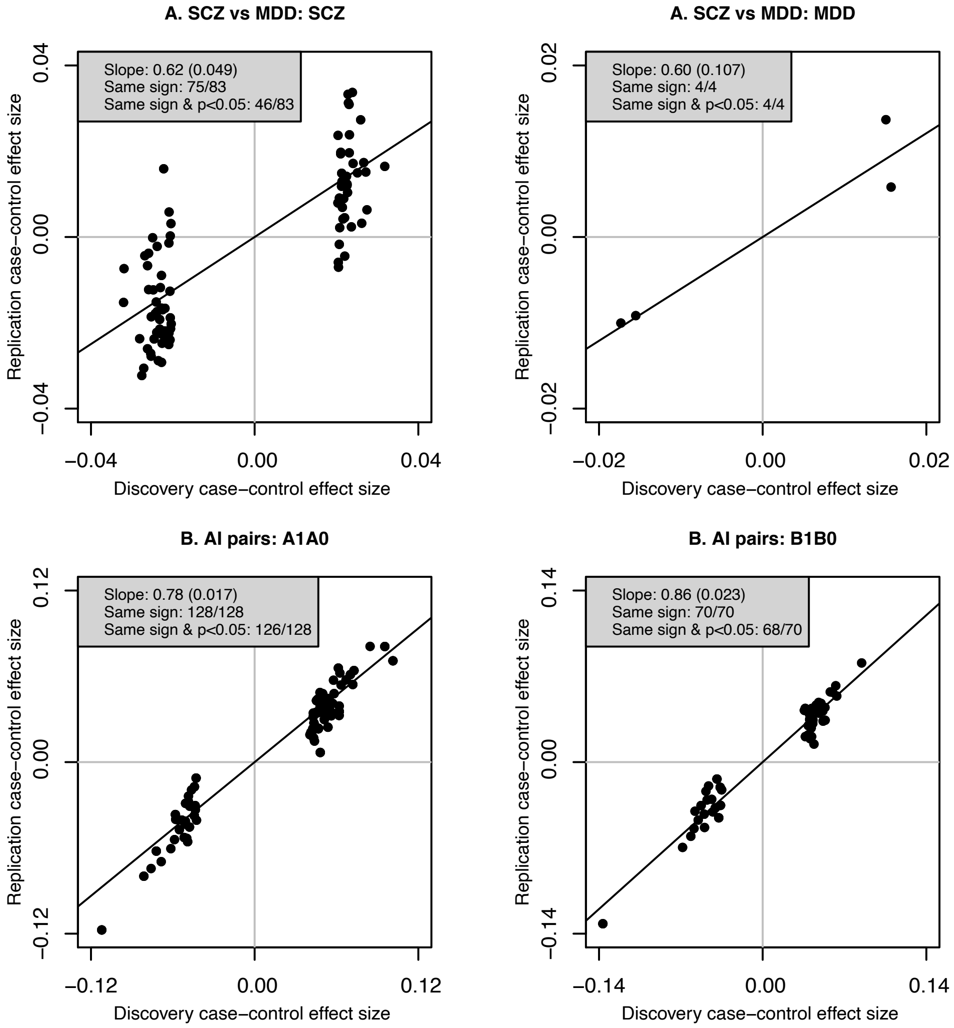

**Figure S15. Independent replication of within-disorder case-control GWAS results.** This Figure is analogue to main Figure 5, and shows within disorder replication of significant case-control loci. We report replication case-control effect sizes vs. discovery case-control effect sizes for schizophrenia (SCZ), major depressive disorder (MDD), disorder A of the three autoimmune disorders comparison, and disorder B of the three autoimmune disorders. We also report regression slopes (SE in parentheses), effect sign concordance, and effect sign concordance together with replication $P_{OLS}$<0.05. Numerical results are reported in Table S27. Note that the significant case-control loci are plotted here, whereas in Figure 5 the significant CC-GWAS loci are plotted; the number of loci reported in this Figure and in Figure 5 differ accordingly.

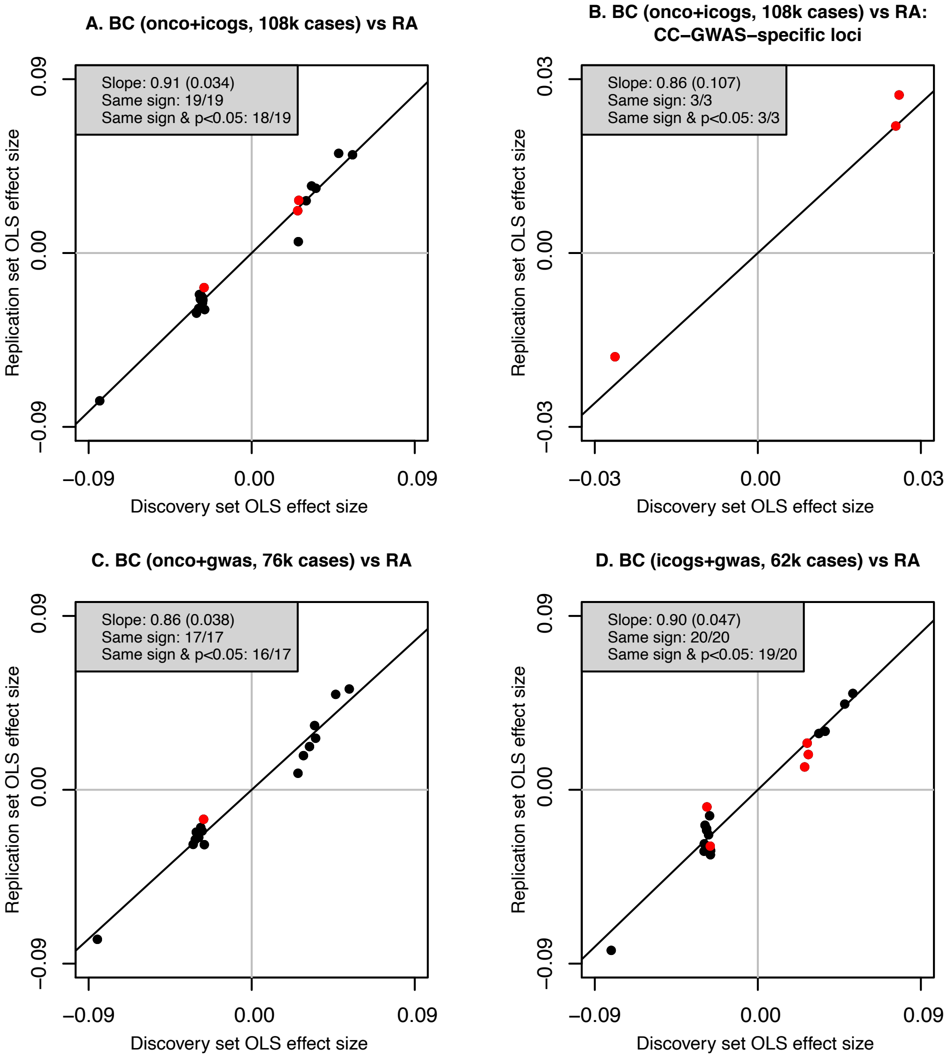

**Figure S16. Independent replication of breast cancer vs. rheumatoid arthritis CC-GWAS results.** We analysed breast cancer (BC)[3] and rheumatoid arthritis (RA)[27], as these were disorders with two sets of independent, publicly available GWAS summary statistics in large sample size (i.e. well-powered for replication analyses) without requiring the use of MetaSubtract[28] as in our main replication analyses. For RA, for all presented results, we used 8,875 cases and 29,367 controls (gwas sample in ref.[27]) for discovery analyses, and 5,486 cases and 14,556 controls (Immunochip sample in ref.[27]) for replication analyses. For BC, first, we used 108,067 cases and 88,386 controls (meta-analyses of the OncoArray sample and iCOGs sample in ref.[3]) for discovery analyses, and 14,910 cases and 17,588 controls (gwas sample in ref.[3]) for replication analyses (results in Panel A for all loci, and in Panel B for CC-GWAS-specific loci only). For BC, second, we used 76,192 cases and 63,082 controls (meta-analyses of the OncoArray sample and gwas sample in ref.[3]) for discovery analyses, and 46,785 cases and 42,892 controls (iCOGs sample in ref.[3]) for replication analyses (results in Panel C for all loci). For BC, third, we used 61,695 cases and 60,480 controls (meta-analyses of the iCOGs sample and gwas sample in ref.[3]) for discovery analyses, and 61,282 cases and 45,494 controls (OncoArray sample in ref.[3]) for replication analyses (results in Panel D for all loci). These results consistently confirm that CC-GWAS results replicate well in independent data, for all loci as well (Panels A, C and D) as for CC-GWAS-specific loci (although there were only few loci to evaluate; Panel B).

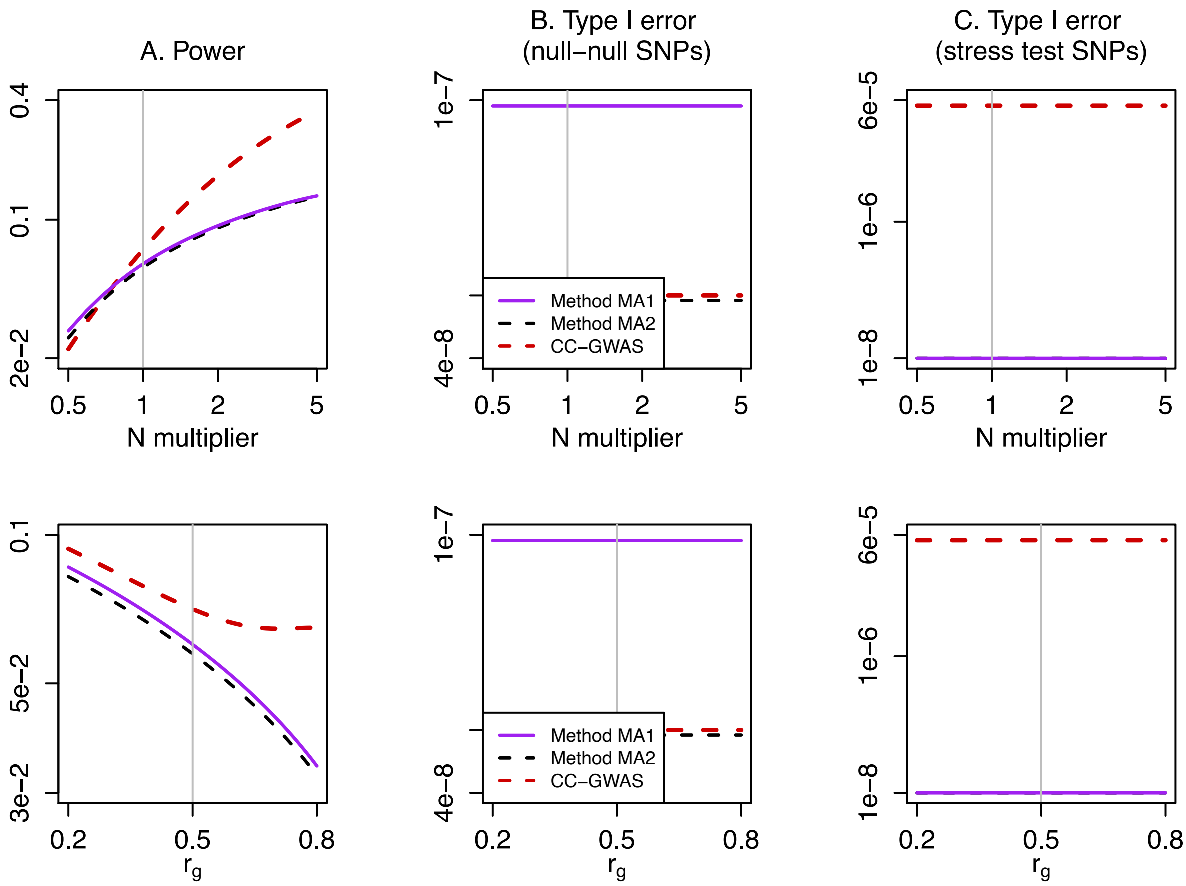

**Figure S17. Power and type I error of simple meta-analysis based method and CC-GWAS.** This Figure is analogue to main Figure 2 and based on the same parameter values, and compares the meta-analysis method (Method MA) to CC-GWAS. We assessed two different versions of Method MA: Method MA1, which outputs SNPs with $p<5x10^{-8}$ for one or both disorders and meta-analysis of both disorder GWAS $p\geq5x10^{-8}$; and Method MA2, which outputs SNPs with $p<2.5x10^{-8}$ for one or both disorders and meta-analysis of both disorder GWAS $p\geq2.5x10^{-8}$. We report (A) the power to detect SNPs with effect sizes following a bivariate normal distribution, (B) the type I error rate for loci with no effect on A1A0 or B1B0 ("null-null" SNPs) and (C) the type I error rate for SNPs with the same allele frequency in A1 vs. B1 that explain 0.10% of variance in A1 vs. A0 and 0.29% of variance in B1 vs. B0 ("stress test" SNPs). We reach three conclusions. First, Method MA1 and Method MA2 yield less power than CC-GWAS. Second, at null-null SNPs, Methods MA1 has an increased type I error rate ($>5x10^{-8}$). When aiming to apply a meta-analysis method, we thus advise to apply Method MA2. Three, both Method MA1 and Method MA2 have more stringent control of type I error at stress test SNPs than CC-GWAS, but we believe that the type I error control at stress test SNPs of CC-GWAS is adequate as stress test SNPs cannot be numerous (see Discussion in main text) and a type I error control at stress test SNPs of $<5x10^{-8}$ may be overly conservative.

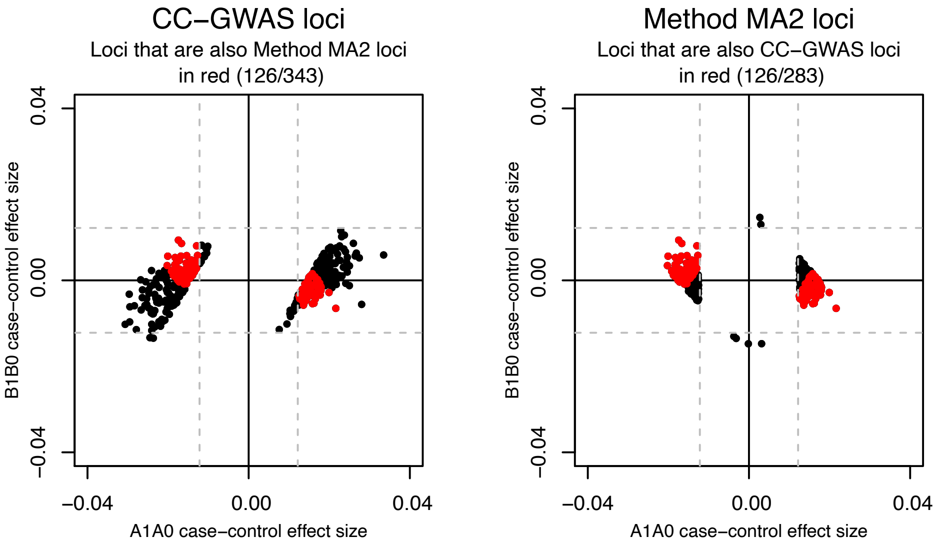

**Figure S18. Loci detected in simulations of simple meta-analysis based method and CC-GWAS.** We compared loci detected by CC-GWAS with loci detected by the meta-analysis method 2 (Method MA2, which outputs SNPs with p<2.5x10$^{-8}$ for one or both disorders and meta-analysis of both disorder GWAS p≥2.5x10$^{-8}$). See Figure S17 for comparison of power, type I error at null-null SNPs and type I error at stress test SNPs. Here we report true positive loci from simulation of case-control GWAS results in line with the parameter values in main Figure 2. The dashed grey lines mark the respective case-control significance thresholds. CC-GWAS detected more loci (343) that Method MA2 (283), and 126 loci were detected by both (points labelled in red in both Panels). The 217 loci detected by CC-GWAS and not by Method MA2 (points labelled in black in left Panel) had relatively large case-control effects for one of both disorders (yielding meta-analysis of both disorder p<2.5x10$^{-8}$, thus not being detected by Method MA2). The 157 loci detected by Method MA2 and not by CC-GWAS (points labelled in black in right Panel) had relatively small case-control effect-sizes only just significant in one both disorder and often non-significant but with same direction in the other disorder (resulting in a non-significant weighted difference of effect-sizes, thus not being detected by CC-GWAS). Note that method MA1 and method MA2 both fail to detect CC-GWAS-specific loci, which are of particularly high interest.

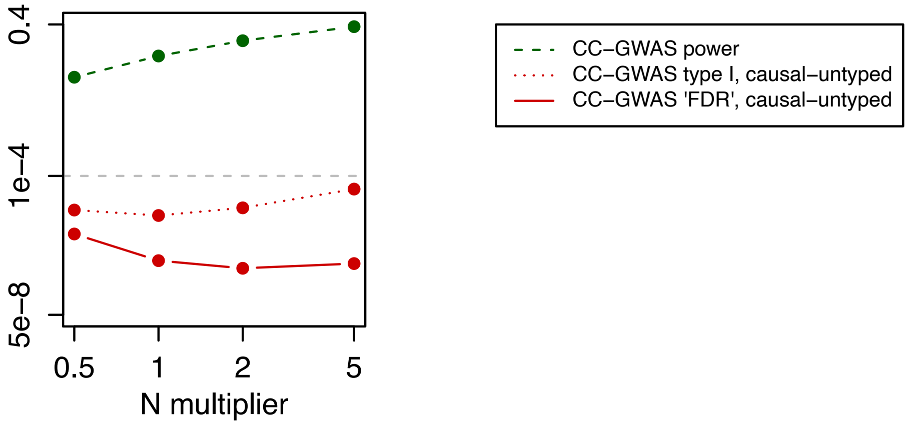

**Figure S19. Type I error and FDR due to differential tagging of a causal stress test SNP.** This Figure reports a rough approximation of the false discovery rate (FDR) due to differential tagging of a causal stress test SNP, based on the parameter values from main Figure 2A and main Figure 3B. We combine information from main Figure 2A (the green dashed line reporting the power of CC-GWAS for true positive association) and main Figure 3B (the red dotted line reporting the per locus type I error rate due to differential tagging of a causal stress test SNP that is untyped). We assume a total of 15 untyped stress test SNPs and loci in line with Figure 3B (explaining almost half the heritability in disorder B: 15 x 0.29% =4.35% = ~0.5 x $h^2_{l,B}$), and 2,500 causal SNPs with different allele frequencies among cases of disorders A and B and effect-sizes following the bivariate normal distribution from main Figure 2A. The roughly approximated FDR (red solid line) is thus reported as {15 times the type I error (red dotted line)} divided by {2,500 times the power (green dashed line) plus 15 times the type I error (red dotted line)}. This suggests that the FDR does not (or much less) increase with increasing sample-size than the type I error rate due to differential tagging of causal stress test SNPs that is untyped, even in the extreme scenario where 50% of the heritability in disorder B would be explained by strong-effect untyped stress test SNPs. We emphasize this plot provides only a rough approximation and results may differ based on different assumption and other parameter values.

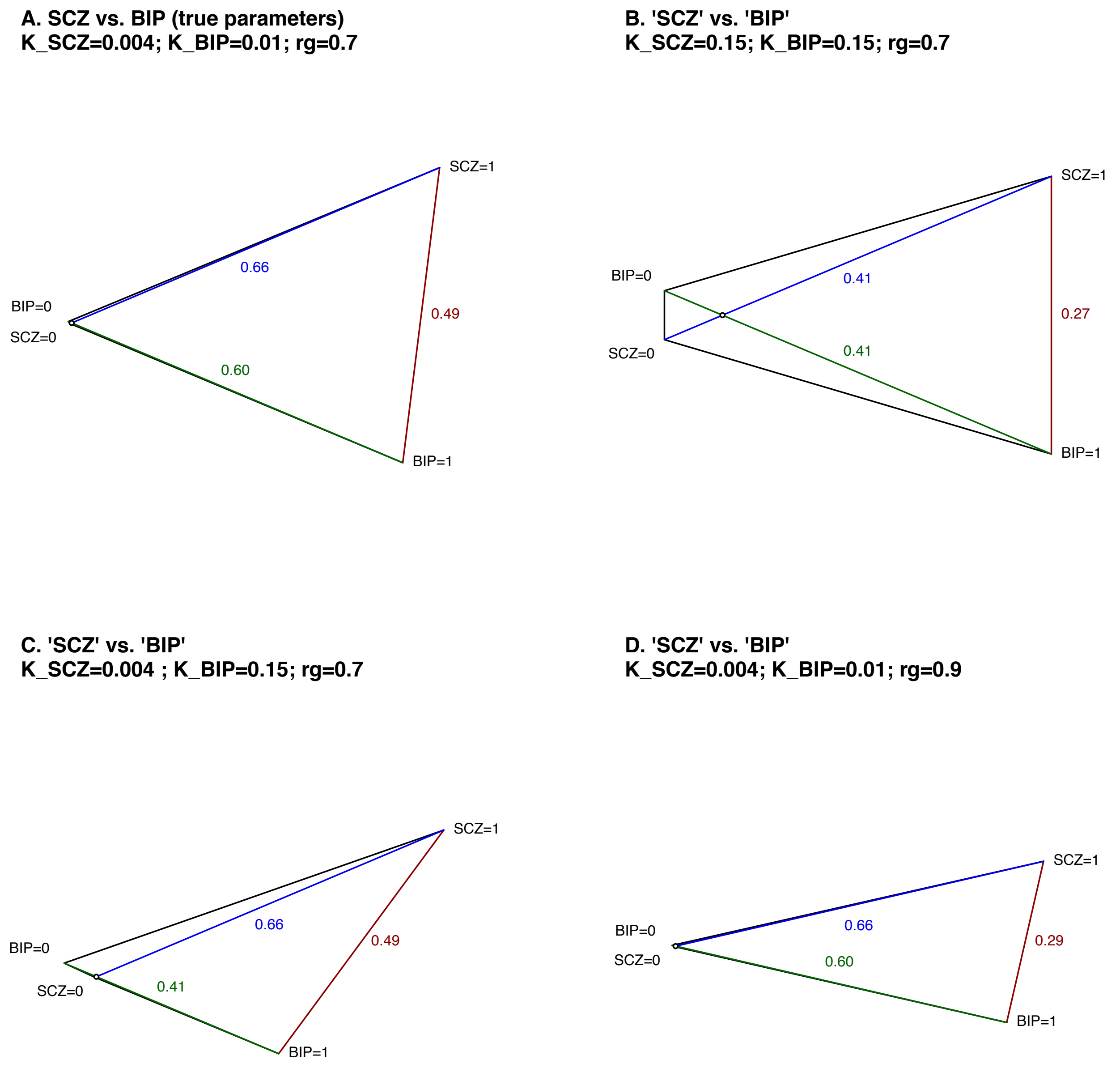

**Figure S20. $F_{ST,causal}$ between SCZ cases and BIP cases when varying the disorder prevalences and genetic correlation.** This Figure corresponds to main Figure 1. Genetic distances are displayed as $\sqrt{m * F_{ST,causal}}$, derived based on the respective population prevalences, SNP-based heritabilities and genetic correlations. We explore why, despite the large genetic correlation ($r_g$ = 0.7), the genetic distance between SCZ cases and BIP cases is only slightly smaller ($\sqrt{m * F_{ST,causal}} = 0.49$) than the case-control distances in SCZ (0.66) and in BIP (0.60) (see Panel A, equal to main Figure 1B). First, it can be seen that when the ascertainment (due to low disorder prevalences) would have been less profound for both SCZ and BIP, the distance between cases would have been considerably smaller (0.27; Panel B; note that the case-control distances are also smaller due to less ascertainment, but still 0.49/0.6 (Panel A) >> 0.27/0.41 (Panel B)). Second, when the ascertainment would have been less profound for only one disorder, the distance between cases would not have been smaller (Panel C). Third, when the genetic correlation would have been larger, the distance between cases would have been considerable smaller (0.29; Panel D). We conclude that the genetic distance between SCZ cases and BIP cases is only slightly than the respective case-control distances, because of (i) the doubly strong ascertainment in SCZ cases and BIP cases and (ii) because a genetic correlation of 0.7 is still considerable smaller than a genetic correlation of 1.

### References

1. Turley, P. *et al.* Multi-trait analysis of genome-wide association summary statistics using MTAG. *Nat. Genet.* **50**, 229–237 (2018).
2. Frei, O. *et al.* Bivariate causal mixture model quantifies polygenic overlap between complex traits beyond genetic correlation. *Nat. Commun.* **10**, 2417 (2019).
3. Michailidou, K. *et al.* Association analysis identifies 65 new breast cancer risk loci. *Nature* **551**, 92–94 (2017).
4. Sullivan, P. F. & Geschwind, D. H. Defining the Genetic, Genomic, Cellular, and Diagnostic Architectures of Psychiatric Disorders. *Cell* **177**, 162–183 (2019).
5. Finucane, H. K. *et al.* Partitioning heritability by functional annotation using genome-wide association summary statistics. *Nat. Genet.* (2015).
6. Gazal, S. *et al.* Linkage disequilibrium–dependent architecture of human complex traits shows action of negative selection. *Nat. Genet.* **49**, 1421–1427 (2017).
7. Gazal, S., Marquez-Luna, C., Finucane, H. K. & Price, A. L. Reconciling S-LDSC and LDAK functional enrichment estimates. *Nat. Genet.* **51**, 1202–1204 (2019).
8. Bulik-Sullivan, B. *et al.* An atlas of genetic correlations across human diseases and traits. *Nat. Genet.* (2015).
9. Byrne, E. M. *et al.* Conditional GWAS analysis identifies putative disorder-specific SNPs for psychiatric disorders. *bioRxiv* 592899 (2019). doi:10.1101/592899
10. Lee, P. H. *et al.* Genomic Relationships, Novel Loci, and Pleiotropic Mechanisms across Eight Psychiatric Disorders. *Cell* **179**, 1469-1482.e11 (2019).
11. Buniello, A. *et al.* The NHGRI-EBI GWAS Catalog of published genome-wide association studies, targeted arrays and summary statistics 2019. *Nucleic Acids Res.* **47**, D1005–D1012 (2019).
12. Pardiñas, A. F. *et al.* Common schizophrenia alleles are enriched in mutation-intolerant genes and in regions under strong background selection. *Nat. Genet.* **50**, 381–389 (2018).
13. Loh, P.-R., Kichaev, G., Gazal, S., Schoech, A. P. & Price, A. L. Mixed-model association for biobank-scale datasets. *Nat. Genet.* **50**, 906–908 (2018).
14. Bulik-Sullivan, B. K. *et al.* LD Score regression distinguishes confounding from polygenicity in genome-wide association studies. *Nat. Genet.* **47**, 291–5 (2015).
15. Stahl, E. A. *et al.* Genome-wide association study identifies 30 loci associated with bipolar disorder. *Nat. Genet.* **51**, 793–803 (2019).
16. Howard, D. M. *et al.* Genome-wide meta-analysis of depression identifies 102 independent variants and highlights the importance of the prefrontal brain regions. *Nat. Neurosci.* **22**, 343–352 (2019).
17. Ruderfer, D. M. *et al.* Genomic Dissection of Bipolar Disorder and Schizophrenia, Including 28 Subphenotypes. *Cell* **173**, 1705-1715.e16 (2018).
18. O'Connor, L. J. *et al.* Extreme Polygenicity of Complex Traits Is Explained by Negative Selection. *Am. J. Hum. Genet.* **105**, 456–476 (2019).
19. Price, A. L. *et al.* Discerning the ancestry of European Americans in genetic association studies. *PLoS Genet.* **4**, e236 (2008).
20. Bhatia, G., Patterson, N., Sankararaman, S. & Price, A. L. Estimating and interpreting FST: The impact of rare variants. *Genome Res.* **23**, 1514–1521 (2013).
21. Benjamini, Y. & Hochberg, Y. Controlling the False Discovery Rate: A Practical and Powerful Approach to Multiple Testing. *J. R. Stat. Soc. Ser. B* **57**, 289–300 (1995).
22. Lee, S. H., Wray, N. R., Goddard, M. E. & Visscher, P. M. Estimating Missing Heritability for Disease from Genome-wide Association Studies. *Am. J. Hum. Genet.* **88**, 294–305 (2011).
23. Golan, D., Lander, E. S. & Rosset, S. Measuring missing heritability: Inferring the contribution of common variants. *Proc. Natl. Acad. Sci. U. S. A.* **111**, E5272-81 (2014).
24. Lloyd-Jones, L. R., Robinson, M. R., Yang, J. & Visscher, P. M. Transformation of Summary

Statistics from Linear Mixed Model Association on All-or-None Traits to Odds Ratio. *Genetics* **208**, 1397–1408 (2018).
25. Reich, D. E. *et al.* Linkage disequilibrium in the human genome. *Nature* (2001). doi:10.1038/35075590
26. Slatkin, M. Linkage disequilibrium - Understanding the evolutionary past and mapping the medical future. *Nature Reviews Genetics* (2008).
27. Okada, Y. *et al.* Genetics of rheumatoid arthritis contributes to biology and drug discovery. *Nature* **506**, 376–81 (2014).
28. Nolte, I. M. *et al.* Missing heritability: is the gap closing? An analysis of 32 complex traits in the Lifelines Cohort Study. *Eur. J. Hum. Genet.* **25**, 877–885 (2017).
